# Common Dynamic Determinants Govern Quorum Quenching Activity in N-terminal Serine Hydrolases

**DOI:** 10.1101/2022.01.13.476167

**Authors:** Bartlomiej Surpeta, Michal Grulich, Andrea Palyzová, Helena Marešová, Jan Brezovsky

## Abstract

Growing concerns about microbial antibiotic resistance have motivated extensive research into ways of overcoming antibiotic resistance. Quorum quenching (QQ) processes disrupt bacterial communication via quorum sensing, which enables bacteria to sense the surrounding bacterial cell density and markedly affects their virulence. Due to its indirect mode of action, QQ is believed to exert limited pressure on essential bacterial functions and may thus avoid inducing resistance. Although many enzymes display QQ activity against various bacterial signaling molecules, their mechanisms of action are poorly understood, limiting their potential optimization as QQ agents. Here we evaluate the capacity of three N-terminal serine hydrolases to degrade N-acyl homoserine lactones that serve as signaling compounds for Gram-negative bacteria. Using molecular dynamics simulations of the free enzymes and their complexes with two signaling molecules of different lengths, followed by quantum mechanics/molecular mechanics molecular dynamics simulations of their initial catalytic steps, we clarify the molecular processes underpinning their QQ activity. We conclude that all three enzymes degrade bacterial signaling molecules via similar reaction mechanisms. Moreover, we experimentally confirmed the activity of two penicillin G acylases from *Escherichia coli* (ecPGA) and *Achromobacter spp*. (aPGA), adding these biotechnologically well-optimized enzymes to the QQ toolbox. We also observed enzyme- and substrate-dependent differences in the catalytic actions of these enzymes, arising primarily from the distinct structures of their acyl-binding cavities and the dynamics of their molecular gates. As a consequence, the first reaction step catalyzed by ecPGA with a longer substrate had an elevated energy barrier because its shallow acyl binding site could not accommodate a productive substrate-binding configuration. Conversely, aPGA in complex with both substrates exhibited unfavorable energetics in both reaction steps due to the dynamics of the residues gating the acyl binding cavity entrance. Finally, the energy barriers of the second reaction step catalyzed by *Pseudomonas aeruginosa* acyl-homoserine lactone acylase with both substrates were higher than in the other two enzymes due to the unique positioning of Arg297β in this enzyme. The discovery of these dynamic determinants will guide future efforts to design robust QQ agents capable of selectively controlling virulence in resistant bacterial species.

## INTRODUCTION

Efficient control of bacterial populations is critical for various aspects of our daily life. In recent decades, antibiotics have served as the primary tools for addressing this challenge in human and veterinary medicine, animal food production, agriculture, aquaculture, and anti-biofouling applications in various industries; in most of these fields, antibiotics continue to be heavily used.^1-6^ Unfortunately, the interference of antibiotics with the essential bacterial functions exerts strong selective pressure, leading to the development of antibiotic resistance via numerous distinct mechanisms.^6^ The development of resistance has been accelerated and exacerbated by the widespread misuse and overuse of antibiotics in almost all of their areas of application.^7–11^ In addition, recent studies have shown that resistance genes can be transferred between bacteria of the same or even different species, enabling the spread of resistance without direct exposure to antimicrobials.^12,13^ As a result, resistant bacteria have been detected in all studied environments including soil, sea, food products, drinking water, and even samples from Antarctica.^12,14–17^ The rapid spread of antibiotic resistance thus constitutes an alarming global crisis.^7,8,18,19^

To address this threat,^6,12,17,20^ a major global cooperative effort will be needed.^8^ The emergence of resistance towards currently used antibiotics can be delayed by providing extensive education about this problem to increase social awareness, improve diagnosis, ensure that antibiotics are prescribed only where appropriate, limit the use of antibiotics in agriculture and aquaculture, and prioritize the most effective antibiotics for medical use.^8,12,21–24^ The discovery and development of new generations of antimicrobials as alternatives to currently available conventional antibiotics would also provide a valuable way of managing the problem of bacterial resistance. Unfortunately, conventional methods for developing new drugs have not been particularly successful in this respect in recent decades.^25–28^ This is largely due to the existence of multiple bacterial mechanisms for managing the toxicity of active compounds including penetration barriers; systems that inactivate antibiotics by destroying them, modifying them, or excreting them via efflux pumps; and modification, switching, or sequestration of the biological systems targeted by the antibiotics.^26,29,30^ Therefore, alternative strategies that target non-essential bacterial functions and thus exert less selective pressure favoring the development of resistance are needed to complement or even replace antibiotics in some of their areas of application.^31–35^

Bacterial virulence factors are promising targets for strategies aiming to disarm rather than kill bacteria. One of the most extensively studied approaches of this type involves disrupting the quorum sensing (QS) process. QS enables communication between bacteria and results in population density-dependent collective behavior controlled by the concentration of specific signaling molecules.^36,37^ These molecules bind to transcriptional activators and stimulate the expression of genes responsible for regulating virulence factors, secondary metabolite production, biofilm formation, and other processes crucial for pathogenicity.^38^ The disruption of QS, widely known as quorum quenching (QQ), has been identified as a promising strategy for controlling biofouling,^39^ treating bacterial infections,^40^ and protecting crops and aquaculture systems.^36,41^

Various organic molecules have been reported to serve as signaling molecules in QS depending on the bacterial species. A particularly notable class are the *N*-acyl-homoserine lactones (HSLs)^41^ which are predominantly used by highly pathogenic Gram-negative bacteria including the critical priority genus *Pseudomonas*.^42^ For example, the QS signal in the globally prioritized *Pseudomonas aeruginosa* is carried by *N*-(3-oxo-dodecanoyl)-L-homoserine lactone. In this case and analogously in other species,^43^ QQ can be achieved by inhibiting either the synthesis of HSL (which is catalyzed by the protein LasI) or its detection by the protein LasR, which is a transcriptional activator of QS genes.^37,43^ Although quorum sensing inhibition (QSI) can effectively interfere with bacterial communication, resistance to QSI is possible through changes in the structure of its protein targets or via the action of efflux pumps, as was demonstrated for furanone C-30 in *P. aeruginosa*.^43–45^ It has been suggested that the direct inactivation of signaling compounds^37,46^ by the action of QQ enzymes might be less likely to exert pressure on bacteria because such inactivation processes would operate outside the cell in the external environment and so would not be subject to the development of resistance by mechanisms known for antibiotics, drugs, or QS inhibitors.^36^

To date, three enzyme classes capable of QQ of HSL-based communication have been discovered: (i) acylases (amidases), which cleave the amide bond between the homoserine lactone moiety and the acyl chain; (ii) lactonases, which break the lactone ring; and (iii) oxidoreductases, which modify the acyl chain.^36,37,41^ HSL degradation catalyzed by acylases is irreversible and forms neutral cleavage products that are readily metabolized. Furthermore, unlike the other two QQ enzyme types, acylases are often specific for a narrow range of signaling molecules, which enables selective targeting of pathogens while avoiding undesired side effects on beneficial microbes^39^ and the limited efficiency of broad-specificity enzymes in environments with multiple targets.^47^ Most QQ acylases belong to the N-terminal hydrolase superfamily and have common features including auto-proteolytic activation by cleavage of a linker peptide and activity based on an N-terminal serine, threonine or cysteine residue that serves as a nucleophile in the self-activation process and its subsequent catalytic mechanism. They are divided into four main subfamilies: aculeacin A acylase, penicillin G acylase, AmiE amidase, and penicillin V acylase.^37^

*Pseudomonas aeruginosa* acyl-homoserine lactone acylase (paPvdQ) from the aculeacin A acylase subfamily is the most developed acylase in terms of practical QQ applications. It preferentially cleaves long HSLs (∼12 carbon atoms) because of its deep acyl-binding cavity (**Figure 1A)**.^48^ By modifying this cavity, Koch *et al*. successfully altered the specificity of paPvdQ towards C08-HSL, a signaling molecule used by *Burkholderia cenocepacia*, enabling host survival in a *Galleria mellonella* infection model.^49^ Moreover, a dry powder formulation of this protein was developed to treat pulmonary *Pseudomonas aeruginosa* infections by inhalation^50^ and was recently shown to be effective *in vivo* in a mouse model.^40^ While these results make paPvdQ a promising QQ agent, its molecular mechanism of action is not fully understood, restricting its further development.

**Figure 1.**
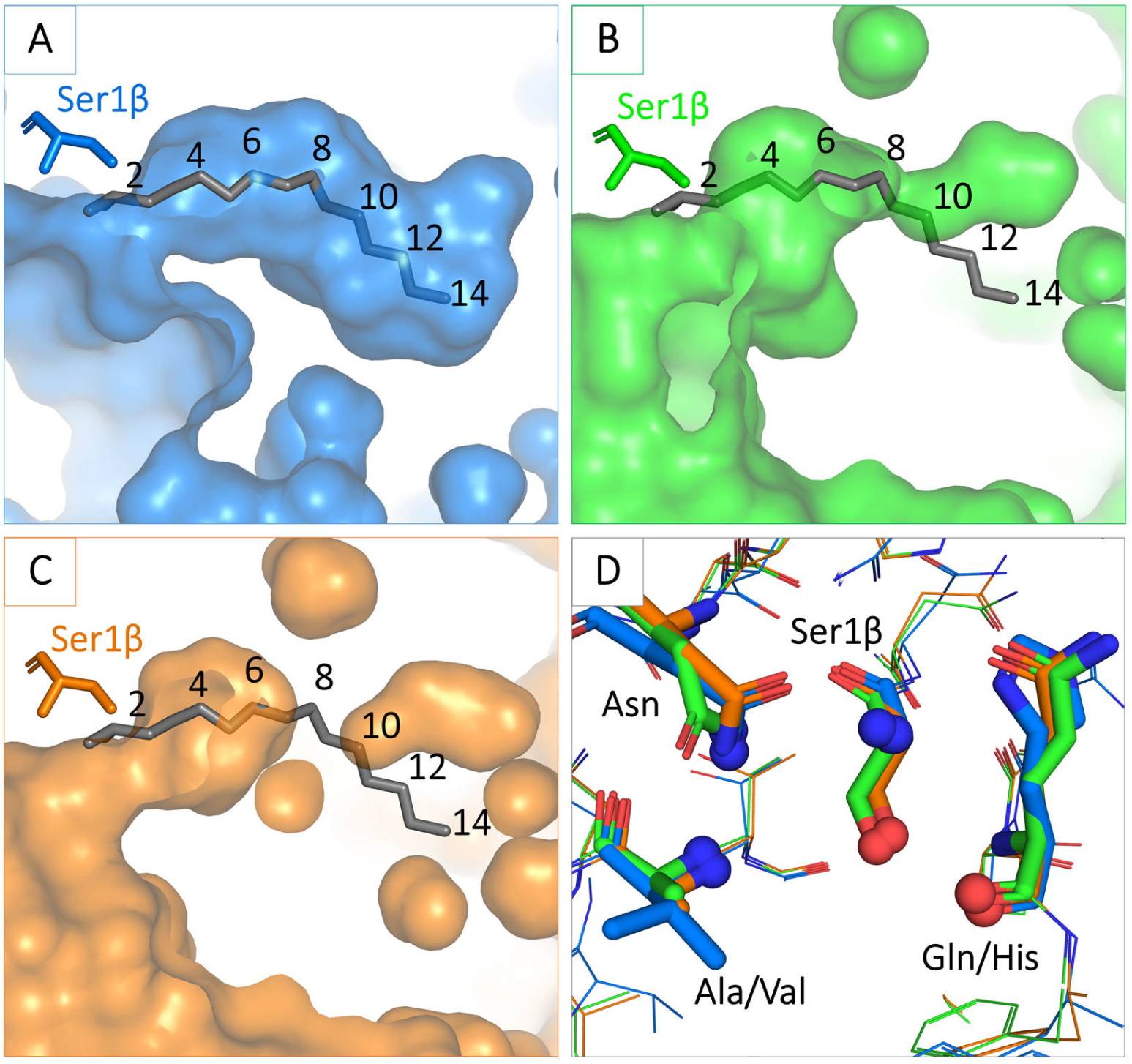
Structural prerequisites for quorum quenching activity in N-terminal serine hydrolases. Geometries of substrate binding pockets in the crystal structures of **A**) paPvdQ (PDB-ID: 4M1J), **B**) kcPGA (PDB-ID: 4PEL), and **C**) ecPGA (PDB-ID: 1GK9). **D**) Conserved positioning of functional groups in catalytic residues across all three enzymes. Gray sticks in A-C represent a transition state analog of paPvdQ, highlighting its acyl-binding subpocket, which is suitable for binding long-chain HSLs. The superposition of this analog on the structures of the PGAs shows that these enzymes could potentially accommodate HSLs of up to 8 carbon atoms.

Whereas paPvdQ is selective for long HSLs, *Kluyvera citrophila* penicillin G acylase (kcPGA) was recently shown to display QQ activity towards short HSLs. Accordingly, analysis of its crystal structure revealed that its acyl-binding cavity is significantly shallower than that of paPvdQ (**Figure 1B**).^51^ However, other PGAs with high levels of sequence and structure similarity to kcPGA (**Figure S1, Table S1**), including the well-characterized *Escherichia coli* PGA (ecPGA, **Figure 1C**), exhibit no appreciable QQ activity.^52^ Since ecPGA, kcPGA, and paPvdQ all have similarly positioned functional groups (**Figure 1D**), it is believed that the observed differences in their activity arise from differences in their functional dynamics.^51,53,54^

To uncover unknown determinants governing the QQ activity of N-terminal serine hydrolases, our study focuses on the role of dynamics in the molecular function of the prototypical enzyme paPvdQ and the related but inactive ecPGA, which serves as a negative control. By comparing the plasticity and pre-organization of their active site residues, their ability to stabilize productive substrate binding modes, and the atomistic details of their reaction mechanisms, we reveal crucial structure-dynamics-function relationships relevant to the future discovery and design of robust enzymatic QQ agents that can replace or complement antibiotics and selectively combat resistant bacteria species.

## METHODS

A full description of the simulations and system setup is provided in the Supporting Information.

### Molecular dynamics of free enzymes

Crystal structures of ecPGA (PDB-ID: 1GK9) and paPvdQ (PDB-ID: 4M1J) were obtained from the PDB database.^55,56^ For *Achromobacter spp*. penicillin G acylase (aPGA), a previously derived homology model was retrieved from the Protein Model Database (ID: PM0080082)^57^ and corrected using the RepairPDB module of FoldX.^58^ Structures were protonated with the H++ webserver^59–61^ at pH 7.5 using the default salinity and internal and external dielectric constants. Protonated structures including crystallographic waters were placed in a truncated octahedral box of TIP3P waters whose boundaries were at least 10 Å away from all atoms in the structure and neutralized using Na+ and Cl-ions to approximately 0.1 M concentration. Initial parameters and topologies were generated using the tleap module of AmberTools17.^62^ Hydrogen atom masses were repartitioned to enable a 4 fs time-step during simulations with the SHAKE algorithm.^63,64^ Energy minimization and molecular dynamics (MD) simulations were performed using the ff14SB force field^65^ with the pmemd and pmemd.CUDA modules of the Amber16 package, respectively.^62^ The systems were energy minimized and equilibrated, after which a 500 ns NPT production MD simulation was performed at 310 K using the Langevin thermostat.^66^ All steps were performed in triplicate to generate three independent replicates. The stability of the production phase was evaluated in terms of the root-mean-square deviation (RMSD) of the backbone heavy atoms (**Figure S3 and S20**).

### Free enzyme dynamics analysis

The opening of the acyl-binding cavity across free enzyme MD trajectories was investigated using the CAVER 3.0.2 program.^67^ The starting point for the calculation of paths was chosen based on the center of mass of three residues: Met142α, Ser67β, and Ile177β for PGAs, and Leu146α, Leu53β, and Trp162β for paPvdQ.^49^ Potential acyl-binding site opening events were identified using a probe radius of 0.5 Å. A time sparsity of 10 was used to reduce the computational cost of this analysis. The paths were clustered using a threshold of 3.5, and one representative path was selected from each cluster.

The cpptraj module of AmberTools17 was used to measure relevant distances between functional atoms in the catalytic residues across all MD trajectories. All distances presented in this work stand for internuclear distances. In addition, the network of distances was subjected to dimensionality reduction using principal component analysis (PCA) as implemented in the scikit-learn Python library.^68^

### Receptor selection and molecular docking

Protein conformations harboring spatially well-shaped acyl-binding cavities and favorably pre-organized catalytic machinery were selected as receptor structures for ligand docking according to the criteria presented in **Table S2** (“*receptor selection for docking*”). These criteria were used to promote proper orientation of the hydroxyl hydrogen of the nucleophilic serine relative to its amine group, which serves as a hydrogen acceptor in the first reaction step, and to ensure appropriate pre-organization of the catalytic machinery (i.e., the nucleophile serine and the oxyanion hole stabilizing residues) for reactive binding of the substrate.^53,57,69,70^

Molecular docking of C06- and C08-HSLs was performed using Autodock4.2.6,^71^ Representative protein-ligand complexes were selected for the following round of simulations based on the favorability of their Autodock binding scores and the mechanism-based selection criteria^72^ specified in **Table S2** (“*selection of complexes for MDs*”), which were chosen to ensure an acceptable distance between the nucleophile and electrophile as well as proper stabilization of the oxyanion hole.

### Molecular dynamics of protein-ligand complexes

Systems were prepared using an approach analogous to that applied in the free enzyme simulations with the tleap and Parmed modules of AmberTools18.^73^ Minimization and equilibration were performed using the protocols applied in the free enzyme simulations with additional restraints on the ligand molecule. Unrestrained NPT production simulations of 50 ns were performed at 310 K using the Langevin thermostat. Fifteen independent replicates (each including the minimization and equilibration steps as well as the production run) were performed for all ligand-protein simulations. The stability of the production runs was evaluated in terms of the RMSD of the protein backbone heavy atoms, as in the free enzyme simulations (**Figure S4-S5** and **S21**), using the Cpptraj module of AmberTools18.

### Complexes binding free energy and ligand stabilization estimation

The MMPBSA.py module of the AmberTools18 package was used to estimate the binding free energy of the complexes using the molecular mechanics/generalized Born surface area approach (MM/GBSA).^74^ Calculations were performed with the Generalized Born implicit solvent model 8 at a salt concentration of 0.1 M. A per-residue binding free energy decomposition was generated and the results were filtered to extract the residues with the greatest contributions, i.e. those with contributions below −0.5 kcal/mol (for favorable contributions) or above 0.5 kcal/mol (for unfavorable contributions).

### Quantum Mechanics/Molecular Mechanics MD simulations

QM/MM MD simulations were performed using the sander module of the Amber18 package. The initial frame for each protein-ligand complex was extracted from the standard MD simulations by searching for reactive-like configurations satisfying the criteria listed in **Table S2** (“*selection of representative configurations for sMDs*”). Systems were equilibrated by performing a 10 ns NPT simulation with the same settings as for protein-ligand production runs with additional 25 kcal mol^-1^ Å^-2^ harmonic restraints. Production simulations of 500 ns were then performed for each complex with distance restraints on selected key interactions (**Table S2**, *“restraints for sMDs inputs generation*”), collecting restart files to obtain an ensemble of 500 equivalent starting points per complex. Each starting conformation was then equilibrated in QM/MM MD simulations while applying the distance restraints specified in **Table S2** (*“restraints for 1*^*st*^ *QM/MM equilibration”*). The QM region consisting of the HSL substrate and selected active site residues (**Figure S2**) was described using the PM6-D semi-empirical method,^75–77^ while the rest of the system was treated at the MM level using the ff14SB force field.

In addition, steered QM/MM MD simulations of the acylation process were performed. The process was divided into two steps: formation of the tetrahedral intermediate (TI) and collapse of the TI to form the acyl-enzyme (AE). In the first step, the reaction coordinate (RC) was represented as a linear combination of distances (LCOD) involved in the proton transfer. The linear combination was d1 – d2, where d1 is the distance from the serine amine nitrogen to the proton of the hydroxyl group and d2 is the distance between the proton and the oxygen of the hydroxyl group as shown in **Figure 2A**. This RC was steered from 1.1 to −1.1 Å with a harmonic restraint of 1000 kcal mol^-1^ Å^-2^. Correctly formed TIs were then equilibrated on the same level of theory as before, with a distance restraints on the newly formed covalent bond (1.5 Å) and the distance from the serine amine hydrogen to the substrate’s amide nitrogen (2.0 Å), which serves as a proton acceptor in the next step, as shown in **Table S2** (*“restraints for 2*^*nd*^ *QM/MM equilibration”*).

**Figure 2.**
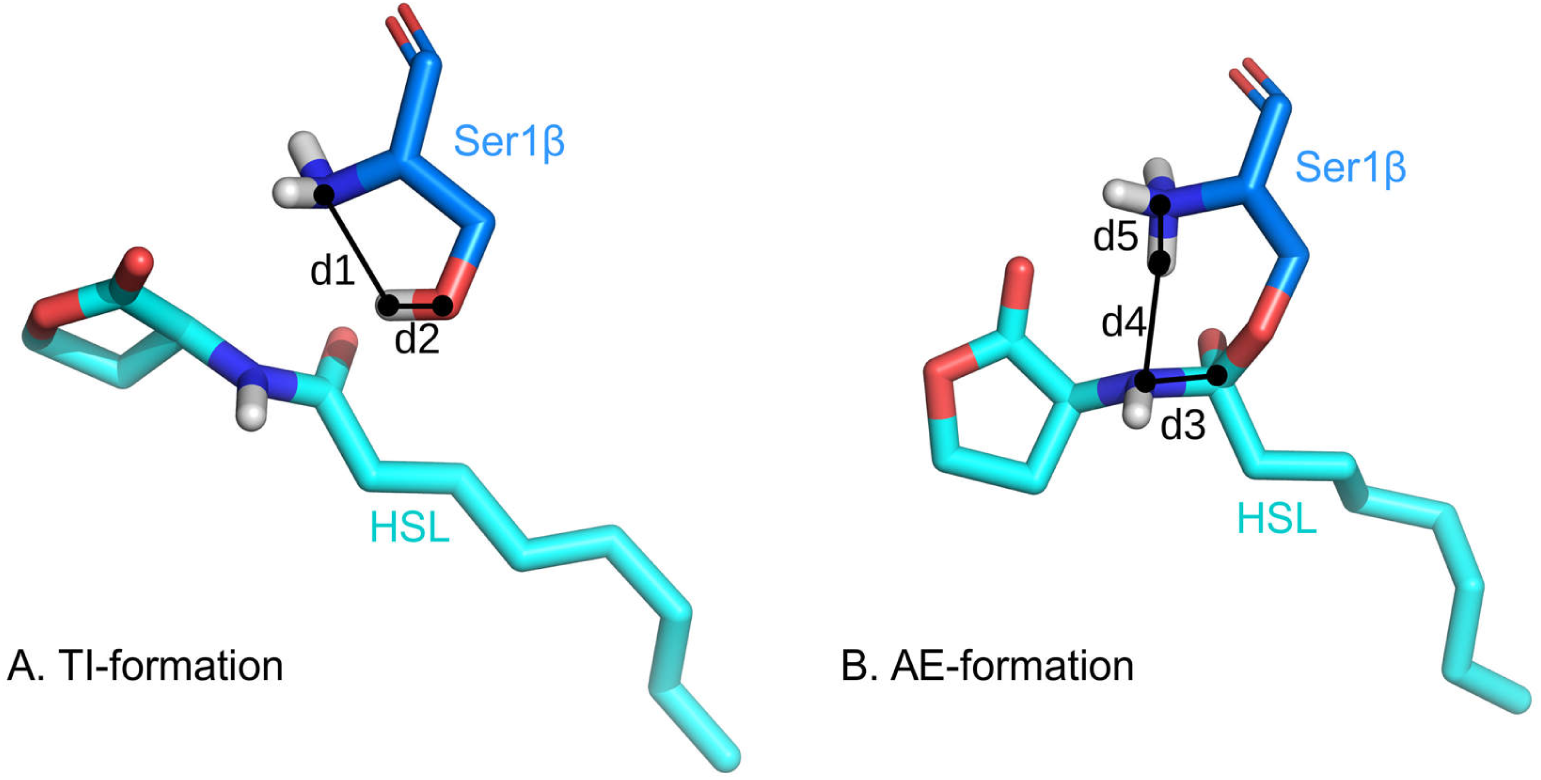
Linear combinations of distances (LCOD) representing the reaction coordinates for the two steps in the steered MD simulations of HSL acylation. A) LCOD for tetrahedral intermediate formation, consisting of the distance from the transferred hydrogen to the amine nitrogen and the distance from the same hydrogen to the hydroxyl oxygen (d1-d2, steered from 1.1 to −1.1 Å). B) LCOD used in the second step of the acylation leading to formation of the acyl-enzyme, consisting of the length of the substrate amide bond and the distances from the transferred hydrogen to the amide nitrogen of the substrate and the amine nitrogen of the serine residue (d3-d4+d5, steered from 0.6 to 4.9 Å).

Successfully equilibrated TIs were used as starting points for the second step of the acylation reaction. Here, the RC was represented as a LCOD of three distances relevant to the formation of the AE, as shown in **Figure 2B:** d3 – d4 + d5, where d3 is the distance from the carbonyl carbon of the HSL to its amide N, d4 is the distance from the amide N to the closest proton of the serine amino group, and d5 is the distance from the nitrogen of the serine amino group to the proton being transferred. This RC was steered from 0.6 to 4.9 Å with a harmonic restraint of 1000 kcal mol^-1^ Å^-2^. The results of these simulations were used as inputs to generate potential of mean force (PMF) profiles using Jarzynski’s equality.^78^ Errors in the PMF profiles were estimated using a block-averaging scheme.

### Microorganisms and culture conditions

The production recombinant strains *E*.*coli* RE3(pKA18)^79^ and *E. coli* BL21(pKX1P1)^80^ grown in a stirred bioreactor as fed-batch cultures were used to prepare biomass for purification of ecPGA and aPGA. The cultivations were performed in a Biostat MD bioreactor (B. Braun Biotech Int., Melsungen, Germany) with a working volume of 6 l at 28 °C for 24 h. Both strains were cultivated in M9 minimal medium (0.4 % (NH_4_)_2_SO_4_, 1.36 % KH_2_PO_4_, 0.3 % NaOH, 0.2 % MgSO_4_·7H_2_O, 0.02 % CaCl_2_·6H_2_O, 0.01 % FeSO_4_·7H_2_O, pH 7.2) supplemented with glycerol (10 g/l) and casein hydrolysate (10 g/l) as carbon and energy sources for the strain *E. coli* BL21 (pKX1P1) and sucrose (10 g/l) for the strain *E. coli* RE3 (pKA18). A 40% glycerol solution was used to feed *E. coli* BL21 (pKX1P1) and a 50% sucrose solution was used for *E. coli* RE3 (pKA18) when the concentration of carbon sources in the bioreactor dropped to zero. The fermentation operating parameters were: initial stirrer speed 300 rpm, airflow rate 1 vvm, and pH 6.5 maintained using 25% NH_4_OH. The concentration of dissolved oxygen (pO_2_) was maintained at 20% of the value in air-saturated medium by cascade regulation of the stirring frequency during the initial batch phase of the culture. During the fed-batch phase, the stirring rate was set to 800 rpm and feeding was controlled to maintain a constant level of dissolved oxygen (pO_2_ of 20%). A culture from 200 ml of minimal medium M9 grown for 16 h at 28 °C was used as the inoculum.

### Enzyme purification and hydrolytic activity assay

aPGA was purified as described by Škrob *et al*.^81,82^ and ecPGA according to Kutzbach and Rauenbusch.^83^ The activity of 1 unit (U) was defined as the amount of aPGA or ecPGA cleaving 1 μmol of the corresponding HSL per minute in 0.05 M sodium phosphate buffer containing 2 % (w/v) HSL at 35 °C and pH 8.0 or 7.0, respectively. All HSLs were obtained from Sigma-Aldrich.

### Effect of pH and temperature on activity of PGAs

The optimal temperature for activity was determined by conducting cleavage experiments at temperatures from 30 °C to 70 °C in 0.05 M phosphate buffer at pH values from 5.0 to 8.0 for both studied HSL signaling molecules. For the ecPGA enzyme, the maximum activity was observed at 35-40°C and pH 7.0, whereas the activity of aPGA peaked at 50 °C and pH 8.0.

### Contribution of autohydrolysis to activity of PGAs

To estimate the extent of substrate autohydrolysis occurring during the enzymatic reactions, the stability of C06-HSL was evaluated at substrate concentrations of 5, 10, and 20 mM in 0.05 M phosphate buffer at pH 7 and 35 °C.

### Enzyme kinetics

Kinetic characterization of HSL degradation by ecPGA and aPGA was carried out in 0.05 M phosphate buffer at pH 7.0 and 8.0, respectively, at the optimal temperature for ecPGA (35°C). Concentrations of reactants were monitored by HPLC. Aliquots for analysis were adjusted to pH 2 to terminate the reaction and then evaporated to dryness at 35°C, after which the residues were dissolved in 0.2 mL of HPLC grade acetonitrile. A 20 μL aliquot of the resulting solution was loaded onto an analytical RP-C18 HPLC column (250 × 4.6, 5 μm particle size, Hypersil ODS) and eluted isocratically in methanol-water (50:50, v/v) for 10 minutes, followed by a linear gradient from 50 to 90 % methanol in water over 15 minutes and isocratic elution for 25 minutes. The flow rate was 0.4 mL/min and the eluate’s absorption was monitored at 210 nm. The retention times of the substrates of interest are listed in **Table S4**. The relationship between the initial reaction rate and the substrate concentration (between 1 and 1000 μM) was determined for each substrate in three independent experiments. The kinetic parameters *K*_M_ and *V*_max_ were calculated using Hans-Volf plots and an ANOVA calculator.

### Confirmation of the enzymatic activity

To verify that the observed quorum quenching activity was coupled to the action of ecPGA, its activity was determined at increasing enzyme concentrations of 1.5, 3.0, 30 and 250 μM. The reaction was started by adding the enzyme to a reaction mixture containing 10 mM of C06-HSL as a substrate in 0.05M phosphate buffer at pH 7 and 35°C.

## RESULTS

### The arrangement of the acyl-binding cavity and catalytic machinery in ecPGA and paPvdQ enables productive binding and stabilization of moderately long HSLs

The behavior of the acyl-binding cavities and catalytic residues of the paPvdQ and ecPGA enzymes in the absence of HSLs was analyzed in three 500 ns long MD simulations. We initially used the CAVER tool to explore the ability of these cavities to open their entrances wide enough to admit the acyl-chains of the substrate molecules.^84^ Opening events were frequent; the entrances were open for least 10 % (ecPGA) or 20 % (paPvdQ) of the simulated period in each of three replicates (**Figure S6-S7**), providing ample opportunities for ligands to access the acyl-binding sites. Next, we investigated the overall geometric profiles of the acyl-binding cavities and the degree to which their shapes fit the shortest ligand under consideration (C06-HSL), considering not just the entrance bottleneck but also the depth of the cavity and the location of the entry relative to the catalytic residues (**Table S5**). An analysis of all the open conformations of the binding sites revealed that the cavity entrance was broader in ecPGA than in paPvdQ but the cavity of paPvdQ was more than twice as long as that of ecPGA (**Figure S8)**. In both enzymes, the acyl-binding cavities adopted geometries that enabled binding of short-chain HSLs (**Table S5)**. These geometries are consistent with those seen in the corresponding crystal structures and imply that ecPGA is open with sufficient frequency for effective catalysis and has a geometry suitable for the binding of HSLs with short and intermediate length acyl chains, indicating that its lack of activity toward such compounds has some other cause.

To better understand the different catalytic behaviors of ecPGA and paPvdQ, we studied the dynamics of crucial functional atoms of catalytic residues. Mechanism-based geometric criteria defining the atomic arrangements required for the activity of N-terminal serine hydrolases can be expressed in terms of three criteria that must be satisfied simultaneously (**Figure 3**): there must be a relatively short nucleophile attack distance (< 3.3 Å), it must be possible to form two stabilizing hydrogen bonds with oxyanion hole stabilizing residues, and the nucleophile attack angle must be ∼90°.^53,57,85^

**Figure 3.**
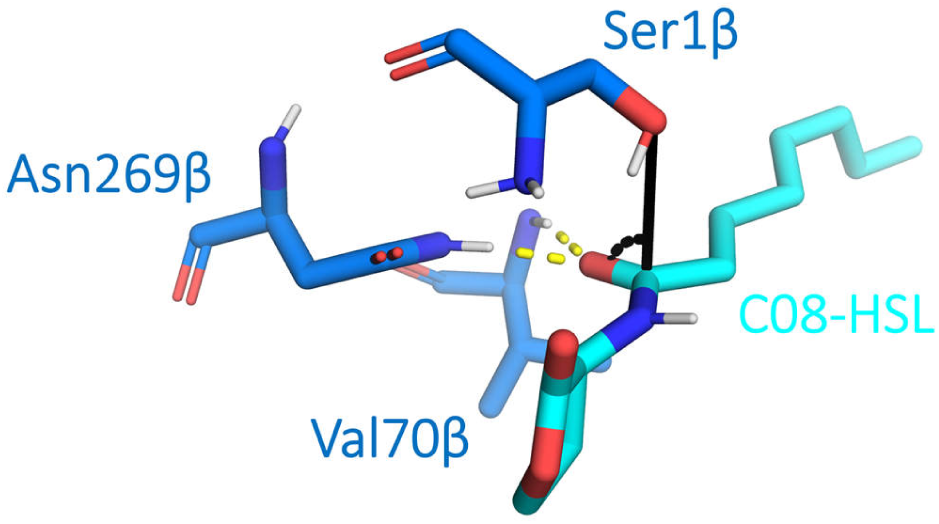
Interaction criteria for productive stabilization of HSL, illustrated using the paPvdQ-C08-HSL complex. The criteria relate to the nucleophile attack distance and angle (black lines) and the potential to form two oxyanion-stabilizing hydrogen bonds (yellow dashed lines).

In light of these criteria, we focused on the following geometric parameters: (i) the distances between the N-terminal serine hydroxyl oxygen (Ser1β-O-hydroxyl) that serves as the initial nucleophile in the catalytic process^86^ and the two nitrogen atoms of the oxyanion hole stabilizing residues that function as hydrogen bond donors (Ala69β and Asn241β in ecPGA; Val70β-N-backbone and Asn269β-N^δ^ in paPvdQ),^70^ (ii) the distance between the two oxyanion hole stabilizing hydrogen bond donors, and (iii) the distance between the Ser1β-O-hydroxyl and the backbone oxygen of a nearby glutamine/histidine residue (Gln23β/His23β-O-backbone) known to stabilize the ligand during the reaction.^48^ To map the most important conformational transitions of the catalytic machinery (**Figure 4**), PCA was applied to the networks of these distances. The first and second principal components (PC1 and PC2) cumulatively explained 78% and 84% of the total variance in these distances for ecPGA and paPvdQ, respectively. PC1 mainly reflected a change in the separation of Asn241β/Asn269β from Ser1β and Ala69β/Val70β, whereas changes along PC2 were mainly governed by the distance between Ser1β and Gln23β/His23β (**Table S9**).

**Figure 4.**
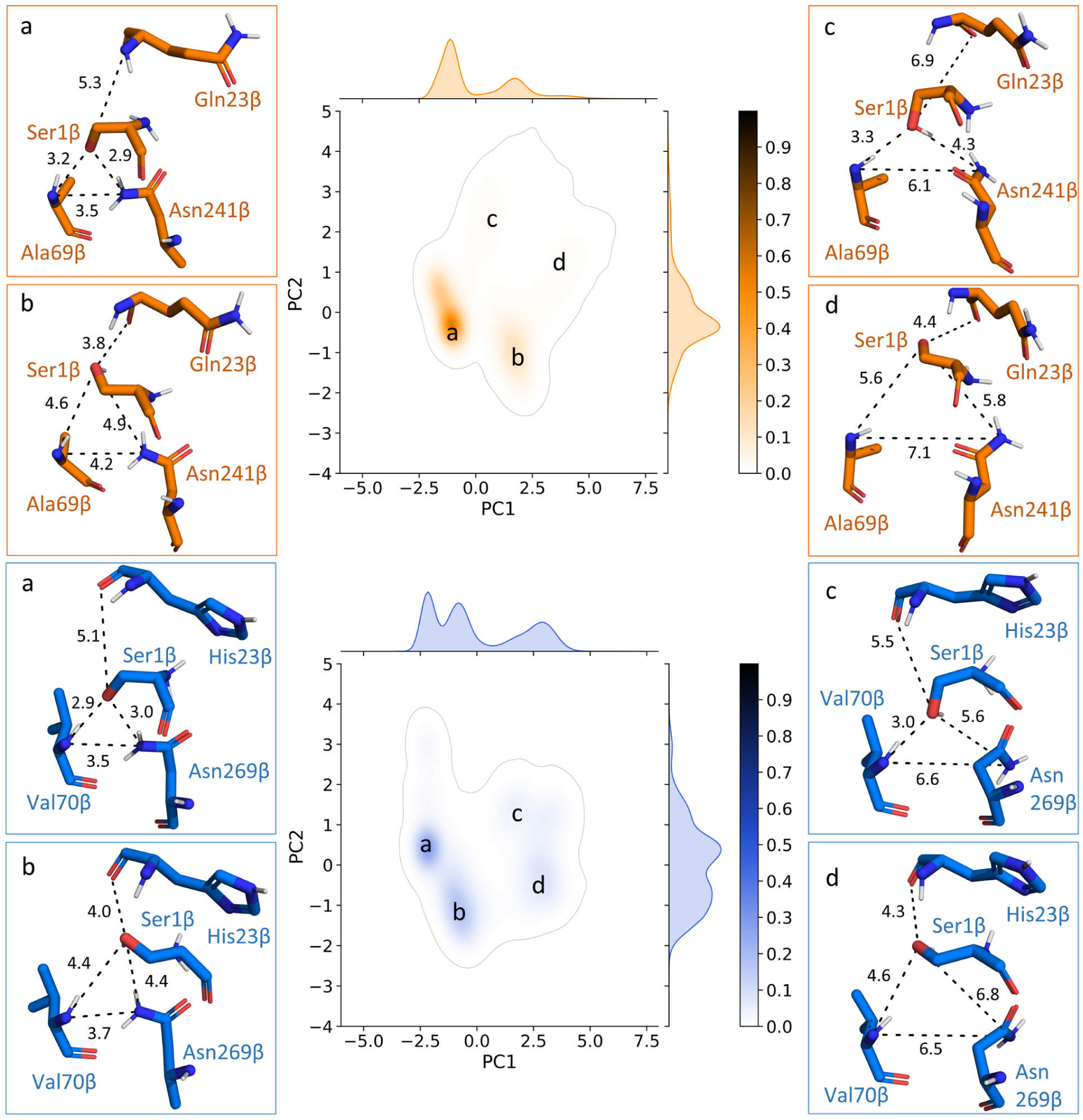
Primary conformations adopted by the catalytic machinery of paPvdQ and ecPGA and PCA analysis of distances between catalytic residues in ecPGA (top, orange) and paPvdQ (bottom, blue) during MD simulations without substrates. In state b, the catalytic residues are favorably pre-organized for productive substrate binding. States a, c, and d require conformational rearrangement to enable productive substrate binding.

Based on the highest density peaks in the conformational landscape formed by these two PCs, we identified the four most frequently visited **states a-d** (**Figure 4** and **Table S6)**. Comparative analysis of both enzymes showed that the four identified states for each enzyme correspond well to each other in terms of the arrangement of catalytic residues. In particular, in both enzymes, **state b** exhibits a favorable pre-organization of the catalytic site for productive stabilization of HSLs, with the nucleophilic serine oxygen being at similar distances from the functional atoms of Ala69β/Val70β, Asn241β/Asn269β, and Gln23β/His23β while also forming a hydrogen bond with the backbone hydrogen of Gln23β/His23β (**Table S6**). This arrangement permits stabilization of HSL in the proper orientation for nucleophilic attack, i.e. with the Ser1β hydroxyl oxygen positioned over the carbonyl carbon of the ligand’s amide moiety with a nucleophilic attack angle of approximately 90°. In the remaining three states, conformational rearrangements would be needed to enable HSL binding and nucleophilic attack. In **state a**, the nucleophile serine hydroxyl group was within hydrogen bonding distance of Ala69β/Val70β and Asn241β/Asn269β but was more distant from Gln23β/His23β (**Table S6**). Additionally, the hydroxyl group of Ser1β blocked access to the oxyanion hole stabilizing residues, essentially preventing the binding of HSLs in reactive-like poses. Interestingly, **state a** was more common than **state b** in the apo-form of ecPGA but paPvdQ explored these states almost equally, suggesting that paPvdQ is better pre-organized for productive HSL binding and stabilization. In **state c** the nucleophile serine hydroxyl group is within hydrogen bonding distance of Ala69β/Val70β as in **state a**, preventing potential oxyanion hole stabilization. Moreover, the conformation of Asn241β/Asn269β is flipped relative to that in **states a** and **b**, placing its crucial nitrogen atom too far from the remaining elements of the catalytic machinery for efficient HSL stabilization. The geometry of **state d** resemble **state c** in terms of the flipping of the Asn241β/Asn269β side-chain and **state b** in terms of the geometry of the remaining residues. **State d** would require conformational rearrangement of Asn241β/Asn269β to achieve favorable pre-organization for ligand stabilization. Importantly, **states c** and **d** were visited markedly less frequently than the other two states (**Table S6**), making **states a** and **b** the most prominent of the four chosen for analysis.

Detailed investigation of binding site dynamics and the behavior of catalytic residues thus revealed states of both ecPGA and paPvdQ that are sufficiently open and appropriately pre-organized for HSL binding (**Table S5**). Next, we performed molecular docking of two moderately long HSLs: C06-HSL, which is cleaved by kcPGA^51^, and C08-HSL, one of the shortest confirmed substrates of paPvdQ.^47,49^ The docked poses were subjected to geometric filtering using the previously discussed mechanism-based geometric criteria to select complexes in configurations that stabilize the ligand and nucleophilic serine sufficiently for nucleophilic attack to proceed. In this way, two or three well-stabilized reactive binding poses were identified for each complex, with all distances satisfying the specified criteria and favorable binding energies according to the Autodock4.2 scoring function (**Table S7)**.^71^ This shows that both enzymes have acyl-binding cavity geometries that are favorable for binding moderately long HSLs while maintaining the catalytic machinery in an arrangement that is properly pre-organized for catalysis.

### Both enzymes maintain productive stabilization of HSL binding, but paPvdQ does so to a slightly greater degree than ecPGA

To compare the ability of both enzymes to maintain the productive binding of HSLs, we performed 15 replicate simulations of each enzyme starting from their bound poses. Following the approach adopted by Novikov *et al*.^53^ we established a set of analogous geometric criteria that must be satisfied to provide productive stabilization of the ligand in the acyl-binding site while also enabling initiation of the nucleophilic attack reaction. As discussed in the preceding section, these criteria related to the nucleophile attack distance, the hydrogen bond stabilization provided by the oxyanion hole stabilizing residues, and the nucleophile attack angle. The latter parameter was required to be between 75° and 105°, and reflects the accessibility of the carbonyl carbon of the substrate’s amide group (**Figure 3**).

Initially, all stabilization components were evaluated simultaneously in two stages. First, we searched for all states in which the nucleophile distance, angle, and oxyanion hole criteria were satisfied simultaneously. We then quantitatively investigated whether these states were adopted repeatedly in the replicate simulations to verify that their detection did not simply represent an insignificant incident. The stabilized state was sustained somewhat more frequently in paPvdQ than in ecPGA (**Figure S9-S10)**, but in both cases it occurred with sufficient frequency for catalysis. The difference between the two enzymes was more pronounced when their complexes with C08-HSL were examined: repeated formation of the productive configuration was observed in 3 replicates for ecPGA and 11 replicates for paPvdQ. This outcome is consistent with the preference for shorter substrates observed in kcPGA, which is closely related to ecPGA.^51^ In contrast, the repeated productive stabilization of the C06-HSL complex of ecPGA was very similar to that for paPvdQ: repeated stabilization was observed in 9 and 13 replicate simulations of these complexes, respectively. Interestingly, ecPGA provided very persistent stabilization of this shorter ligand in productive configurations that were maintained for almost the entire duration of the simulation in some replicates (**Figure S9**, replicates 8 and 14 for ecPGA-C06-HSL).

We next dissected the individual contributions of each substrate stabilizing feature. The nucleophile attack distance criterion was satisfied in simulations of both enzymes’ complexes with C06-HSL and C08-HSL. However, there was a clear preference for paPvdQ in that the ligand’s positioning generated an optimal attack distance most frequently in this enzyme (**Figure 5)**. The same was true with respect to the distribution of the nucleophile attack angle (**Figure 5)**: paPvdQ frequently bound HSL in an orientation that exposed the carbonyl carbon to nucleophilic attack, whereas the angle distribution of ecPGA was significantly wider and the attack angle was more frequently in an unfavorable range. Analysis of the distances relevant to stabilization of the oxyanion hole showed that these interactions were also maintained in both proteins (**Figure S11**). Interestingly, the distance between the HSL carbonyl oxygen and the oxyanion-stabilizing asparagine is larger in paPvdQ complexes; this is consistent with the PCA results for the free-enzyme simulations, which showed that **states c** and **d** (**Figure 4**) with the flipped asparagine conformation were more prevalent in paPvdQ than in ecPGA.

**Figure 5.**
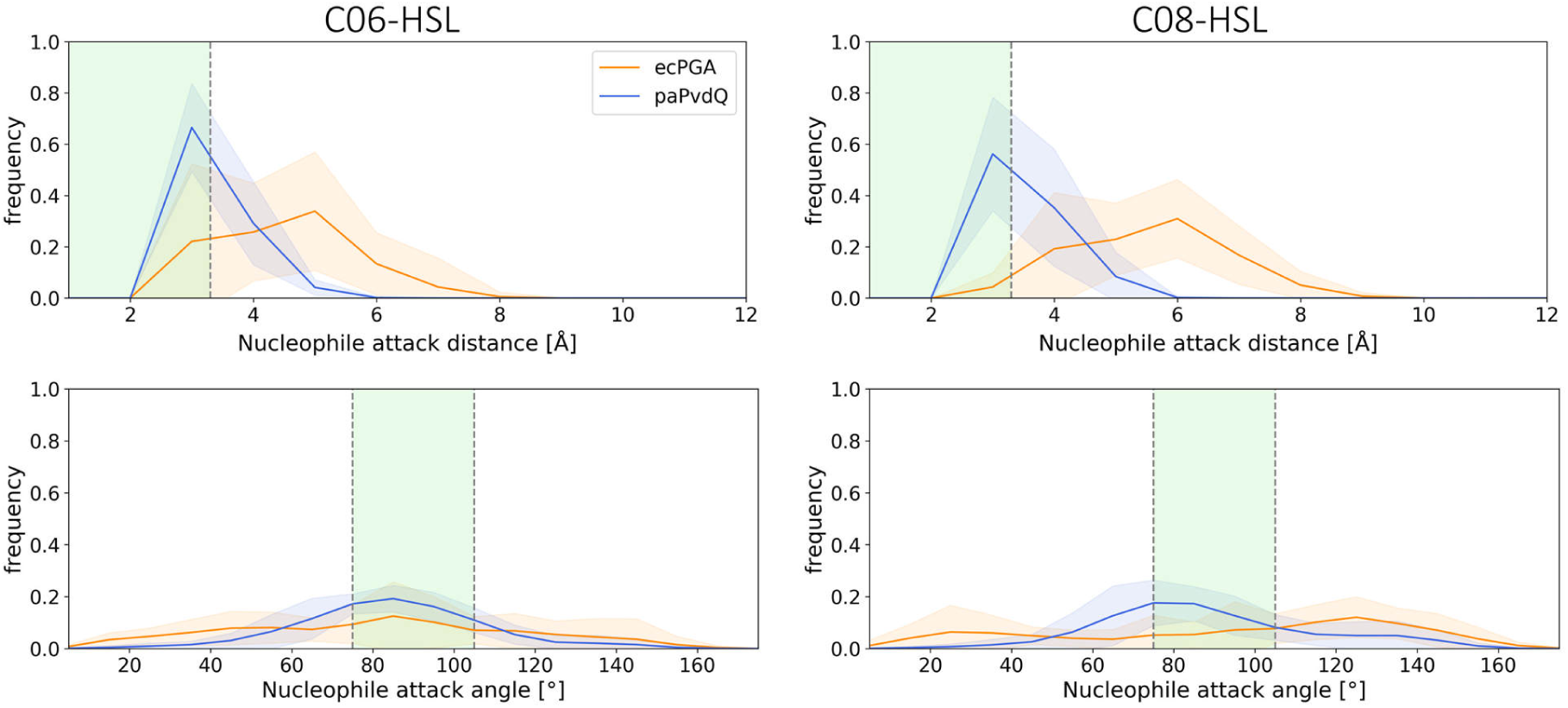
Maintenance of properly stabilized protein-ligand complexes. Distribution of nucleophile attack distance (upper plots) and angle (bottom plots) for ecPGA (orange) and paPvdQ (blue) enzymes in complex with C06-HSL (left) and C08-HSL (right) from simulations where the average ligand RMSD did not exceed 5.0 Å (**Figure S12-13)**. Regions of the plots highlighted in green indicate the optimal ranges of the parameters.

To further understand factors that contributed to the apparent difference in substrate stabilization, we explored the mobility of individual heavy atoms of HSL (**Figure 6)**. These analyses consistently indicated that the most mobile part of the ligand in all complexes was the exposed lactone ring, followed by the atoms of the scissile amide bond stabilized by interactions with residues of the catalytic machinery. Least mobile was the acyl chain, which was buried in the binding site. For the complexes of C06-HSL, the atomic fluctuations in both enzymes were similar, although the standard deviations were slightly higher for ecPGA than for paPvdQ. More pronounced differences were seen with the longer substrate; although the overall trend in the fluctuations was similar for both systems, the absolute fluctuations were larger in ecPGA. This correlates well with the two enzymes’ acyl-binding cavity profiles (**Figure S8**), which showed that the entry to the acyl-binding pocket of ecPGA was wider and shallower, giving the bound ligand more conformational freedom and thus producing less optimal stabilization, especially for the longer substrate that could not be fully accommodated inside the pocket.

**Figure 6.**
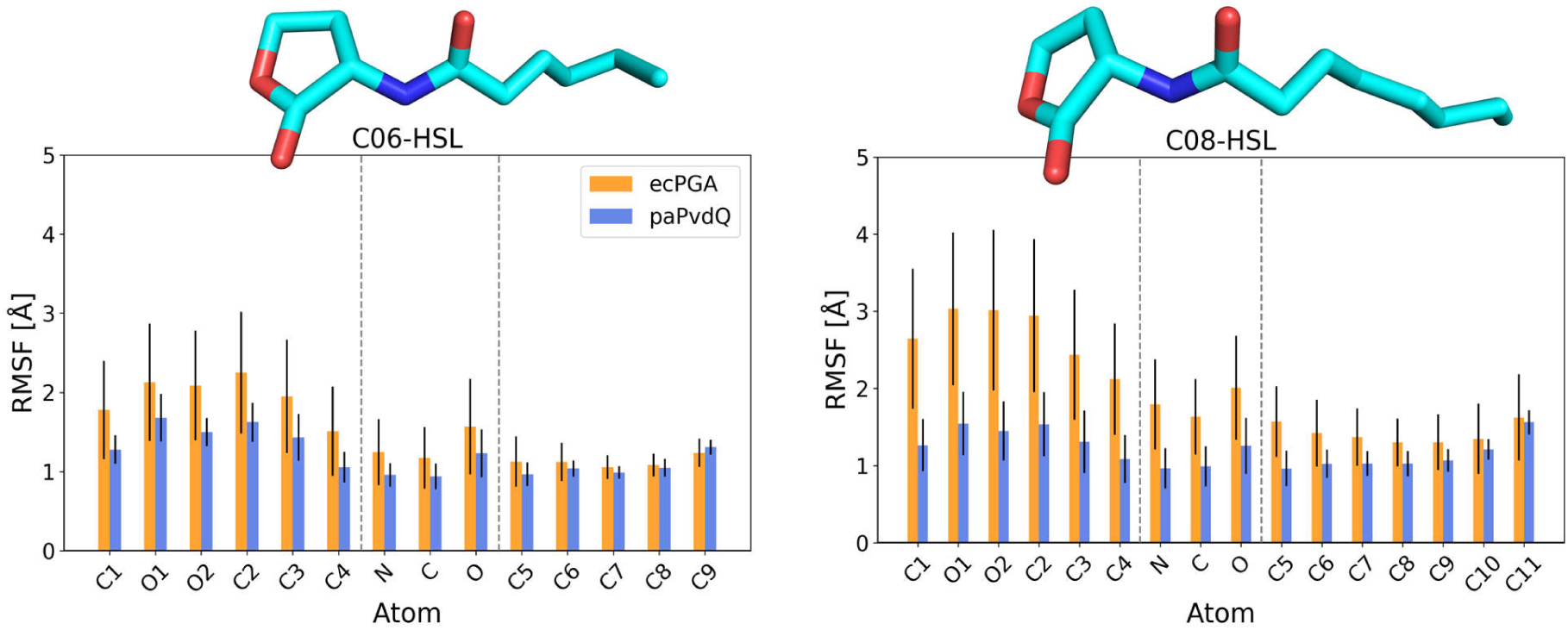
The mobility of the HSL atoms during simulations. Root-mean-square fluctuation (RMSF) of HSL heavy atoms in simulations where the average ligand RMSD did not exceed 5.0 Å (**Figure S12-13)**.

Finally, we estimated the binding free energies of productively bound protein-ligand complexes using the MMGB/SA method. The obtained binding energies were favorable for both ligands in both paPvdQ and ecPGA and proportional to the size of the ligand (**Table S8**). A per-residue decomposition of these binding free energies showed that the residues exhibiting the largest differences between the two enzymes were Met142α, Phe146α, and Phe256β in ecPGA. All of these residues are located in regions where the backbones of the two enzymes cannot be spatially aligned (**Figure S14**). Met142α forms the bottom part of the cavity in ecPGA. In paPvdQ, this cavity is buried significantly more deeply in the protein core. Phe146α is located at the entrance to the acyl-binding cavity, which is substantially wider in ecPGA than in paPvdQ (**Figure S8**). Phe256β in ecPGA interacts with the lactone ring of the ligand from the outside of the cavity; this interaction was only observed for this enzyme, and only when complexed with C06-HSL. While these residues contribute favorably to the binding of short to medium-length ligands, they could potentially hinder the efficient binding of longer substrates, as observed for kcPGA.^51^ In contrast, paPvdQ residues Phe32β and Trp186β (**Figure S14**), which are located approximately in the middle of the acyl-binding cavity (corresponding to the bottom part of the cavity in ecPGA), contributed favorably because they narrow this part of the cavity, stabilizing the substrate’s alkyl chain and restricting its mobility. The remaining residues did not differ significantly between the enzymes in terms of their contributions to the binding free energy.

### The initiation of acylation in N-terminal serine hydrolases

Encouraged by the computational results indicating that ecPGA can productively bind and stabilize HSLs, we moved to investigate its putative catalytic mechanism. Here we focused on the acylation step, which is assumed to be rate-limiting for the hydrolysis of various substrates by acylases and serine proteases.^87–89^ We began this process by selecting the protein-ligand complexes from the previous simulations that had the most favorable configurations for the reaction (**Table S10**) and performed restrained simulations to generate a uniform ensemble of Michaelis complexes (MCs) as starting positions for replicated steered QM/MM MD simulations (**Figure S15**). Next, we performed QM/MM MD simulations mimicking the mechanism defined by Grigorenko *et al*. for ecPGA with its native substrate, penicillin G.^69^ This acylation proceeds via two steps - nucleophilic attack by Ser1β, which is directly activated by its own α-amino group leading to the formation of the TI, followed by the decomposition of TI into the AE complex and release of the homoserine lactone as the first reaction product (**Table S11**).

### Nucleophilic attack is concerted with internal proton transfer from the serine hydroxyl to its amine group and is feasible in both enzymes

Starting from the MCs (**Figure 7** and **Table S12**), we simulated the proton transfer from the hydroxyl oxygen to the amine nitrogen of the nucleophile Ser1β (**Figure 2A**). During the transfer process, we observed a spontaneous nucleophilic attack in which the distance between the nucleophilic oxygen and the electrophilic carbon of the HSL contracted by up to 0.5 Å in the MC (RC 1.1 Å) and TS1 (RC ∼ 0.2 Å) states (**Table S12**). This apparently concerted mechanism resembles the mechanism of penicillin G hydrolysis by ecPGA.^69^ Interestingly, the shortening of the carbon-oxygen distance became apparent when the proton was positioned approximately equidistantly between the oxygen and nitrogen centers (RC ∼ 0.0 Å) for all systems except ecPGA-C08-HSL, in which the formation of TS1 and the subsequent formation of a covalent bond between the substrate and Ser1β were notably delayed to RC 0.1 Å and RC −0.2 Å, respectively (**Figure 7** and **Figure S16**). The first step was considered to be complete once the proton was fully transferred to the amine nitrogen of Ser1β and a stable TI was formed (**Table S12**). Visual inspection of the TI structure revealed the following structural features (**Table S12**): the presence of a 1.5-1.6 Å covalent bond between the nucleophilic oxygen and the attacked carbon of the substrate, stabilization of the oxyanion of the TI by enhanced hydrogen bonding with residues Ala69β/Val70β (1.9-2.0 Å), Asn241β/Asn269β (1.9 Å), and Gln23β/His23β (2.1-2.3 Å), elongation of the scissile bond between the amide nitrogen and the electrophilic carbon of the substrate by 0.1 Å (from 1.4 to 1.5 Å), and reduction of the distance between one of the amine group hydrogens of Ser1β and the substrate’s amide nitrogen, setting up the proton transfer in the second step of the mechanism. The activation barrier connected with TS1 was ca. 7 kcal/mol in all cases except that of the ecPGA-C08-HSL complex, for which the barrier was over 1.2 kcal/mol higher than in the other systems (**Table S13**). This increased barrier height was accompanied by a greater nucleophile attack distance (2.7 ± 0.4 Å) and preferential interaction of the TI with the oxyanion hole stabilizing residue Asn241β over Ala69β (**Table S12**).

**Figure 7.**
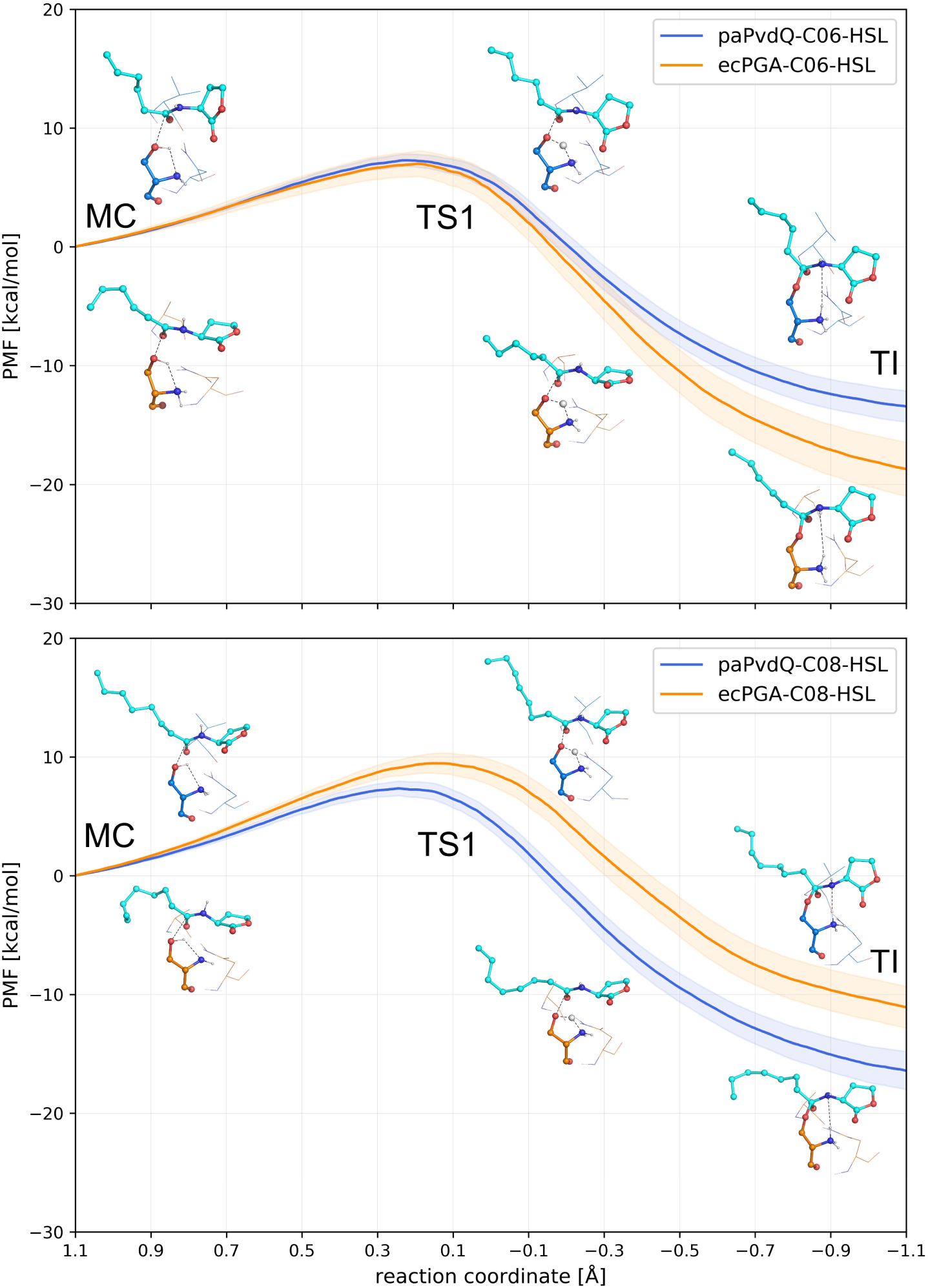
Potential of mean force profiles from steered MD simulations of the tetrahedral intermediate formation step. C06-HSL (upper panel) and C08-HSL (lower panel) acylation catalyzed by the enzymes ecPGA (orange) and paPvdQ (blue). Starting from the MC (RC 1.1 Å), both systems proceed towards TS1 (RC ∼ 0.2 Å) where the proton of the catalytic serine is shared between the residue’s hydroxyl and amine groups while the hydroxyl oxygen engages in nucleophilic attack on the substrate carbonyl carbon, leading to TI formation (RC −1.1 Å).

### In both enzymes, proton transfer initiates the amide bond cleavage leading to acyl-enzyme formation

Continuing from the ensembles of TI states for each protein-HSL complex, we simulated the second step of the acylation process, i.e., the decomposition of the TIs to the corresponding AEs (**Figure 8, Table S12**). During this process, the proton transfer between the amine nitrogen of Ser1β and the amide nitrogen of the leaving group preceded the cleavage of the amide bond, with each stage having distinct energy maxima - TS2a and TS2b **(Figure 8 and Table S14)**. At the first maximum, TS2a, the transferred proton was shared between the serine amine nitrogen and the amide nitrogen of the leaving group but was somewhat closer to the latter (RC ∼ 1.9 Å), and there was a modest elongation of the scissile amide bond from 1.5 Å in the TI to 1.6 Å in TS2a (**Figure S17** and **Table S12**).

**Figure 8.**
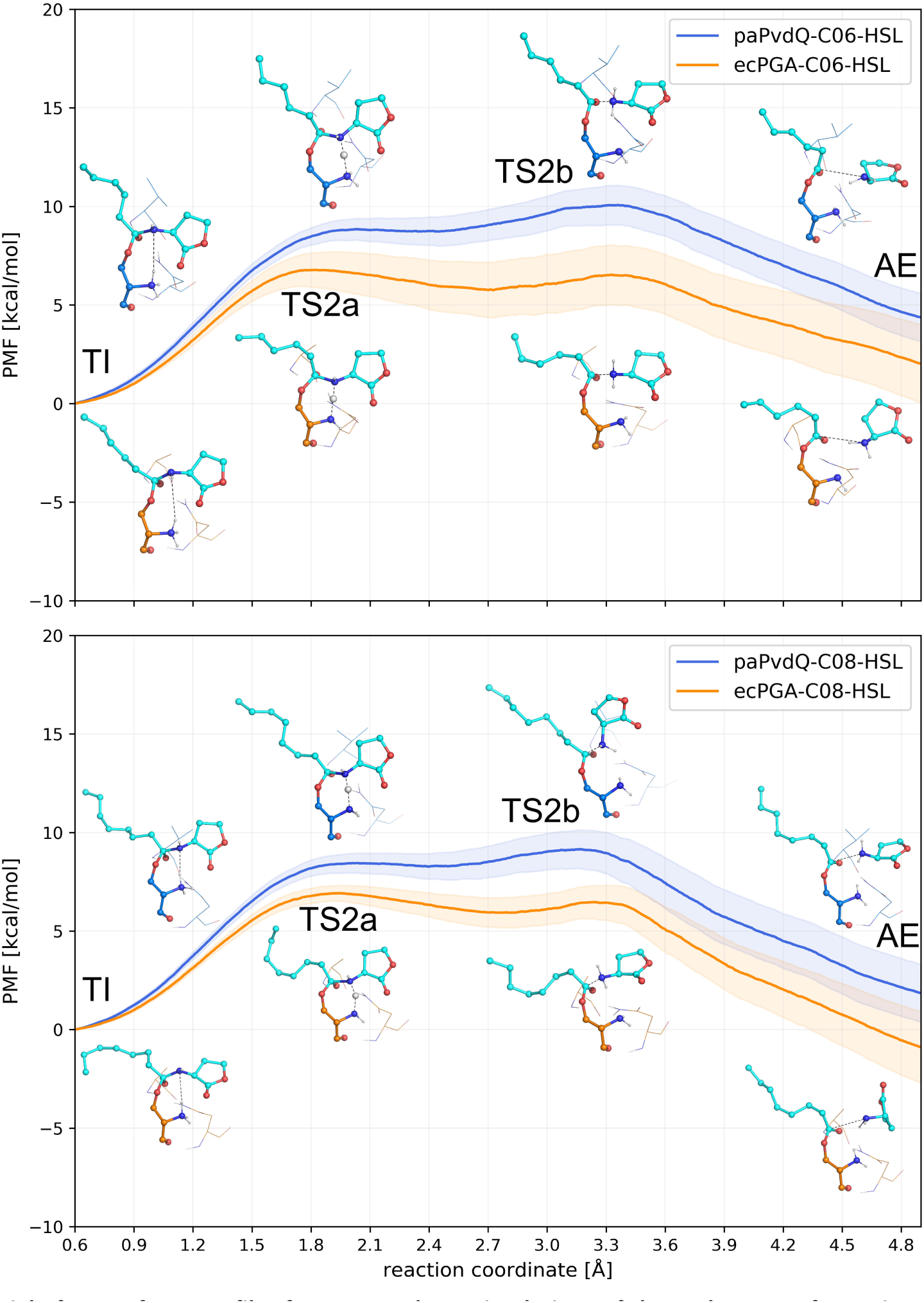
Potential of mean force profiles from steered MD simulations of the acyl-enzyme formation step. C06-HSL (upper panel) and C08-HSL (lower panel) acylation of the enzymes ecPGA (orange) and paPvdQ (blue). Starting from the TI (RC 0.6 Å), the systems progress towards TS2a (RC ∼ 1.9 Å), corresponding to the sharing of the proton between the catalytic serine amine group and the nitrogen of the leaving group. The proton is subsequently fully transferred to the leaving group nitrogen and the amide bond is broken, corresponding to TS2b (RC ∼ 3.3 Å). Finally, the first reaction product, the homoserine lactone, is released from the active site and the AE is formed (RC 4.9 Å).

The elongation of the scissile amide bond became much more pronounced after the proton was fully transferred to the substrate nitrogen (RC ∼ 3.0 Å), proceeding towards full bond cleavage with distances of ≥3.3 Å (**Figure S17** and **Table S12**). The second energy barrier connected with the bond cleavage process (TS2b) coincided with the length of the amide bond being 1.8-2.0 Å (RC ∼ 3.3 Å). The energy maximum connected with bond breaking (TS2b) was lower by up to 0.5 kcal/mol than that connected mainly with the proton transfer (TS2a) for both ecPGA complexes (**Table S14**). In contrast, the breaking of the amide bond (TS2b) was more demanding by 0.8-1.6 kcal/mol for both HSLs in the case of paPvdQ (**Table S14**).

### Experimental assays confirm the computational prediction of ecPGA QQ activity

Driven by encouraging computational predictions, we experimentally evaluated the activity of ecPGA towards C06- and C08-HSLs. First, we examined the spontaneous hydrolysis of C06-HSLs in abiotic conditions at substrate concentrations of 5, 10, and 20 mM at pH 7.0 and 35°C, which is the optimal temperature for ecPGA. The results of these assays were compared to those obtained when samples of C06-HSL were mixed with ecPGA at concentrations of 1.5-250 μM. The hydrolysis of C06-HSL was far more efficient in reaction mixtures containing ecPGA than in those without the enzyme (**Figure 9**). Additionally, the relative abundance of the hydrolyzed substrate increased with the concentration of the enzyme, clearly showing that ecPGA catalyzes HSLs hydrolysis.

**Figure 9.**
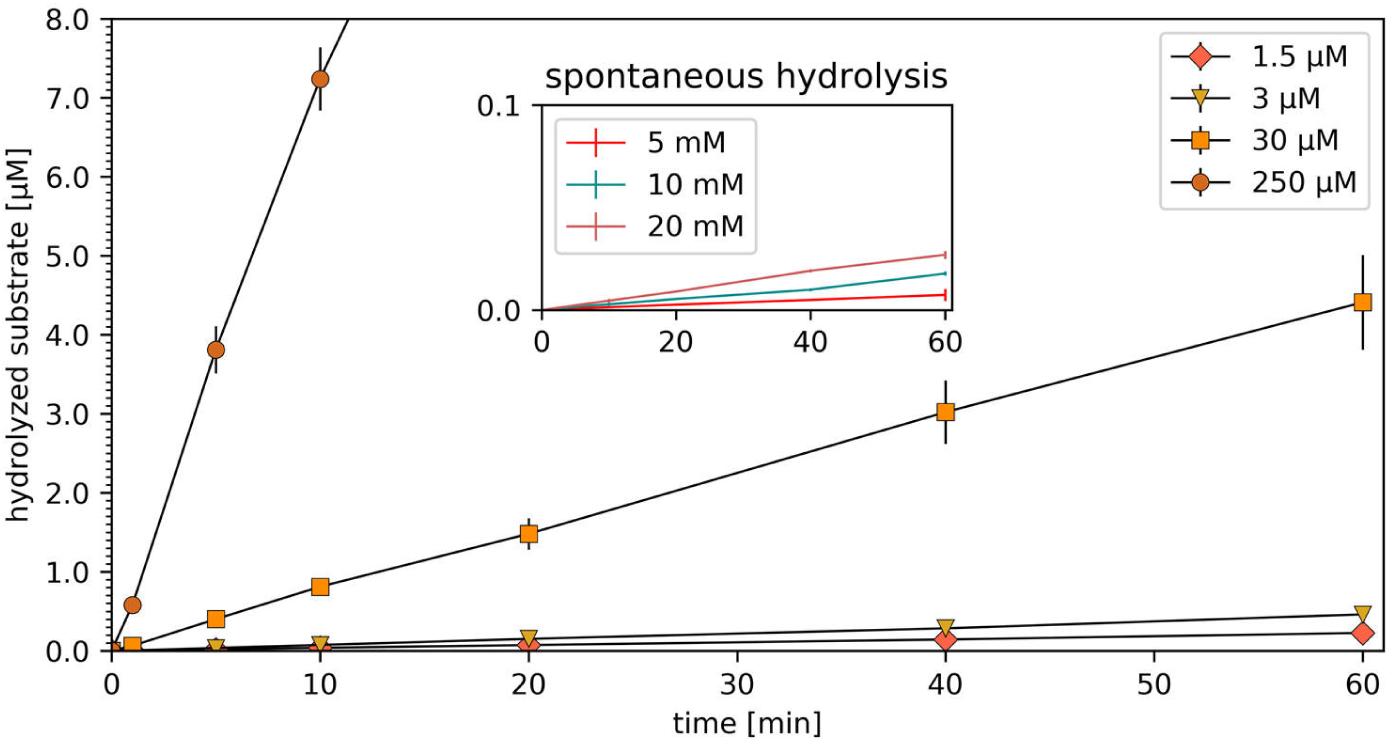
Experimental examination of ecPGA’s catalytic activity. Dependence of the hydrolyzed substrate concentration on the enzyme concentration (main plot), showing that the hydrolyzed substrate concentration increases with the enzyme concentration. The inset shows the rate of spontaneous substrate hydrolysis under the same reaction conditions without the enzyme.

Having verified the activity of ecPGA towards HSLs, we determined the enzyme’s kinetic parameters with both investigated substrates (**Figure S18**). For C06-HSL, the *K*_M_ and *k*_cat_ kinetic constants were 0.59 ± 0.07 mM and 0.0090 ± 0.0003 s^-1^, respectively. The kinetic parameters for the longer substrate, C08-HSL, were less favorable: the *K*_M_ was 0.70 ± 0.04 mM and the *k*_cat_ was 0.0085 ± 0.0003 s^-1^. These results are consistent with the computationally predicted preference for shorter substrates conditioned by less favorable stabilization of the substrate in the productive conformation and greater undesirable fluctuation of the lactone ring and cleaved amide bond.

### Computational and experimental analyses confirm the HSL-degrading activity of remotely related aPGA

Having verified that ecPGA is active against the two chosen HSLs, we probed the QQ activity of a more distantly related member of the PGA family to see how widespread HSL-cleaving activity is among Ntn-hydrolases. For this purpose, we selected aPGA – an enzyme that has been studied extensively in our laboratory, exhibits only around 60 % similarity to both kcPGA and ecPGA **(Table S1)**, and carries a Val56βLeu substitution in its active site.

Free enzyme MD simulations indicated that aPGA efficiently samples the open acyl-binding cavity state, which occurs with a frequency >6% (**Figure S22**). It is thus capable of adopting conformational states that are spatially sufficient to accept HSLs with short or moderately long acyl chains (**Table S15**). Moreover, the entrance to the acyl binding cavity of aPGA was wider than that of both ecPGA and paPvdQ in all three simulations (**Figures S8 and S23)**. In a PCA based on the distances between the functional atoms of the catalytic residues of aPGA, the first two principal components (PC1 and PC2) explained 75% of the total variance. In contrast to the PCA results for the other two enzymes, PC1 depended mainly on the distance between Ser1β and Gln23β/His23β, whereas changes along PC2 depended mainly on the distance between Asn241β/Asn269β and Ala69β/Val70β (**Table S18**), forming an almost inversed picture of the conformational landscape (**Figure S24**). All four of the most prevalent conformational states of aPGA resembled **states a** and **b** of ecPGA and paPvdQ (**Figure S24, Table S16**). In **states a** and **a\***, the serine hydroxyl oxygen was more distant from Gln23β and formed hydrogen bonds with oxyanion hole stabilizing residues, preventing reactive binding of HSLs. Conversely, **states b** and **b\*** were stabilized by hydrogen bonding with the backbone hydrogen of Gln23β, providing a catalytic site organization favorable for productive stabilization of HSLs, although the distance between the oxyanion hole stabilizing residues was higher than in ecPGA and paPvdQ because of the wider opening of the acyl binding site entrance in this enzyme. Consequently, the process of selecting appropriately pre-organized open snapshots yielded a smaller ensemble of snapshots for docking (**Table S15)**. Nevertheless, molecular docking provided complexes of aPGA in which the substrates C06-HSL and C08-HSL were properly stabilized to promote nucleophile attack and which satisfied the previously discussed geometric selection criteria while having favorable binding scores (**Table S17)**.

Using the approach previously applied to ecPGA and paPvdQ, MD simulations of aPGA-HSL complexes were conducted to evaluate this enzyme’s ability to maintain the crucial interactions between the ligand and catalytically relevant residues. As expected, aPGA behaved similarly to ecPGA, meaning that the evaluated distances and angle reached values optimal for catalysis during the simulations, albeit notably less frequently than in paPvdQ (**Figure S25**). Additionally, repeated stabilization of the ligand in a productive state, as shown in **Figure S26**, was more frequently observed for C06-HSL (5 replicates) than for C08-HSL (2 replicates). This reflects the preferential binding of the shorter ligand by aPGA, which was also seen for ecPGA. However, we also observed greater instability of HSLs bound in aPGA; whereas the enzyme-substrate complexes of ecPGA were highly stable (**Figures S12 and S13**), the substrate often dissociated fully from the active site of aPGA (**Figure S27**). An analysis of the mobility of the HSL heavy atoms indicated that the most mobile part of the ligand in the aPGA complexes was the lactone ring, followed by the atoms of the amide bond and the acyl chain (**Figure S28**). The amplitudes of the heavy atom fluctuations of C08-HSL were similar to those observed for the ecPGA complexes, but most of the atoms of C06-HSL exhibited significantly larger fluctuations in aPGA, enabled by the wider opening of this enzyme’s acyl-binding cavity. Finally, binding free energy analysis showed that aPGA bound HSLs as tightly as ecPGA, both in terms of the absolute energy (**Table S19**) and the individual contributions of each residue (**Figure S29**).

When using steered QM/MM MD simulations to study the acylation reactions of aPGA, we had to relax the selection criteria for the aPGA-C08-HSL structures because optimally stabilized representatives were not adequately sampled during the simulations (**Table S20**). However, the selected structures were sufficiently close to the desired parameters to enable their adjustment during restrained MD simulations and to thereby generate an appropriate ensemble of MCs as starting positions for QM/MM MD simulations (**Table S21**).

The reaction mechanisms observed for aPGA with C06-HSL and C08-HSL during the TI and AE formation steps proceeded through TS1, TS2a, and TS2b in a manner consistent with the mechanisms observed for ecPGA and paPvdQ (**Table S22** and **Figures S30-S33**). Curiously, the TS1 energy barrier for aPGA-C06-HSL (**Table S23**) was similar to that for ecPGA-C08-HSL and was substantially higher than the barriers for the other complexes in this reaction step (**Table S13** and **S23**). As in ecPGA-C08-HSL, the nucleophilic attack by Ser1β was delayed in aPGA-C06-HSL (**Figure S32**), increasing the nucleophile-electrophile distance in TS1. However, in this substrate the carbonyl oxygen was closer to Ala69β than to Asn241β (**Table S22)**.

Surprisingly, the energy barrier for the second step of the reaction with C08-HSL for aPGA was 9.0 ± 0.7 kcal/mol in TS2a (**Table S24**), which was much higher than the barrier seen for ecPGA and comparable to that for paPvdQ in this step (**Table S14**). Nonetheless, the overall energetic cost of traversing the RCs in aPGA was comparable to those for the other two enzymes, with TS2a and TS2b being located at similar points along the RC, suggesting that aPGA should also be able to hydrolyze C06-and C08-HSLs. To validate these findings, we determined the kinetic parameters of aPGA with both substrates. The observed *K*_M_ values of 0.62 ± 0.1 mM (C06-HSL) and 0.87 ± 0.01 mM (C08-HSL) were higher than those for ecPGA, in keeping with the computational observation of less frequent productive stabilization of substrates by aPGA. The catalytic constants of aPGA for C06- and C08-HSLs were 0.0079 ± 0.0003 s^-1^ and 0.0064 ± 0.0001 s^-1^, respectively (**Figure S19**).

### Dynamic determinants of quorum quenching activity in N-terminal hydrolases

Intensive computational investigations and experimental studies showed that all three of the chosen representatives of the N-terminal serine hydrolase family – aPGA, ecPGA, and paPvdQ – exhibit appreciable activity towards C06- and C08-HSLs. Importantly, we observed system-dependent conformational and energetic preferences governed by several dynamic determinants that were primarily connected to the behavior of two molecular gates, namely (i) gates to the acyl-binding cavity, and (ii) gates controlling the overall accessibility of the active site.

The barriers to the first acylation step for the aPGA-C06-HSL and ecPGA-C08-HSL complexes were higher than those for the other studied systems. In both cases, the increased barrier height was attributed to suboptimal oxyanion hole stabilization, although with a different shift from the optimum. As illustrated in **Figure 10A**, in the aPGA complexes the substrate C06-HSL tended to interact more strongly with the oxyanion hole stabilizing residue Ala69β than with the second oxyanion hole residue (Asn241β) because Ala69β sampled catalytically suboptimal side-chain conformations more frequently than in the other C06 complexes. These conformations corresponded to **state b\*** identified in the PCA analysis, in which the interaction between the oxyanion stabilizing residues is catalytically suboptimal (**Figure S24**). Ala69β is situated in close proximity to the acyl-binding site entrance, which opens more widely in aPGA than in the other two enzymes (**Figure S23**). Contrasting behavior was seen in the ecPGA-C08-HSL complex, in which oxyanion hole stabilization is primarily performed by Asn241β (**Figure 10B**), which is located closer to the bulk solvent. In both cases, the deviation from the optimal oxyanion hole stabilization causes the substrate molecule to be positioned further from the nucleophile in both the MCs and TS1 (**Table S12 and S22**), delaying covalent bond formation (**Figures S16 and S32**) and increasing the energy barriers of the reaction (**Figures 7 and S30**). In the remaining systems (**Figure 10C-D**), TS1 features balanced stabilizing contributions from both oxyanion hole stabilizing residues, shortening the distance from the nucleophile to the electrophile and reducing the reaction’s energetic barrier.

**Figure 10.**
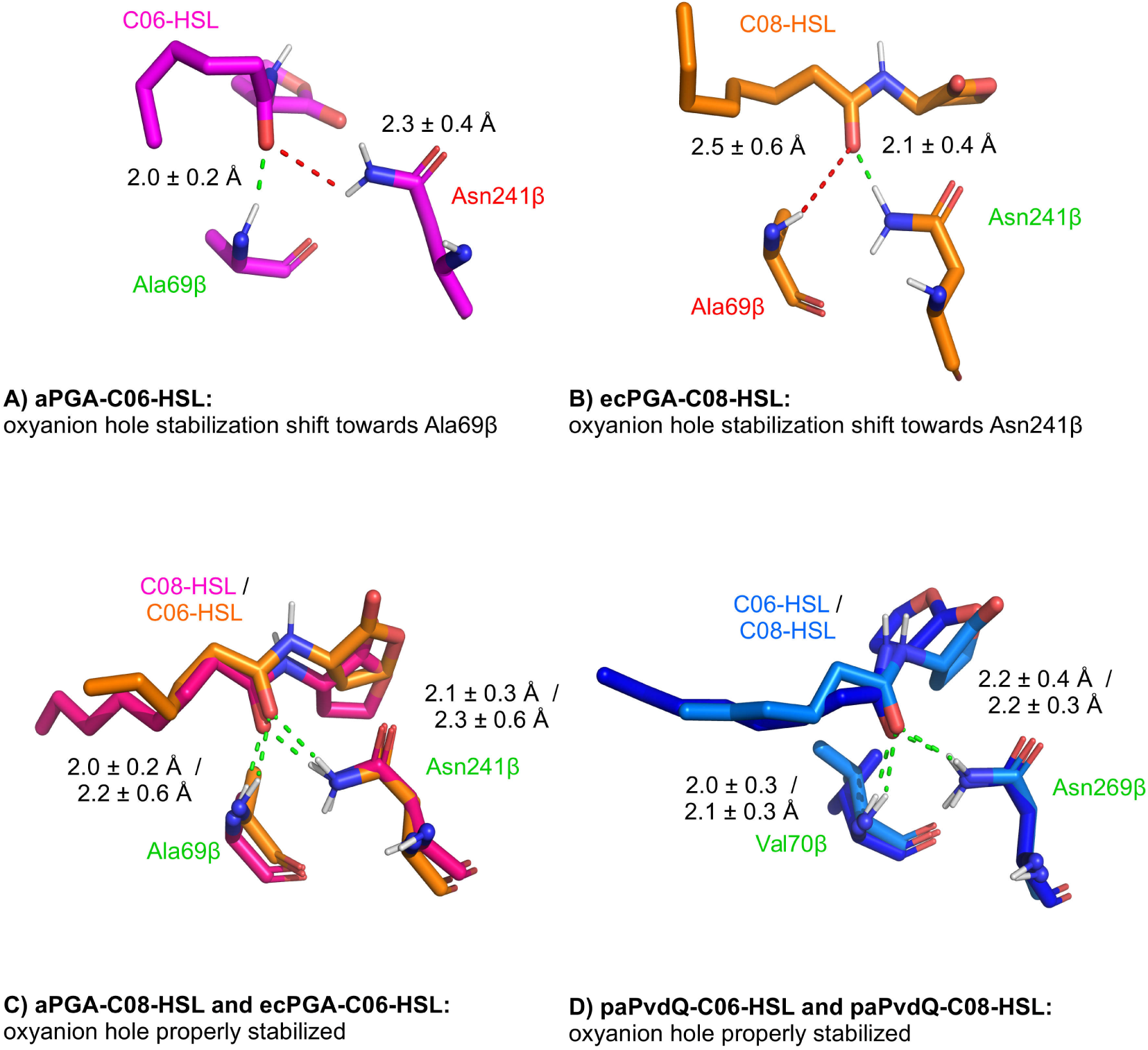
Differences in oxyanion hole stabilization in TS1 contributing to increased energy barriers for the aPGA-C06 and ecPGA-C08 complexes. A) the aPGA-C06-HSL complex, in which the HSL carbonyl oxygen is shifted towards Ala69β, B) the ecPGA-C08-HSL complex with the HSL carbonyl oxygen preferentially stabilized by Asn241β. C) complexes of PGAs with properly stabilized oxyanion holes and D) similar oxyanion hole stabilization in both paPvdQ complexes.

To understand the difference between the aPGA and ecPGA enzymes, we focused on sequential differences between the residues in the vicinity of the catalytic machinery and binding cavity, because the catalytic machinery consists of the same amino acids in both enzymes. There are two differences in this region: (i) aPGA carries a bulky leucine residue in position 56β, which is replaced by the less bulky valine in ecPGA, and (ii) ecPGA contains a tyrosine in position 27β where aPGA has a tryptophan residue. The residue at position 56β influences the depth of the acyl-binding cavity, which is slightly longer in ecPGA. The variability at position 27β affects the dynamics of the gating residue Phe24β, whose backbone is stabilized by hydrogen bonding with the hydroxyl group of the tyrosine side-chain in ecPGA but is free to move in aPGA because the steric hindrance imposed by the hydroxyl group is missing (**Figure S34 and S35**). The increased mobility of the gate in aPGA is consistent with the more pronounced opening of its acyl-binding site (**Figure S23**). Curiously, the availability of additional space resulted in favorable energetics for longer substrates but had an undesired effect for C06-HSL, which bound too deeply in the cavity and was thus stabilized by a less catalytically productive conformation of Ala69β (**Figure 10A**). No such opening of the acyl-binding site is possible in ecPGA because of the efficient stabilization of Phe24β by Tyr27β (**Figure S35**), which is favorable for shorter substrates but less suited for C08-HSL because the acyl binding cavity lacks the space to properly accommodate the longer acyl chain. On the other hand, the differently composed gate of paPvdQ exhibited relatively steady behavior in complexes with both ligands, resulting in a deep and stable acyl-binding cavity (**Figure S36**). Furthermore, while inspecting the geometries of the HSLs in TS1 we observed a clear trend for the HSL acyl chain to bend in aPGA to fit the broader cavity, which was not seen in the other two enzymes (**Figure S37A**).

As mentioned in the previous sections, the energy barriers also varied for the second reaction step. This variation could be traced to differences in the arrangement of Arg263β in the PGAs and Arg297β in paPvdQ with respect to the amine group of Ser1β and the leaving homoserine lactone oxygen (**Figure 11, Table S12** and **S22**). These arginine residues are known to play crucial roles in the acylation step catalyzed by ecPGA and are believed to be important for catalytic activity in paPvdQ as well.^48,69,90^ In paPvdQ, Arg297β was closer to the serine amine group and almost aligned with the direction of proton transfer, rendering the substrate’s lactone oxygen inaccessible. On the other hand, in PGAs, Arg263β adopted a different conformation further from the Ser1β amine group due to a cation-pi interaction with Trp240β, which is absent in paPvdQ **(Figure 11**). This rearrangement gave Arg263β access to the lactone oxygen of HSL, enabling additional stabilization of the leaving group.

**Figure 11.**
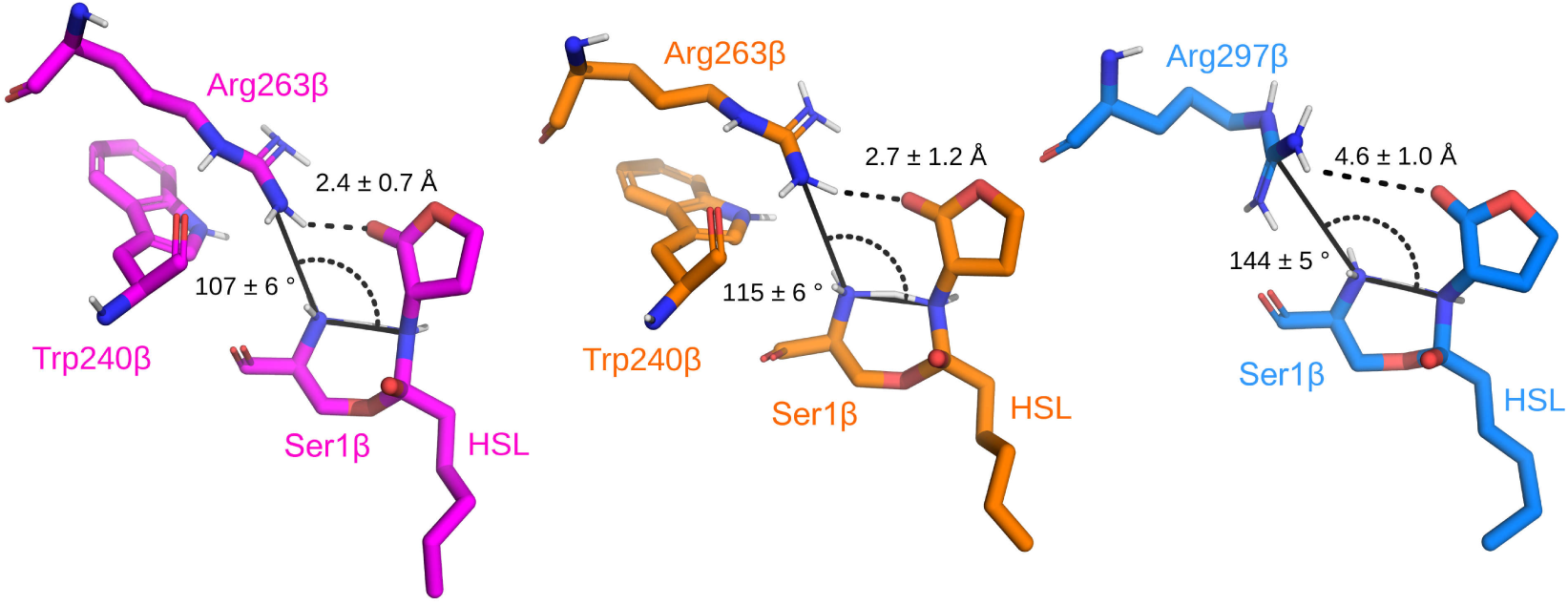
Different stabilization of TS2a by Arg263β in PGAs and Arg297β in paPvdQ. The arginine was, on average, closer to the Ser1β amine group and further from the lactone oxygen of HSL in paPvdQ (in blue). The opposite was true in aPGA (in magenta) and ecPGA (in orange). The angle between the HSL nitrogen, the serine amine nitrogen, and the arginine guanidyl carbon was higher for paPvdQ, blocking the access of the arginine to the lactone. Also note that the plane of the guanidyl group in the PGAs is roughly perpendicular to that in paPvdQ.

Additionally, the energy barriers connected with both TS2a and TS2b were higher for aPGA-C08-HSL than for the other PGA complexes. Interestingly, this complex had a higher tendency to sample the closed conformational state of the whole active site, which was characterized by interactions between Arg145α and Phe24β (state **C** in **Figure S38**).^53,91^ In this closed state, Arg145α is located in the vicinity of the substrate’s scissile amide bond (ca. 7.5 Å) and can thus interfere with the proton transfer. In aPGA, the formation of a closed state was facilitated by Trp27β, which leaves the backbone oxygen of Phe24β more exposed to interactions with the guanidine group of Arg145α.

Furthermore, the bent acyl chain of C08-HSL (**Figure S37B**) in the aPGA complex, which was not seen in the other systems at this reaction stage, interacted preferentially with Phe146α, leaving space for Phe24β to adopt an inclined conformation able to interact with Arg145α (seen in the semi-closed **C/O** and closed **C** states shown in **Figure S38**); this promotes the formation of the closed state by aPGA. In ecPGA, the backbone oxygen of Phe24β forms a hydrogen bond with the hydroxyl group of Tyr27β, effectively blocking its interaction with Arg145α and the adoption of the closed state (**Figure S39**). This agrees with the findings of Alkema and Novikov, who showed that Arg145α influences substrate specificity.^53,90^

## DISCUSSION

QQ enzymes have undoubtedly attracted considerable interest over the last decade, and many new enzymes have been shown to exhibit activity against various QS signaling molecules. These results have expanded our knowledge of QQ and moved us closer to practical QQ-based antibacterial strategies that could be used to address the problems of widespread antibiotic resistance and biofouling. Although many QQ enzymes are known, as shown in several recent comprehensive reviews,^47,92^ some crucial challenges remain unsolved. Most of the characterized enzymes with high QQ activity have not been optimized for large-scale use. In contrast, robust enzymes that are well established in industrial catalysis often exhibit relatively low catalytic rates or even a complete lack of activity towards QS signaling molecules, or may have problematic substrate specificities. This highlights the need for further work to elucidate the molecular determinants of efficient QQ activity to enable their effective transplantation into biotechnologically optimized enzymes.

Only a few investigations seeking to determine the molecular mode of action of the N-terminal serine hydrolases studied in this work have been presented. A catalytic mechanism for paPvdQ was initially proposed based on the crystallographic data for the apo-enzyme,^48^ and was further extended by work using transition state analogues^56^ and mutations that shifted its substrate specificity towards C08-HSL.^49^ Our results confirm that paPvdQ has a deep acyl-binding pocket that remained stable over the timescale of our simulations, whereas PGAs have significantly shallower pockets. We verified the crucial importance of the residues forming catalytic machinery alongside Ser1β, including the oxyanion hole residues Val70β and Asn269β and the additional stabilizing residue His23β, whose importance was shown in previous studies.^48,49^ The results obtained showed that the catalytic Ser1β is capable of self-activation through the transfer of the hydroxyl proton to its N-terminal amine group without needing a bridging water molecule. This finding is consistent with the observations of Clevenger *et al*. during experiments using transition state analogues with paPvdQ and with the computational analysis of ecPGA complexed with penicillin G reported by Grigorenko *et al*.,^56,69^ suggesting that this internal activation process may occur in N-terminal serine hydrolases in general, independently of the substrate molecule. Our analysis of the dynamics of the chosen enzyme-substrate complexes showed that the highly conserved residue Arg297β^48,56^ tends to adopt different conformations to the analogous residue Arg263β in PGAs, which raises the energetic barriers in the second step of the reaction for paPvdQ. The substrate specificity of paPvdQ towards four different QS signaling molecules (C12-HSL, 3-oxo-C12-HSL, C08-HSL and C04-HSL) was also investigated using MD simulations, which were often started from binding poses in which the substrate molecules were rather distant from the catalytic residues.^93^ As a consequence, these simulations only explored catalytically competent states of paPvdQ infrequently (and sometimes, not at all) within the chosen timeframe of 300 ns, as illustrated by the absence of productive binding modes in the C08-HSL-paPvdQ complexes. When using productive binding poses to initiate simulations, we found that paPvdQ has a substantial capacity to stabilize and catalyze the conversion of not only C08-HSL but also C06-HSL, both of which are expected to be productively bound and converted by this enzyme.^49,56,94^ Based on these complexes, we investigated the molecular details of this enzyme’s catalytic action.

The only previously published report confirming QQ activity among PGAs was presented by Mukherji *et al*., who experimentally verified the QQ ability of kcPGA and established its preference for 3-oxo-C06-HSL.^51^ Otherwise, studies on the structure-dynamics-function relationships of the N-terminal serine hydrolases have mainly focused on ecPGA with its native substrates or related compounds due to the importance of this enzyme for the production of semi-synthetic antibiotics. These studies have used both experimental techniques and a broad spectrum of molecular modeling methods including docking, MD simulations, and QM or QM/MM calculations.^53,57,69,70,85,95^ Our study provides a complementary extension of the current knowledge of N-terminal serine hydrolases and particularly their QQ activity based on extensive QM/MM MD simulations. To our knowledge, comparable simulations have only been reported for cysteine N-terminal-hydrolases – closely related enzymes that behave differently to serine or threonine hydrolases because the catalytic cysteine preferentially adopts a zwitterionic form,^96,97^ making an entirely analogous mechanism unlikely.

We observed that ecPGA, aPGA, and paPvdQ can all accommodate and bind HSLs, stabilize HSLs in productive conformation, and catalyze their degradation via similar reaction mechanisms. These mechanisms involve geometrically comparable states in each reaction step that are consistent with the established mechanism for ecPGA with its native substrate–penicillin G.^69^ The energy barriers obtained from hundreds of replicated steered MD simulations for all investigated systems were around 10 kcal/mol or below, which is typical for acylation reactions of this kind, in particular when considering the tendency of the semi-empirical PM6-D method to underestimate them.^98^ For example, Nutho *et al*. found that the barriers to the acylation reaction catalyzed by Zika virus serine protease according to this method were ca. 8 kcal/mol lower than those suggested by high-level QM/MM.^99^ Taking this underestimation into account would significantly reduce the predicted acylation rates for both enzymes, resulting in worse rates than those achieved with their native substrates,^49,69^ as expected for non-primary substrates. Our computational results were supported by experiments that proved the activity of ecPGA and aPGA towards both C06-HSL and C08-HSL and whose results were consistent with the reported substrate preferences of paPvdQ.^49^ The calculated catalytic efficiencies of ecPGA and aPGA for the studied substrates yielded *k*_cat_/*K*_M_ values of around 0.01 mM^-1^ s^-1^, which is around an order of magnitude lower than that for kcPGA (0.11 mM^-1^ s^-1^),^51^ suggesting that although both enzymes exhibit basal activity towards short- and medium-length HSLs, kcPGA remains the most efficient option.

Importantly, our study indicates that paPvdQ stabilizes HSLs in productive binding conformations more efficiently than PGAs. This can be attributed to its narrow and relatively static acyl-binding cavity, which efficiently limits the mobility of the cleaved amide bond. Conversely, the dynamics of the gating residues of the acyl-binding cavity of PGAs allow these enzymes to accommodate a much broader range of substrates including HSL, various amino acids, and penicillins.^51,54,85,87^ Our QM/MM MD-based study revealed protein- and ligand-dependent differences in the dynamics of these gating residues, showing that complexes of aPGA with longer HSL substrates were more likely to sample closed conformations than the other PGA complexes included in the study. This observation agrees well with insights from previous studies suggesting a preference for closed states in the free enzyme and complexes with specifically recognized ligands, whereas open states are preferred for complexes with nonspecific ligands.^53,91^ Consequently, these gates and their structural proximity represent attractive engineering targets for future studies aiming to develop PGA variants tailored to specific bacterial QS molecules.

In summary, this study of three enzymes belonging to two distinct N-terminal serine hydrolase sub-families – the penicillin G acylases and acyl-homoserine lactone acylases – has revealed common mechanisms of QQ activity towards HSLs used in bacterial signaling. Members of this family degrade QS compounds via the same reaction mechanism, but the efficiency of this degradation reaction depends on several enzyme- and substrate-specific determinants. As such, the obtained results expand our understanding of the catalytic mechanisms of N-terminal serine hydrolases at the molecular level, which to our knowledge have not previously been investigated for any of the enzymes included in this study. Furthermore, we have added two new members to the set of known QQ enzymes, both of which are well-optimized and widely used in industrial catalysis: aPGA and ecPGA.^100–102^ Finally, by performing a detailed comparison of the structure-dynamics-function relationships of this QQ activity in paPvdQ and PGAs, we have identified structural and dynamic features of the PGAs that explain their relatively low activity in individual catalytic steps, providing further direction for future efforts to develop more potent antibacterial agents.

## ASSOCIATED CONTENT

Multiple sequence alignment of investigated serine Ntn hydrolase family members; sequence similarity matrix for investigated serine Ntn hydrolase family members; detailed descriptions of computational protocols; details of the evolution of the backbone RMSD plots for MD simulations; time evolution of the entrance bottleneck to the acyl-binding site; acyl-binding site dynamics; characterization of PCA states; detailed characterization of docked HSL poses; visualizations of ligands in productive conformations throughout the MD trajectories; ligand RMSD plots for MD simulations; per-residue decompositions of MM/GBSA binding energies; characterization of favorable organizations of protein-HSL complexes suitable for the amide cleavage reaction; a description of the QM/MM MD sampling process; characterization of the active site geometries in specific stages of the reaction based on the QM/MM MD results; energetics of the acylation reaction steps; evolution of RC components during the acylation reaction; experimental measurements of the activity of the aPGA and ecPGA enzymes at different substrate concentrations; definitions of differences in gating dynamics in the studied systems; distributions of ligand bending in different stages of the reaction; and information on the different open/closed state preferences of aPGA and ecPGA enzymes as well as their dependence on the bound ligand (PDF)

## AUTHOR CONTRIBUTIONS

B.S. performed the computational analyses; M.G. performed experimental characterization of ecPGA and aPGA; A.P. and H.M. executed all the bacterial flask and fed-batch cultivation in a bioreactor to produce enzymes; J.B. designed the project, analyzed and interpreted data. The manuscript was written through the contributions of all authors. All authors have approved the final version of the manuscript.

## ACKNOWLEDGMENT

This work was supported by the National Science Centre, Poland (grant number 2017/25/B/NZ1/01307) and by the Institutional Research project RVO61388971 from the Institute of Microbiology of the CAS. B.S. is a recipient of a scholarship provided by POWER project POWR.03.02.00-00-I022/16. The computations were performed at the Poznan Supercomputing and Networking Center.

## ABBREVIATIONS

QS: quorum sensing
QQ: quorum quenching
HSL: N-acyl-homoserine lactone
QSI: quorum sensing inhibitor
paPvdQ: *Pseudomonas aeruginosa* acyl-homoserine lactone acylase
PGA: penicillin G acylase
kcPGA: *Kluyvera citrophila* PGA
ecPGA: *Escherichia coli* PGA
aPGA: *Achromobacter spp*. PGA
MD: molecular dynamics
RMSD: root-mean-square deviation
PCA: principal component analysis
MM/GBSA: Molecular Mechanics / Generalized Born Surface Area
QM/MM MD: Quantum Mechanics / Molecular Mechanics MD simulation
NPT: isothermal-isobaric ensemble
TI: tetrahedral intermediate
AE: acyl-enzyme
RC: reaction coordinate
LCOD: linear combination of distances
PMF: potential of mean force
HPLC: high-performance liquid chromatography
PC: principal component
RMSF: root-mean-square fluctuation
MC: Michaelis complex
TS1: transition state 1
TS2a: transition state 2a
TS2b: transition state 2b

## Notes

### Competing Interest Statement

The authors have declared no competing interest.

### Summary of Updates

only minor edits to improve clarity

