## Supplementary information for "Common Dynamic Determinants Govern Quorum Quenching Activity in N-terminal Serine Hydrolases"

---

**KEYWORDS**

Quorum quenching, N-terminal serine hydrolase, acyl-homoserine lactone acylase, penicillin G acylase, QM/MM, molecular dynamics, reaction mechanism

Tables:

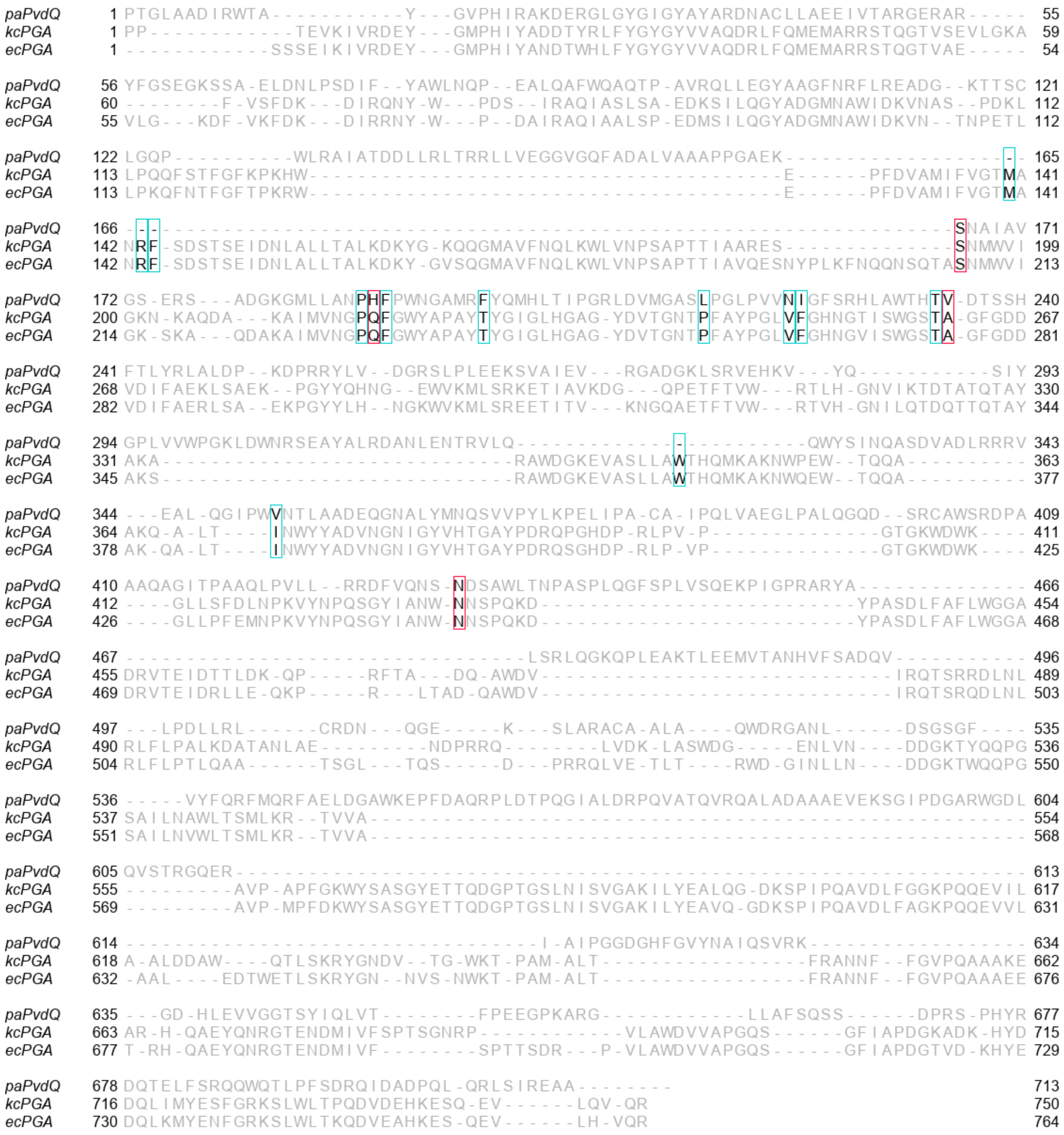

**Figure S1.** Multiple sequence alignment of paPvdQ, kcPGA, and ecPGA enzymes. Active site and catalytic residues are highlighted by cyan and red frames, respectively. Multiple sequence alignment (MSA) was prepared using Clustal Omega 1.2.4. The sequence-structure relationships between proteins were examined by multiple sequence alignment followed by an iterative structural superposition in PyMOL. MSA was visualized using Jalview 2.7.

**Table S1.** Sequence similarity matrix for aPGA, ecPGA, kcPGA, and paPvdQ enzymes generated with SIAS web-server (http://imed.med.ucm.es/Tools/sias.html).

| | | | | |
|---|---|---|---|---|
| aPGA | 100% | 62% | 64% | 29% |
| ecPGA | 62% | 100% | 90% | 26% |
| kcPGA | 64% | 90% | 100% | 26% |
| paPvdQ | 29% | 26% | 26% | 100% |
| protein | aPGA | ecPGA | kcPGA | paPvdQ |

### DETAILED COMPUTATIONAL PROTOCOL

**Molecular dynamics of free enzymes.** Crystal structures of ecPGA (PDB-ID: 1GK9) and paPvdQ (PDB-ID: 4M1J) were obtained from the PDB database.[1,2] For *Achromobacter spp.* penicillin G acylase (aPGA), a previously derived homology model was retrieved from the Protein Model Database (ID: PM0080082)[3] and corrected using the RepairPDB module of FoldX.[4] Structures were protonated with the H++ webserver[5–7] at pH 7.5 using default salinity (0.15) and internal and external dielectric constants (10 and 80, respectively). Protonation states were checked in detail visually in PyMOL and VMD 1.9.2[8] to preserve the functional protonation state of the active site residues for all proteins in the study. The N-terminal serine (Ser1β) was assumed to be in the neutral protonation state ($NH_2$-Ser-OH) according to the nucleophile attack reaction mechanism as described elsewhere.[9–13] Partial charges of the N-terminal serine in its neutral form were obtained at the HF/6-31G* level of theory using Gaussian09.[14] Further, the charges were derived using RESP-A1 charge model by multi-conformational RESP[15] fit employing RESP ESP web-sever.[16,17] The atom types were derived analogously to Cornell et al. force field.[18] Protonated structures including crystallographic waters were placed in a truncated octahedral box of TIP3P waters whose boundaries were at least 10 Å away from all atoms in the structure and neutralized using Na+ and Cl- ions to approximately 0.1 M concentration. Initial parameters and topologies were generated using tleap module of AmberTools17.[19] For paPvdQ structure, three known disulfide bridges were explicitly set up (Cys44α → Cys125α, Cys217β → Cys237β, and Cys339β

→ Cys352β),[20] which are not present in any of the investigated PGAs. Using Parmed module of AmberTools17, hydrogen atom masses were repartitioned to enable 4 fs time-step during simulations using the SHAKE algorithm.[21,22] Energy minimization and molecular dynamics (MD) simulations were performed using the ff14SB force field[23] with the pmemd and pmemd.CUDA modules of the Amber16 package, respectively.[19] The systems were energy minimized by 1000 steepest descent steps followed by 1000 conjugate gradient steps in five stages with decreasing positional restraints. In the first stage, the protein heavy atoms were restrained with a 500 kcal $mol^{-1}$ $Å^{-2}$ force constant. In the following four minimization steps, protein backbone heavy atoms were restrained using 500, 125, 25, and 0.001 kcal $mol^{-1}$ $Å^{-2}$ harmonic restraints. Subsequently, systems were equilibrated in four steps, starting with 100 ps NVT simulation at 100 K using Langevin thermostat[24] with protein heavy atoms harmonically restrained with 5 kcal $mol^{-1}Å^{-2}$ force constant, followed by three NPT stages of dynamics with Langevin thermostat and Monte Carlo (MC) barostat: (i) 300 ps in 100 K with protein heavy atoms harmonically restrained using 5 kcal $mol^{-1}$ $Å^{-2}$ force constant, (ii) 600 ps with gradual heating from 100 K to 310 K during first 200 ps and backbone heavy atoms restraint of 5 kcal $mol^{-1}$ $Å^{-2}$ and (iii) 1 ns at 310 K without positional restraints. Finally, 500 ns NPT production MD simulation was performed at 310 K was performed using the Langevin thermostat and by controlling the pressure using the MC barostat. The trajectory was generated by saving coordinates every 20 ps during simulation. All steps were performed in triplicate to generate three independent replicates. The stability of the production phase was evaluated in terms of the root-mean-square deviation (RMSD) of the backbone heavy atoms (**Figure S3 and S20**).

**Free enzyme dynamics analysis.** The opening of the protein binding cavity across free enzyme MD trajectories was investigated using the CAVER 3.0.2 program.[25] The starting point for the calculation of paths was chosen based on the center of mass of three residues: Met142α, Ser67β, and Ile177β for PGAs, and Leu146α, Leu53β, and Trp162β for paPvdQ,[26] constraining the maximum starting point distance from the computed point as 1 Å. Further, potential binding site opening events were identified using a probe radius of 0.5 Å. A time sparsity of 10 was used to reduce the computational cost of this analysis. The paths were clustered using a threshold of 3.5, and one representative path was selected from each cluster. Cluster corresponding to binding cavity was selected based on visual examination in PyMOL and VMD. Time evolution of the binding cavity opening was examined by analyzing the entrance narrowest point of the CAVER cluster starting from the protein surface. The adequate opening was defined when the entrance bottleneck exceeded 1.4 Å.[27,28] Further, spatially well-shaped cavities were identified using the following criteria: (i) the minimum depth of the binding site should be at least 5 Å, (ii) the minimum width of the entrance to the binding site should be greater than 1.4 Å, and (iii) the maximum distance between real and defined starting- and end-points for tunnel calculations to be lower than 1 Å.

The cpptraj module of AmberTools17 was used to measure relevant distances between functional atoms in the catalytic residues across all MD trajectories. All distances considered in this work relate to internuclear distances. In addition, the network of distances was subjected to dimensionality reduction using principal component analysis (PCA) method as implemented in the *decomposition* module of the scikit-learn Python library.[29] The PCA was performed combining information from three replicates for each protein separately. The first two components were visualized as density maps using Python seaborn library.[30] Representative structures of the most sampled regions were extracted from corresponding trajectories based on the information from density maps by searching for the four highest density peaks using the *peak_local_max* method from the *feature* module of Python scikit-image library.[31]

**Receptor selection and molecular docking**. Protein conformations harboring spatially well-shaped binding cavities and favorably pre-organized catalytic machinery were selected as receptor structures for ligand docking according to the criteria presented in **Table S2** (*"receptor selection for docking"*). These criteria were used to promote proper orientation of the hydroxyl hydrogen of the nucleophilic serine relative to its amine group, which serves as a hydrogen acceptor in the first reaction step and to ensure appropriate pre-organization of the catalytic machinery (i.e., the nucleophile serine and the oxyanion hole residues) for reactive binding of the substrate.[3,13,32,33] Structures of C06-HSL and C08-HSL ligands were created in Avogadro,[34] partial charges were obtained at HF/6-31G* level of theory using Gaussian09.[14] Further, the charges were derived using RESP-A1 charge model by multi-conformational RESP[15] fit employing the RESP ESP web-sever.[16,17] The ligand structures with updated charges were used to generate input pdbqt files for docking using prepare_ligand.py script from AutoDockTools4. For molecular dynamics simulations with Amber, the ligand atom types were derived analogously to Wang et al. force field.[35] Molecular docking was performed using Autodock4.2.6[36], using 250000 maximum number of energy evaluations, 1000 dockings per Lamarckian genetic algorithm engine search, with the varying number of individuals in the population (50, 75 and 100). Receptor and ligand structures were prepared using prepare_receptor4.py and prepare_ligand4.py scripts from AutoDockTools4,[36] respectively. The searching space was reduced by selecting a grid box covering the binding site of the protein with the initial positioning of the center of the box calculated based on the center of mass of residues: Met142α, Ser67β, Ile177β, Ser1β, Ala69β, and Asn241β for PGAs, and Leu146α, Leu53β, Trp162β, Ser1β, Val70β, and Asn269β for paPvdQ. Further, the center and the size of the box were adjusted manually using Autodock/vina PyMOL plug-in,[37] based on visual examination to cover the entire binding cavity.[36] Representative protein-ligand complexes were selected for the following round of simulations based on the favorability of their Autodock binding scores and the mechanism-based selection criteria,[38] as listed in **Table S2** ("*selection of complexes for MDs*") which were chosen to ensure an acceptable distance between the nucleophile and electrophile as well as proper stabilization of the oxyanion hole.

**Table S2.** Distance criteria for particular parts of the modeling.

| stage/label[id] | *SerO →HSL-C[1] | **SerO >HSL-C> HSL-O[2] | *Ala/ValNH →HSL-O[3] | *AsnNH2 →HSL-O[4] | *Gln/HisO →HSL-H[5] | *SerH →SerN[6] | *SerNH1 →SerO[7] | *SerNH2 →SerO[8] | *AsnN→ Ala/ValN[9] | *Ala/Val N→SerO[10] | *AsnN →SerO[11] | *ArgN →HSL-O[12] | *SerN→ AHL-N[13] |
|---|---|---|---|---|---|---|---|---|---|---|---|---|---|
| receptor selection for docking | - | - | - | - | - | <3.0 | - | - | <5.0 | <5.0 | <5.0 | - | - |
| selection of complexes for MDs | <3.3 | - | <3.0 | <3.0 | - | - | - | - | - | - | - | - | - |
| restraints for complexes MDs | 3.3 | - | 3.3 | 3.3 | - | - | - | - | - | - | - | <4.0 | - |
| selection of representatives for sMDs | 3.0 | 90 | <3.0 | <3.0 | <3.0 | <3.0 | <3.0 | <3.0 | - | - | - | - | - |
| restraints for sMDs inputs generation | 3.0 | - | <3.0 | <3.0 | <3.0 | - | <2.5 | <2.5 | - | - | - | - | - |
| restraints for 1st QM/MM equilibration | 3.0 | - | <3.0 | <3.0 | <3.0 | - | <2.5 | <2.5 | - | - | - | - | - |
| restraints for 2nd QM/MM equilibration | 1.5 | - | - | - | - | - | - | - | - | - | - | - | 2.0 |

*distance provided in [Å]; **angle provided in [°]; Note: detailed description of particular criteria is available below

| id | description | atoms involved |
|---|---|---|
| **1** | nucleophile distance | Ser1β-O-hydroxyl → HSL-C-carbonyl |
| **2** | nucleophile angle | Ser1β-O-hydroxyl > HSL-C-carbonyl > HSL-O |
| **3** | oxyanion stabilization 1 (Ala/Val) | Ala69β/Val70β-N-backbone → HSL-O-carbonyl |
| **4** | oxyanion stabilization 2 (Asn) | Asn241β/Asn269β-N$^{\delta}$ → HSL-O-carbonyl |
| **5** | additional stabilization (Gln23β/His23β) | Gln23β/His23β-O-backbone → HSL-H-amide |
| **6** | Ser1β hydroxyl stabilization | Ser1β-H-hydroxyl → Ser1β-N-amine |
| **7** | Ser1β amine stabilization 1 (Asn) | Ser1β-H1-amine → Asn241β/269β-N$^{\delta}$ |
| **8** | Ser1β amine stabilization 2 (Gln/His) | Ser1β-H2-amine → Gln23β-O$^{\epsilon}$/His23β-N$^{\delta}$ |
| **9** | oxyanion 1 to oxyanion 2 (Ala/Val to Asn) | Ala69β/Val70β-N-backbone → Asn241β/Asn269-N$^{\delta}$ |
| **10** | oxyanion 1 to nucleophile (Ala/Val to Ser) | Ala69β/Val70β-N-backbone → Ser1β-O-hydroxyl |
| **11** | oxyanion 2 to nucleophile (Asn to Ser) | Asn241β/Asn269β-N$^{\delta}$ → Ser1β-O-hydroxyl |
| **12** | Arg to HSL lactone oxygen | Arg263β/Arg297β-N-guanidyl → HSL-O-lactone |
| **13** | Ser1β amine to HSL leaving nitrogen | Ser1β-H-amine → HSL-N-amide |

**Molecular dynamics of protein-ligand complexes.** Water molecules around 2 Å to the protein were extracted for the frames of free enzyme MD simulations corresponding to selected ones from the docking study and placed back in the structure of the protein-ligand complex. Water molecules in too close contact with protein or ligand were removed to avoid potential steric clashes at the onset of simulations. Systems were prepared using an approach analogous to that applied in the free enzyme simulations with the tleap and Parmed modules of AmberTools18.[39] Minimization and equilibration were performed using the protocols applied in the free enzyme simulations with additional restraints on the ligand molecule. In the first minimization step, 500 kcal $mol^{-1}$ $Å^{-2}$ harmonic restraint was applied to ligand heavy atoms, followed by the following four minimization steps with decreasing restraints on amide bond atoms of the ligand equal 500, 125, 25 and 0.001 kcal $mol^{-1}$ $Å^{-2}$, respectively. In the first two stages of equilibration, the ligand heavy atoms were restrained using 5 kcal $mol^{-1}$ $Å^{-2}$, followed by the 25 kcal $mol^{-1}$ $Å^{-2}$ harmonic restraints on the distances as presented in **Table S2** (*"restraints for complexes MDs"*) covering nucleophile attack distance, oxyanion hole stabilizing hydrogen bonds, and proper orientation of the lactone ring oxygen towards stabilizing arginine, known to be important in the coordination of ligand orientation.[13,20] In the last step, the distance restraints gradually decreased from 25 to 0 kcal $mol^{-1}$ $Å^{-2}$. Finally, unrestrained NPT production simulations of 50 ns were performed at 310 K using the Langevin thermostat and controlling pressure using MC barostat. Fifteen independent replicates (each including the minimization and equilibration steps as well as the production run) were performed for all ligand-protein simulations to effectively profile stability of reactive-like complexes, for which even ensembles of markedly shorter simulations were shown as sufficient.[40,41] The stability of the production runs was evaluated in terms of the RMSD of the protein backbone heavy atoms, as in the free enzyme simulations (**Figure S4-S5** and **S21**), using the Cpptraj module of AmberTools18.

**Complexes binding free energy and ligand stabilization estimation.** The MMPBSA.py module of the AmberTools18 package was used to estimate the binding free energy of the complexes using the molecular mechanics/generalized Born surface area (MM/GBSA).[42] Complex, receptor and ligand topologies were generated using ante-MMPBSA.py script of the AmberTools18. Representative snapshots corresponding to the ligand adequately stabilized in the active site were extracted using mechanism-based criteria analogous to the previous one, used for docking poses selection, including oxyanion hole stabilization and nucleophile distance (**Table S2**

"*selection of complexes for MDs*"). Preselected frames were merged into a single ensemble for each complex separately and calculations were performed with Generalized Born implicit solvent model 8 with recommended mbondi3 set of atomic radii[43] at salt concentration of 0.1 M. A per-residue binding free energy decomposition was generated and the results were filtered to extract residues with the greatest contributions, i.e. those with contributions below -0.5 kcal/mol (for favorable contributions) and above 0.5 kcal/mol (for unfavorable contributions).

**Quantum Mechanics/Molecular Mechanics MD simulations.** QM/MM MD simulations were performed using the sander module of the Amber18 package. The initial frame for each protein-ligand complex was extracted from the standard MD simulations by searching for reactive-like configurations satisfying the criteria listed in **Table S2,** ("*selection of representative configurations for sMDs*") composed of (i) close nucleophile attack distance, oxyanion hole stabilization, additional stabilization of HSL by Gln23β/His23β, catalytic serine hydroxyl hydrogen oriented towards its amine, stabilization of the catalytic serine amine group, and appropriate nucleophile attack angle. Only for the aPGA-C08-HSL complex, these criteria could not be fully applied to obtain the desired configuration (see **Table S20**). Systems were further equilibrated by performing a 10 ns NPT simulation with the same settings as for protein-ligand production runs with additional 25 kcal $mol^{-1}$ $Å^{-2}$ harmonic restraints on distances described above for the selection of reactive-like configuration to achieve consistent starting structures for the QM/MM MD simulations across all systems. Production simulations of 500 ns were then performed for each complex with distance restraints on selected key interactions (**Table S2** *"restraints for sMDs inputs generation*") including nucleophile distance, oxyanion hole stabilization, additional stabilization by Gln23β/His23β, and proper stabilization of the nucleophile serine amine group, collecting restart files every 1 ns to obtain an ensemble of 500 equivalent starting points per complex. Each starting conformation was then equilibrated in 25 ps QM/MM MD simulations while applying the distance restraints analogous to restarts generation specified in **Table S2** (*"restraints for 1$^{st}$ QM/MM equilibration"*) using 1 fs time-step. The QM region consisted of HSL substrate, and residues: Ser1β, Ala69β/Val70β, Asn241β/Asn269β, Gln23β/His23β, and Arg263β/Arg297β and all necessary neighboring atoms from Thr68β/Thr69β, Pro22β, and Phe24β necessary to keep the junctions on non-polar carbon-carbon bonds as presented on **Figure S2**. The interface between QM and MM region was capped using eight linking hydrogen atoms. The QM/MM interface was treated with an electronic embedding scheme and

SHAKE algorithm inactivated. The QM region was described using the PM6-D semi-empirical method,[44–46] while the rest of the system was treated at the MM level using the ff14SB force field.

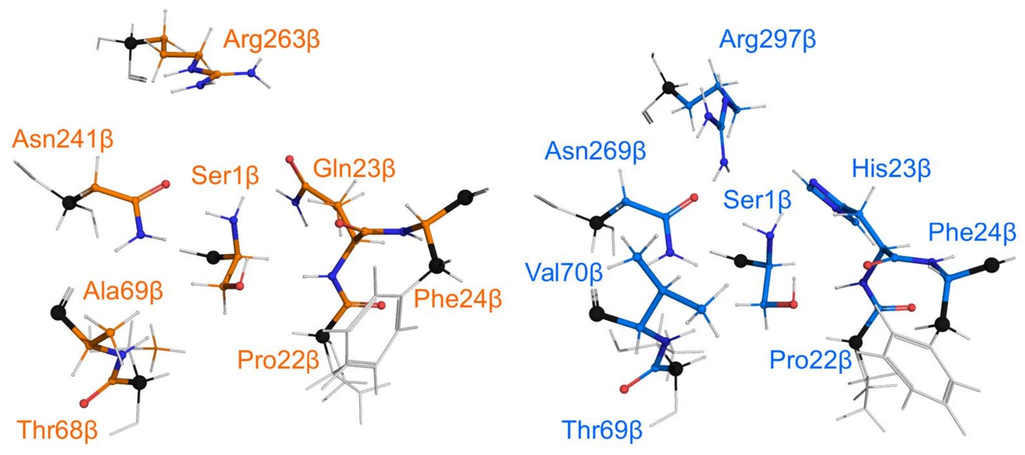

**Figure S2.** Definition of regions treated with QM in ecPGA (left, orange) and paPvdQ (right, blue) enzymes. The junctions between MM and QM regions are shown as black spheres and constitute the position of eight hydrogen linking atoms.

After initial QM/MM MD equilibration, distance restraints were turned off and steered QM/MM MD simulations of the acylation process were performed. The process was divided into two steps: formation of the tetrahedral intermediate (TI) and collapse of TI to form the acyl-enzyme (AE). In the first step, the reaction coordinate (RC) was represented as a linear combination of distances (LCOD) involved in the proton transfer. The linear combination was d1 – d2, where d1 is the distance from the serine amine nitrogen to the proton of the hydroxyl group and d2 is the distance between the proton and the oxygen of the hydroxyl group as shown in **Figure 2A**. This RC was steered from 1.1 to -1.1 Å with a harmonic restraint of 1000 kcal $mol^{-1}$ Å$^{-2}$ to satisfy the stiff-spring approximation necessary for the subsequent potential of mean force (PMF) profile calculation.[47] Steered MD simulation runs of the first stage were simulated for 35 ps resulting in approximately 0.06 Å/ps steering speed. After completion, each system was checked in terms of correctness of TI formed by the following criteria: (i) no instabilities occurred during simulation that finished successfully, (ii) covalent bond between serine hydroxyl oxygen and substrate carbonyl carbon was formed and no other changes were observed in bonding among heavy atoms, (iii) serine hydroxyl hydrogen was transferred to the amine group of the same catalytic serine amine nitrogen and no other proton jumps were observed. Correctly formed TIs were then equilibrated on the same level of theory as before, with a distance restraints on the newly formed covalent bond (1.5 Å) and the distance from the serine amine hydrogen to the substrate's amide nitrogen (2.0 Å), as shown in **Table S2** (*"restraints for 2$^{nd}$ QM/MM equilibration"*) using a force constant of 25 kcal mol$^{-1}$ Å$^{-2}$. This was followed by the validation to ensure that the simulations were stable, the newly created C-O bond in the previous reaction step was maintained, and transferred proton remained on the serine amine group and no other changes were observed.

Successfully equilibrated TIs were used as starting points for the second stage of the acylation reaction. Here, the RC was represented as a LCOD of three distances relevant to the formation of the AE, as shown in **Figure 2B:** d3 – d4 + d5, where d3 is the distance from the carbonyl carbon of the HSL to its amide N, d4 is the distance from the amide N to the closest proton of the serine amino group, and d5 is the distance from the nitrogen of the serine amino group to the proton being transferred. This RC was steered from 0.6 to 4.9 Å with a harmonic restraint of 1000 kcal mol$^{-1}$ Å$^{-2}$. The second stage of the acylation was simulated for 70 ps resulting in approximately 0.06 Å/ps steering speed similar to the previous stage. Simulations were validated in terms of correctness of AE formed according to the following criteria: (i) no instabilities occurred during simulation that finished successfully, (ii) hydrogen was transferred from the serine amine group to the substrate leaving nitrogen and no additional hydrogen jumps were observed, (iii) substrate amide bond was cleaved without any other changes in bonding among heavy atoms. Only steered QM/MM MD simulations verified for both the first and second stages were further analyzed (**Table S11 and S21**).

The results of these verified simulations were used as inputs to generate the PMF profiles using Jarzynski's equality.[48] Errors in the PMF profiles were estimated using a block-averaging scheme dividing each dataset corresponding to a particular complex into blocks containing approximately 20 repetitions (to preserve equal or near-equal block size).

During the development of steered QM/MM MD simulation protocol described above, the following parameters were optimized to balance the accuracy and the calculation throughput: the minimal QM region size, usage of the smallest steering force constant fulfilling the stiff-spring approximation,[47,48] and the maximal steering speed (**Table S3**).

**Table S3.** Testing of the sMD protocol sensitivity for various steering speeds.

| stage | length [ps] | number of repetitions | TS1/TS2 [kcal/mol] | TI/AE [kcal/mol] |
|---|---|---|---|---|
| ecPGA-C08-HSL | | | | |
| TI | 35 | 53 | 9.5 ± 0.8 | -11.7 ± 1.9 |
| | 70 | 39 | 9.2 ± 0.8 | -8.3 ± 1.0 |
| AE | 70 | 53 | 6.9 ± 0.9 | -1.5 ± 2.1 |
| | 140 | 39 | 7.3 ± 0.1 | -0.2 ± 1.2 |
| paPvdQ-C08-HSL | | | | |
| TI | 35 | 72 | 7.3 ± 0.7 | -16.8 ± 2.0 |
| | 70 | 72 | 7.4 ± 0.7 | -14.9 ± 1.6 |
| AE | 70 | 72 | 9.5 ± 1.1 | 2.8 ± 1.9 |
| | 140 | 72 | 8.8 ± 1.1 | 1.6 ± 1.5 |

**Table S4.** Retention times of HSLs obtained from HPLC measurement.

| Substrate | Retention time [min] |
|---|---|
| C06-HSL | 15.8 |
| C08-HSL | 28.4 |

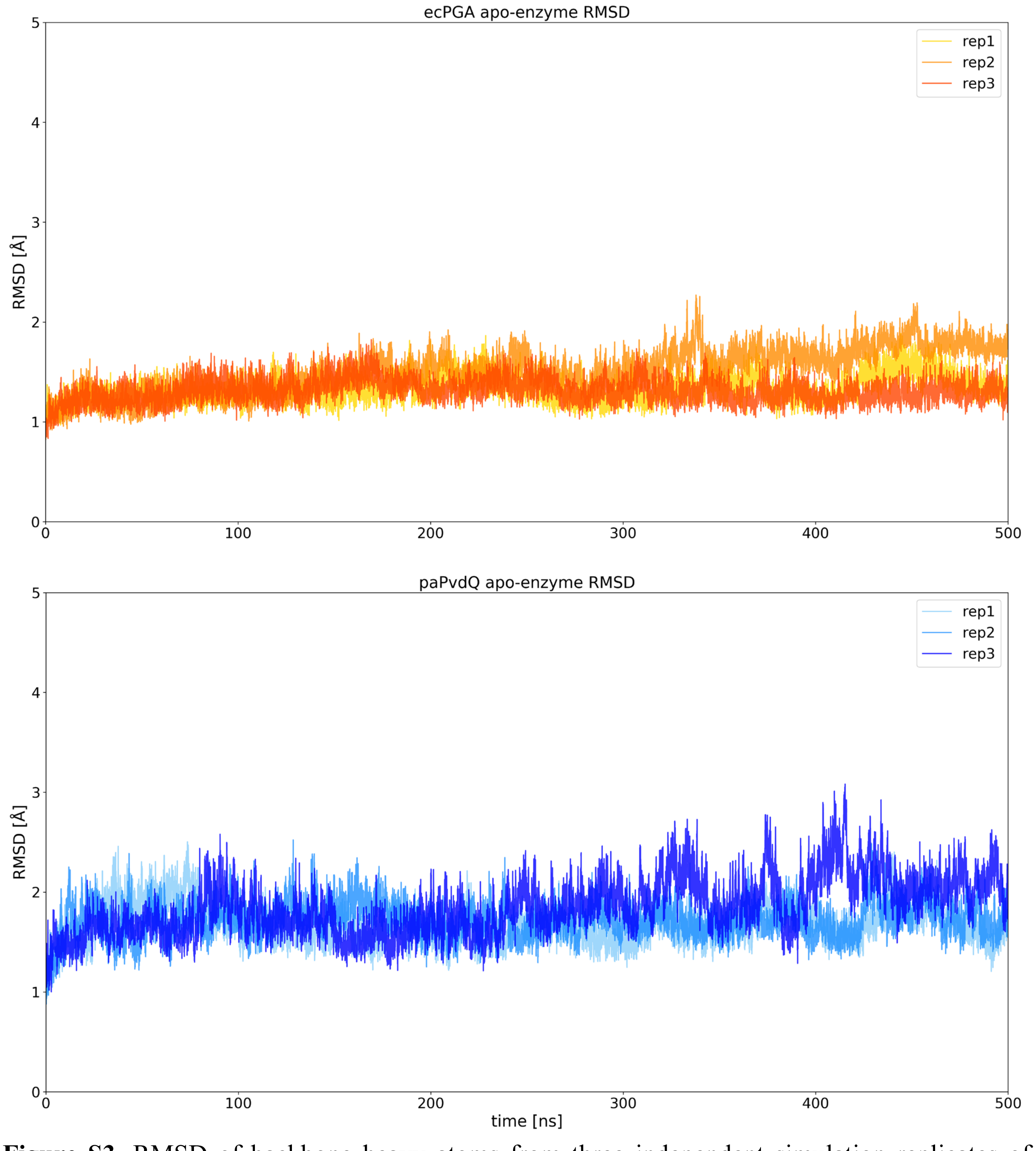

**Figure S3.** RMSD of backbone heavy atoms from three independent simulation replicates of ecPGA (top) and paPvdQ (bottom) in the absence of substrate.

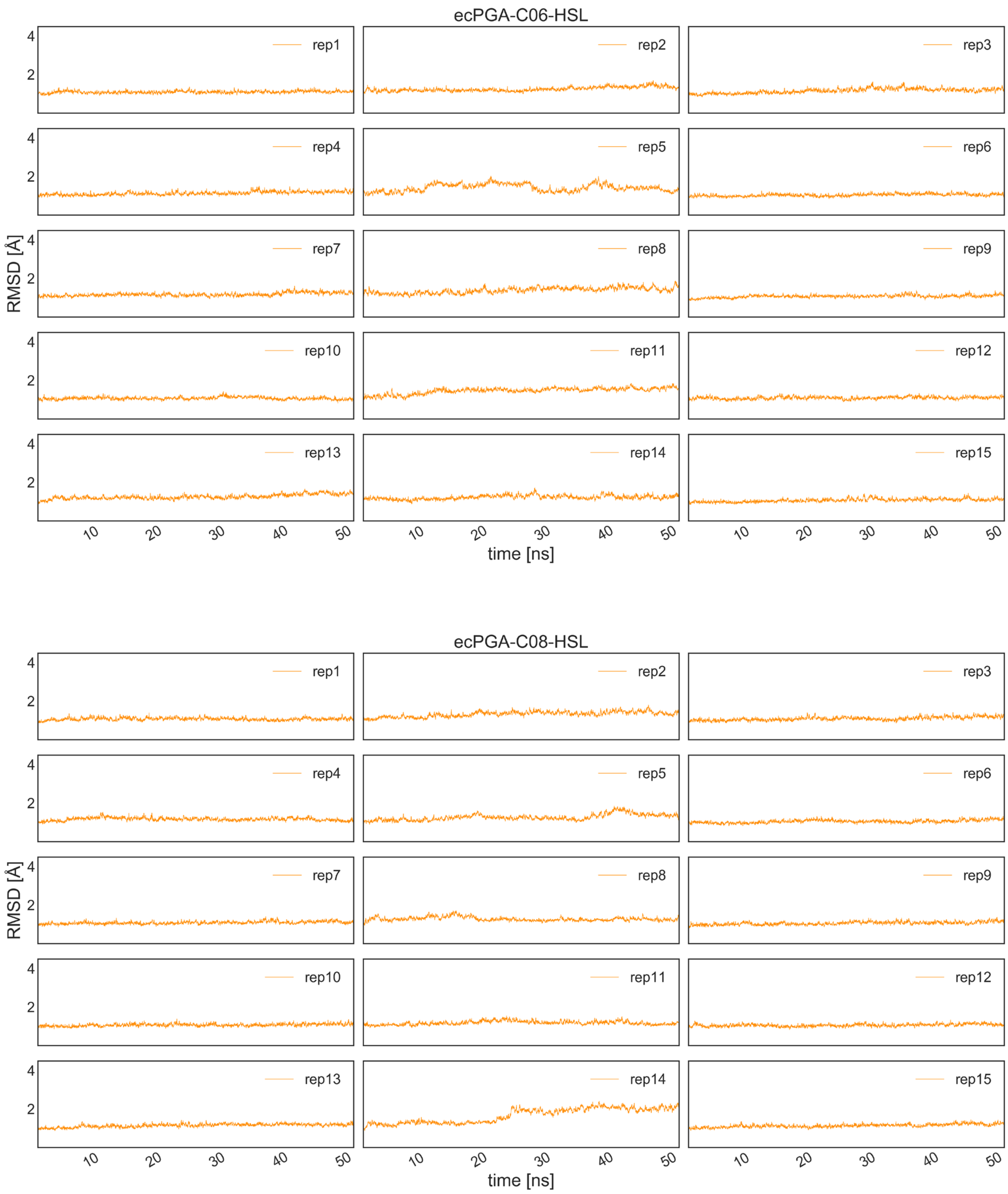

**Figure S4.** RMSD of backbone heavy atoms from 15 independent simulation replicates of ecPGA-C06-HSL (top) and ecPGA-C08-HSL (bottom) complexes.

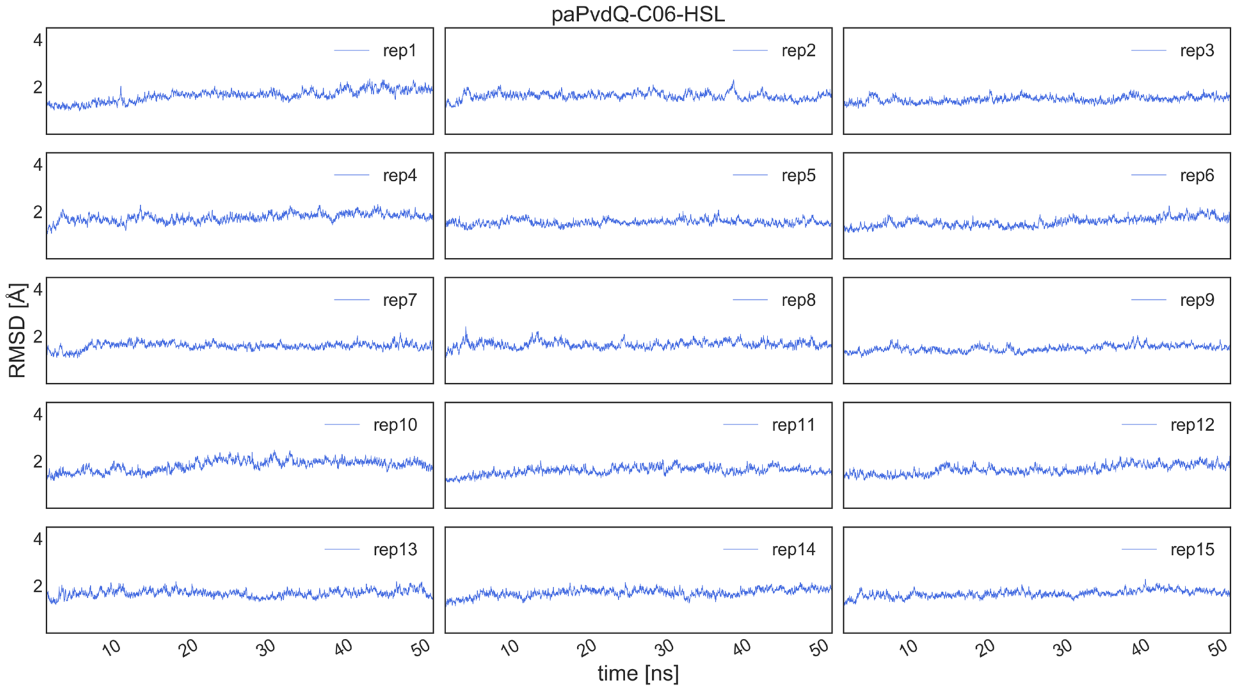

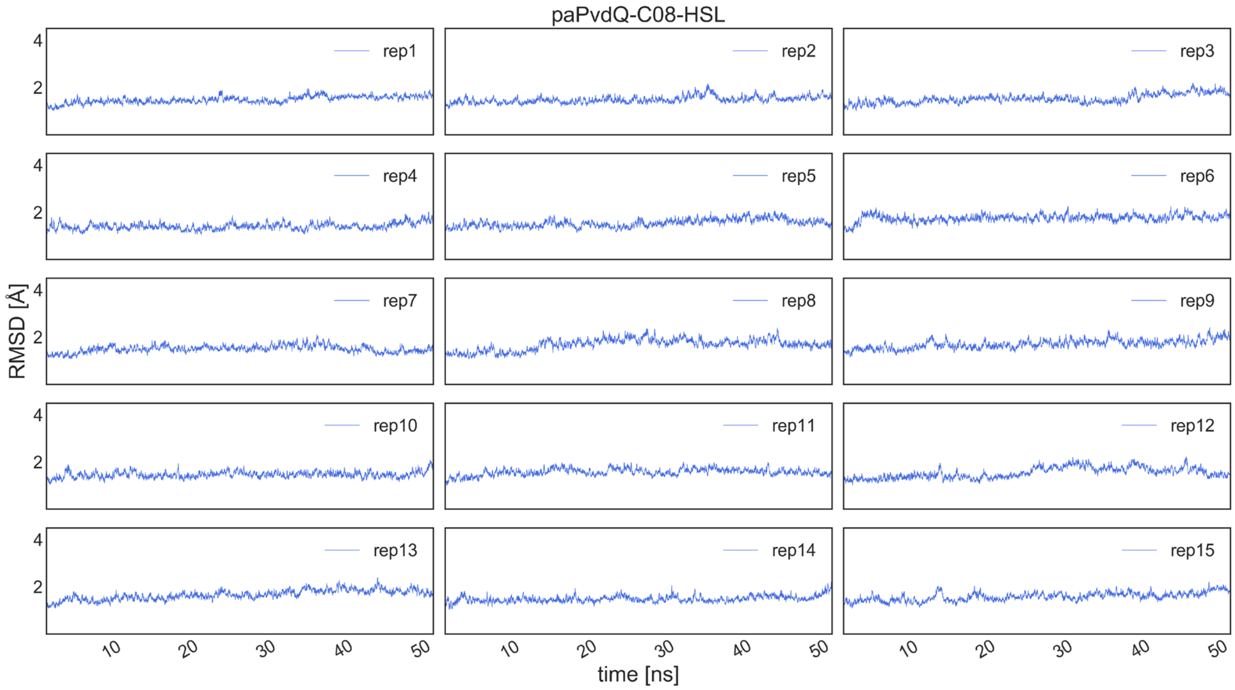

**Figure S5.** RMSD of backbone heavy atoms from 15 independent simulation replicates of paPvdQ-C06-HSL (top) and paPvdQ-C08-HSL (bottom) complexes.

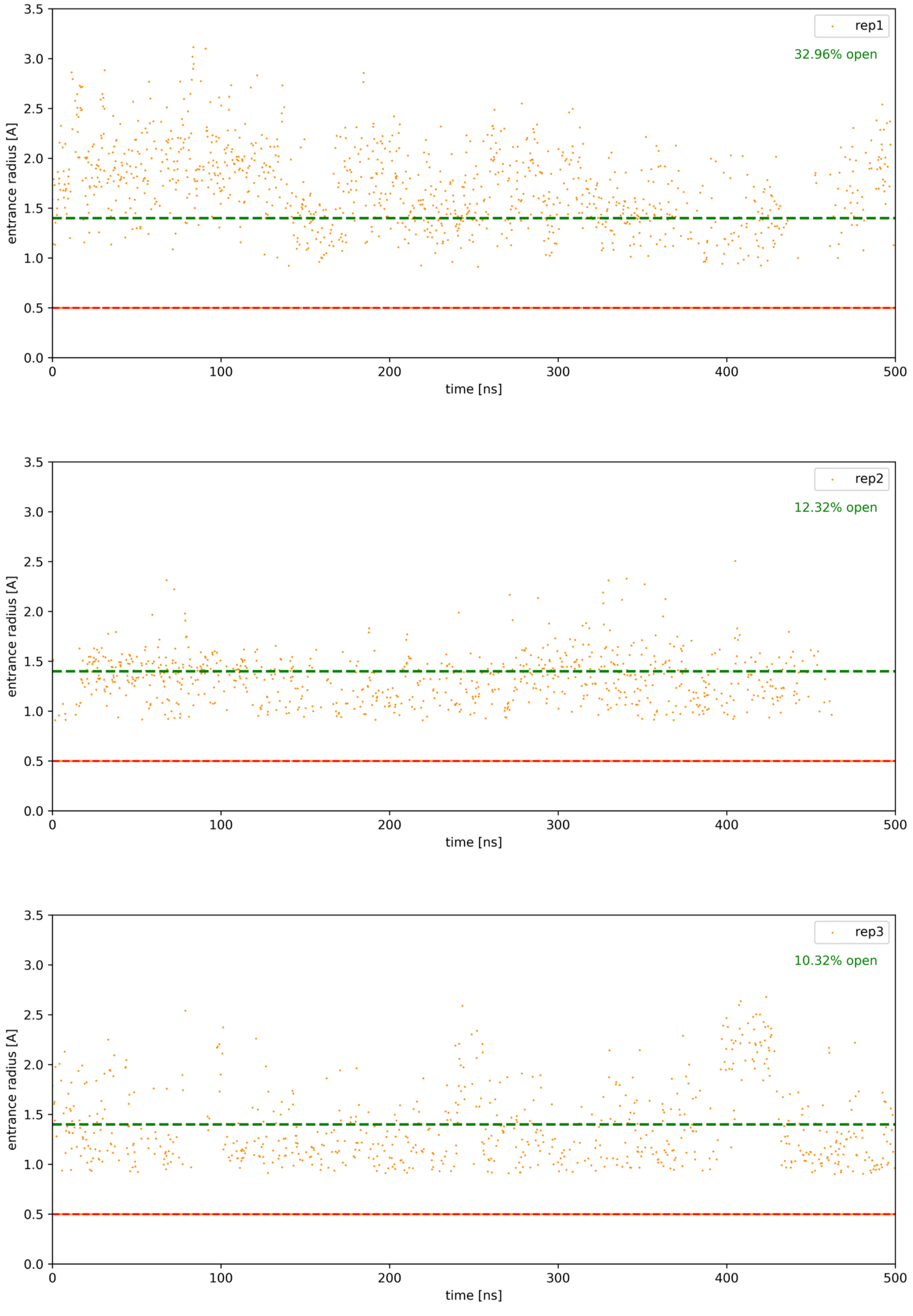

**Figure S6.** Time evolution of the entrance bottleneck to the acyl-binding cavity in ecPGA. The open entrance is defined as 1.4 Å (green dashed line). For the frames where CAVER did not detect the pocket, the entrance bottleneck was defined as the searching probe of 0.5 Å (red dashed line).

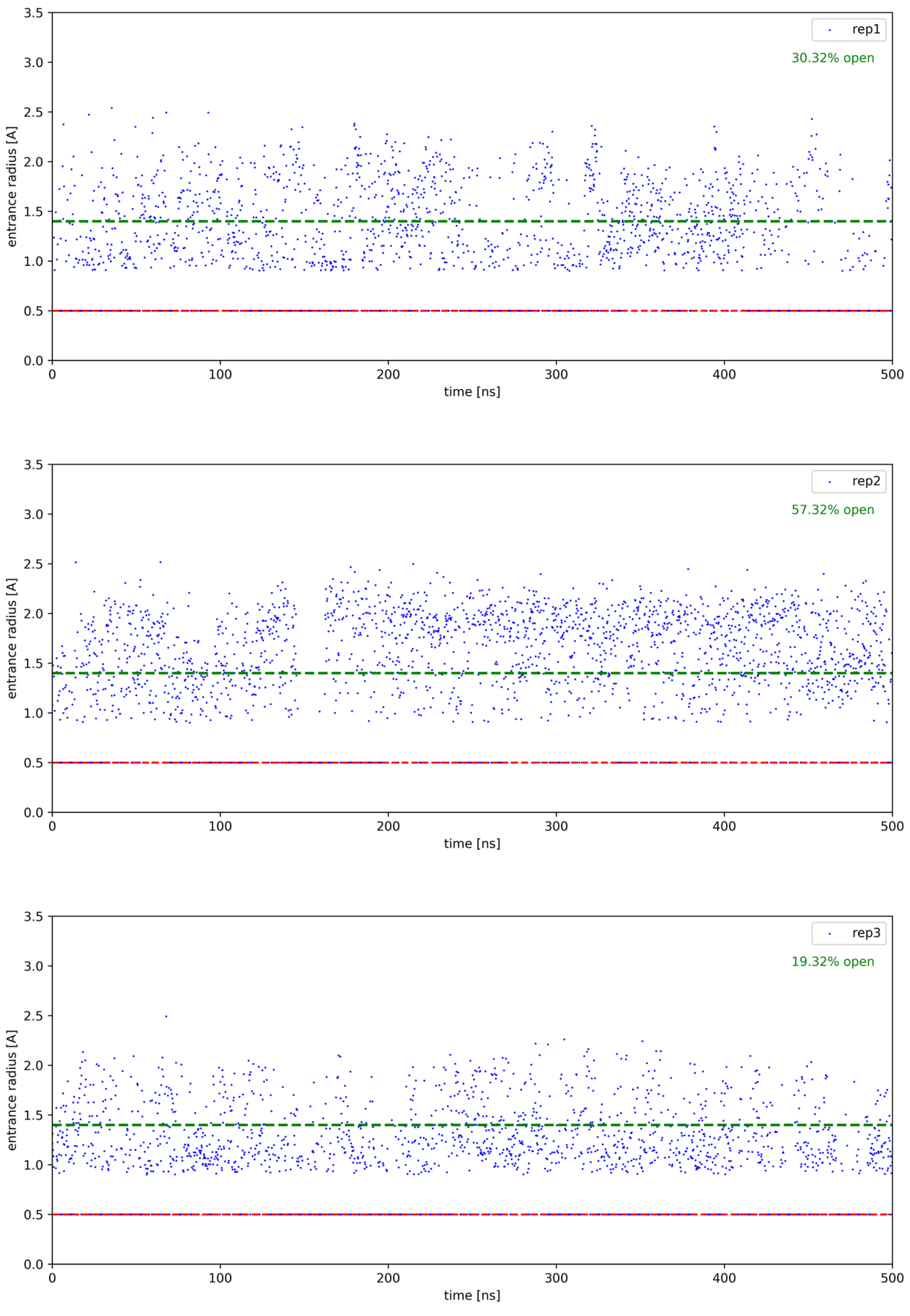

**Figure S7.** Time evolution of the entrance bottleneck to the acyl-binding cavity in paPvdQ. The open entrance is defined as 1.4 Å (green dashed line). For the frames where CAVER did not detect the pocket, the entrance bottleneck was defined as the searching probe of 0.5 Å (red dashed line).

**Table S5.** Opening of the acyl-binding cavity during free molecular dynamics simulation sufficient for HSLs binding for ecPGA and paPvdQ enzymes.

| protein | replica | cluster size | analyzed frames | well-shaped* | for docking** |
|---|---|---|---|---|---|
| ecPGA | 1 | 1080 | 2500 | 102 | 27 |
| | 2 | 782 | 2500 | 143 | 130 |
| | 3 | 716 | 2500 | 29 | 13 |
| paPvdQ | 1 | 1472 | 2500 | 106 | 84 |
| | 2 | 1944 | 2500 | 244 | 213 |
| | 3 | 1466 | 2500 | 87 | 0 |

*Well-shaped binding sites for HSLs defined based on the radius of the entrance bottleneck (narrowest point) > 1.4 Å, minimum depth of the cavity (corresponding to the acyl-chain length) > 5.0 Å, and the real starting point for the calculation is within 1Å to the desired location.

**Filtering for docking based on the favorable arrangement of catalytic residues promoting productive binding of HSLs defined as: Ser1β-H-hydroxyl → Ser1β-N-amine < 3.0 Å, Asn241β/Asn269β-N$^{\delta}$ → Ala69β/Val70β-N-backbone < 5.0 Å, Asn241β/Asn269β-N$^{\delta}$ → Ser1β-O-hydroxyl < 5.0 Å, Ala69β/Val70β-N-backbone → Ser1β-O-hydroxyl < 5.0 Å from PCA analysis of representative conformational states.

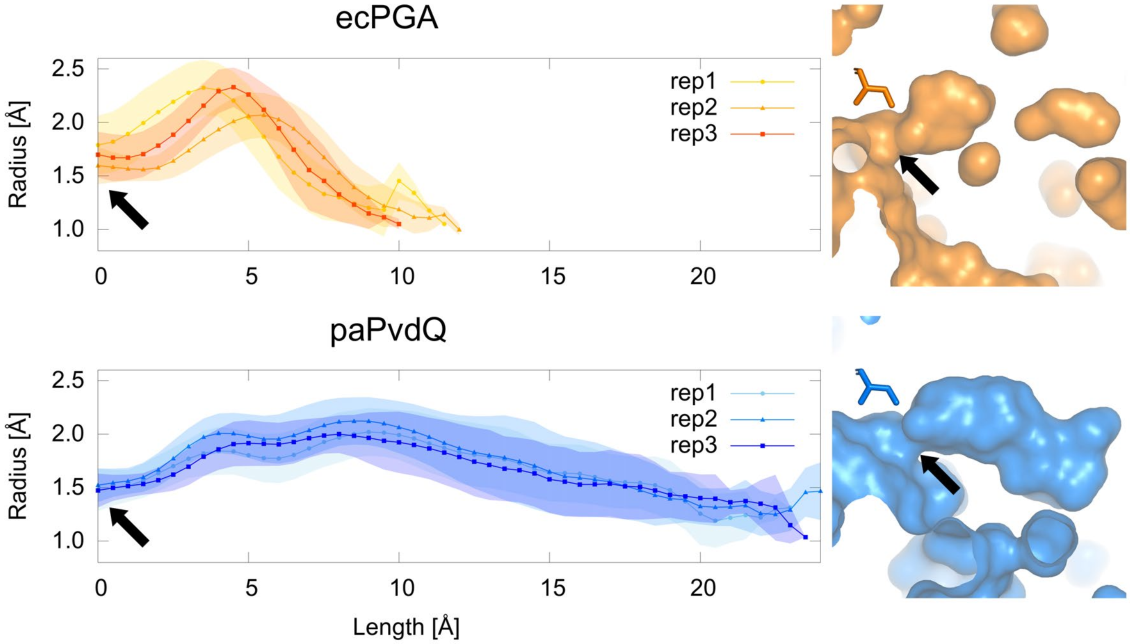

**Figure S8.** Well-shaped binding cavities profiles of ecPGA and paPvdQ enzymes in terms of HSLs binding as defined in **Table S5** (left). The volumetric shape of the cavity is shown as surface representation based on the protein crystal structures (right) for ecPGA (orange) and paPvdQ (blue). The narrowest part of the entrance to the acyl-binding cavity is indicated by black arrows.

**Table S6.** The geometry of representative states of catalytic residues extracted from the principal component analysis of free-enzyme molecular dynamics simulations for ecPGA and paPvdQ.

| Protein | State | Probability density | distance [Å] | | | |
|---|---|---|---|---|---|---|
| | | | AsnN → Ala/ValN[a] | AsnN → SenO[b] | Ala/ValN → SenO[c] | Gln/HisO → SenO[d] |
| ecPGA | a | 0.57 | 3.5 | 2.9 | 3.2 | 5.3 |
| | b | 0.13 | 4.2 | 4.9 | 4.6 | 3.8 |
| | c | 0.02 | 6.1 | 4.3 | 3.3 | 6.9 |
| | d | 0.02 | 7.1 | 5.8 | 5.6 | 4.4 |
| paPvdQ | a | 0.28 | 3.5 | 3.0 | 2.9 | 5.1 |
| | b | 0.18 | 3.7 | 4.4 | 4.4 | 4.0 |
| | c | 0.05 | 6.6 | 5.6 | 3.0 | 5.5 |
| | d | 0.09 | 6.5 | 6.8 | 4.6 | 4.3 |

[a] Asn241β/Asn269β-N$^{\delta}$ → Ala69β/Val70β-N-backbone
[b] Asn241β/Asn269β-N$^{\delta}$ → Ser1β-O-hydroxyl
[c] Ala69β/Val70β-N-backbone → Ser1β-O-hydroxyl
[d] Gln23β/His23β-O-backbone → Ser1β-O-hydroxyl

**Table S7.** Geometries and binding energies of representative protein-ligand complexes obtained in molecular docking experiment with strict filtering based on reaction mechanism for ecPGA and paPvdQ enzymes.

| protein | ligand | energy [kcal/mol] | distance [Å] SerO → HSL-C[a] | Ala/ValNH → HSL-O[b] | $AsnNH_2$ → HSL-O[c] |
|---|---|---|---|---|---|
| ecPGA | C06-HSL | -4.3 | 2.7 | 2.0 | 3.0 |
| | | -3.3 | 3.2 | 2.9 | 3.0 |
| | | -3.6 | 2.7 | 1.8 | 2.5 |
| | C08-HSL | -4.9 | 2.8 | 2.0 | 2.8 |
| | | -3.9 | 3.2 | 2.9 | 2.8 |
| | | -4.1 | 2.8 | 1.7 | 2.7 |
| paPvdQ | C06-HSL | -3.9 | 2.8 | 2.0 | 2.8 |
| | | -4.1 | 2.7 | 1.9 | 2.3 |
| | C08-HSL | -4.8 | 2.7 | 2.0 | 3.0 |
| | | -5.0 | 2.7 | 2.1 | 3.0 |

[a] Ser1β-O-hydroxyl → HSL-C-carbonyl
[b] Ala69β/Val70β-H-backbone → HSL-O-carbonyl
[c] Asn241β/Asn269β-$N^{\delta}$-$H_x$ (closest) → HSL-O-carbonyl

**Table S8.** Absolute binding free energies from MMGB/SA calculations for protein-ligand complexes properly stabilized during molecular dynamics simulations of ecPGA and paPvdQ complexes.

| protein | ligand | binding energy [kcal/mol] |
|---|---|---|
| ecPGA | C06-HSL | -31 ± 3 |
| | C08-HSL | -33 ± 4 |
| paPvdQ | C06-HSL | -28 ± 4 |
| | C08-HSL | -33 ± 3 |

**Table S9.** PCA weights for ecPGA and paPvdQ enzymes' catalytic machinery dynamics in ligand free form.

| PC/distance | SerNH1 → SerO[a] | SerNH2 → SerO[b] | SerH → SerN[c] | AsnN → Ala/ValN[d] | AsnN → SerO[e] | Ala/ValN → SerO[f] | Gln/HisO → SerO[g] |
|---|---|---|---|---|---|---|---|
| | | | | **ecPGA** | | | |
| PC1 (0.58) | 0.08 | 0.03 | 0.04 | 0.50 | 0.60 | 0.46 | -0.41 |
| PC2 (0.20) | -0.04 | -0.02 | -0.01 | 0.61 | 0.21 | -0.30 | 0.70 |
| | | | | **paPvdQ** | | | |
| PC1 (0.66) | 0.05 | 0.06 | 0.05 | 0.68 | 0.70 | 0.10 | -0.16 |
| PC2 (0.18) | 0.08 | 0.03 | 0.24 | 0.32 | -0.09 | -0.37 | 0.83 |

[a] Ser1β-H1-amine → Ser1β-O-hydroxyl
[b] Ser1β-H2-amine → Ser1β-O-hydroxyl
[c] Ser1β-H-hydroxyl → Ser1β-N-amine
[d] Asn241β/Asn269β-N$^{\delta}$ → Ala69β/Val70β-N-backbone
[e] Asn241β/Asn269β-N$^{\delta}$ → Ser1β-O-hydroxyl
[f] Ala69β/Val70β-N-backbone → Ser1β-O-hydroxyl
[g] Gln23β/His23β-O-backbone → Ser1β-O-hydroxyl

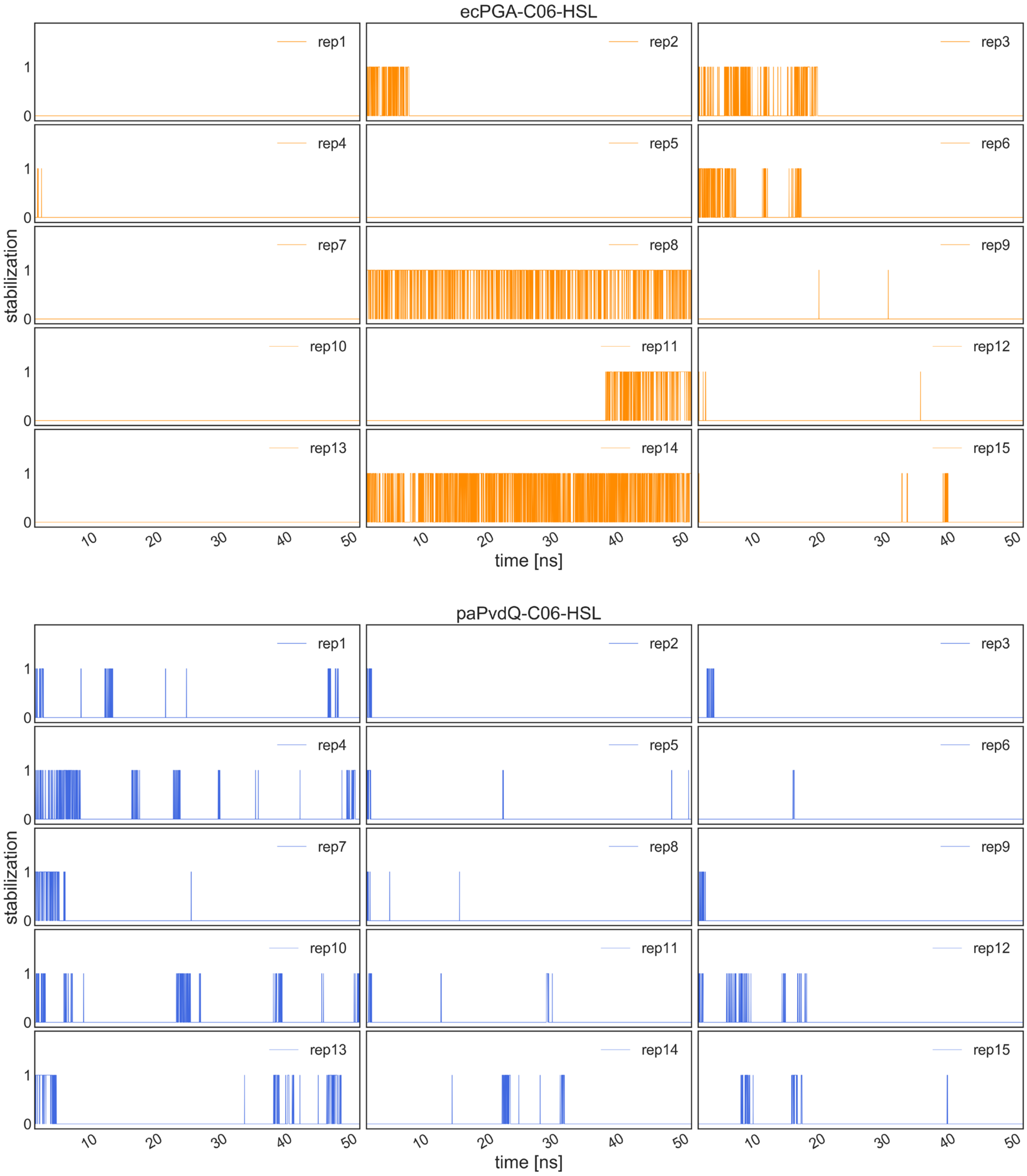

**Figure S9.** Simultaneous stabilization of ecPGA-C06-HSL (top, orange) and paPvdQ-C06-HSL bottom, blue) complexes in fully productive conformation.

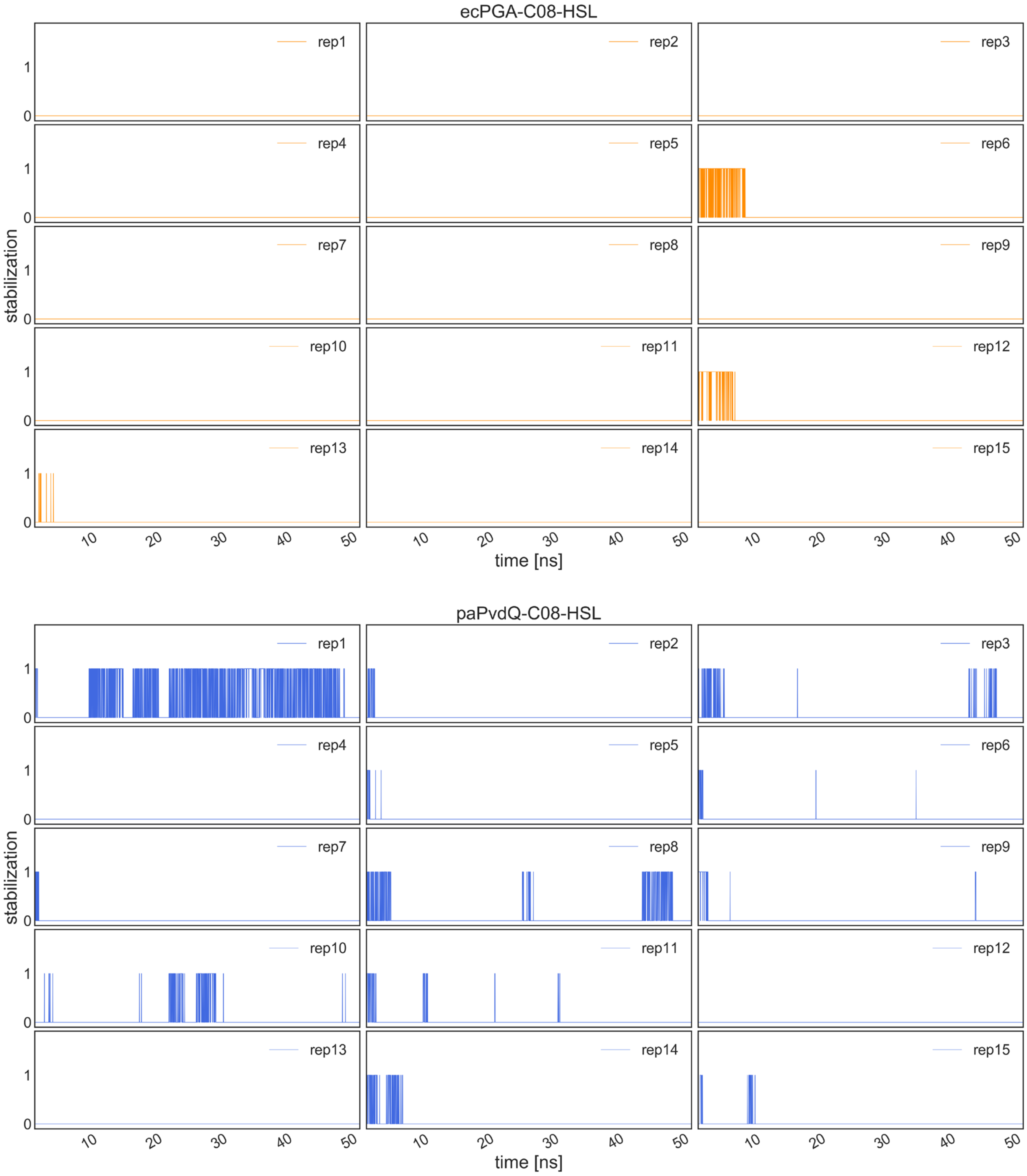

**Figure S10.** Simultaneous stabilization of ecPGA-C08-HSL (top, orange) and paPvdQ-C08-HSL (bottom, blue) complexes in fully productive conformation.

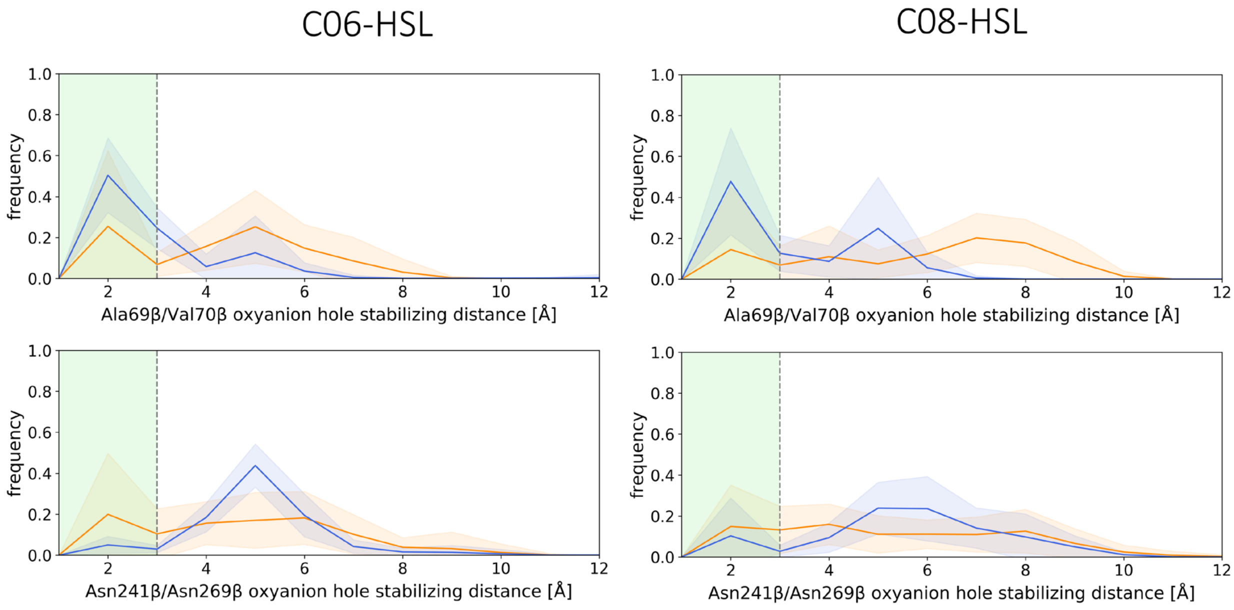

**Figure S11.** The distributions of oxyanion hole stabilizing distances for ecPGA and paPvdQ complexes. Ala/Val contribution (upper plots), Asn contribution (bottom plots) for C06-HSL (left column) and C08-HSL (right column) in complex with ecPGA (orange) and paPvdQ (blue). Green regions highlighted on the plots indicate the optimal range of the distance.

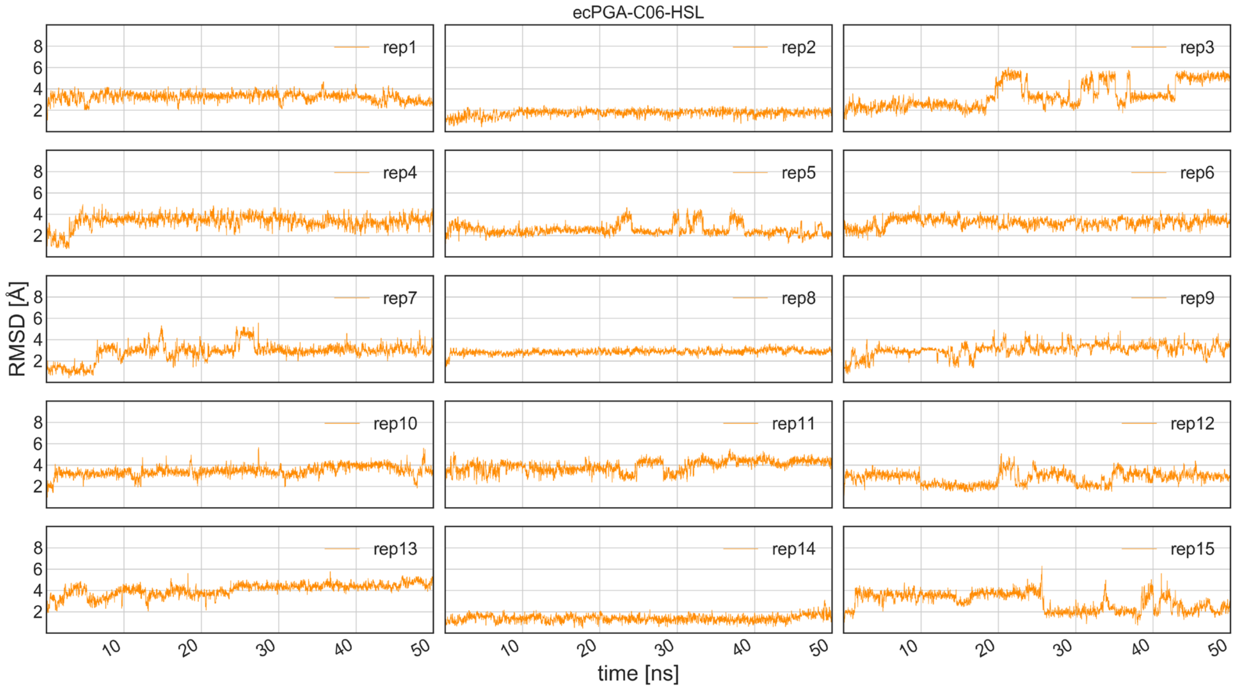

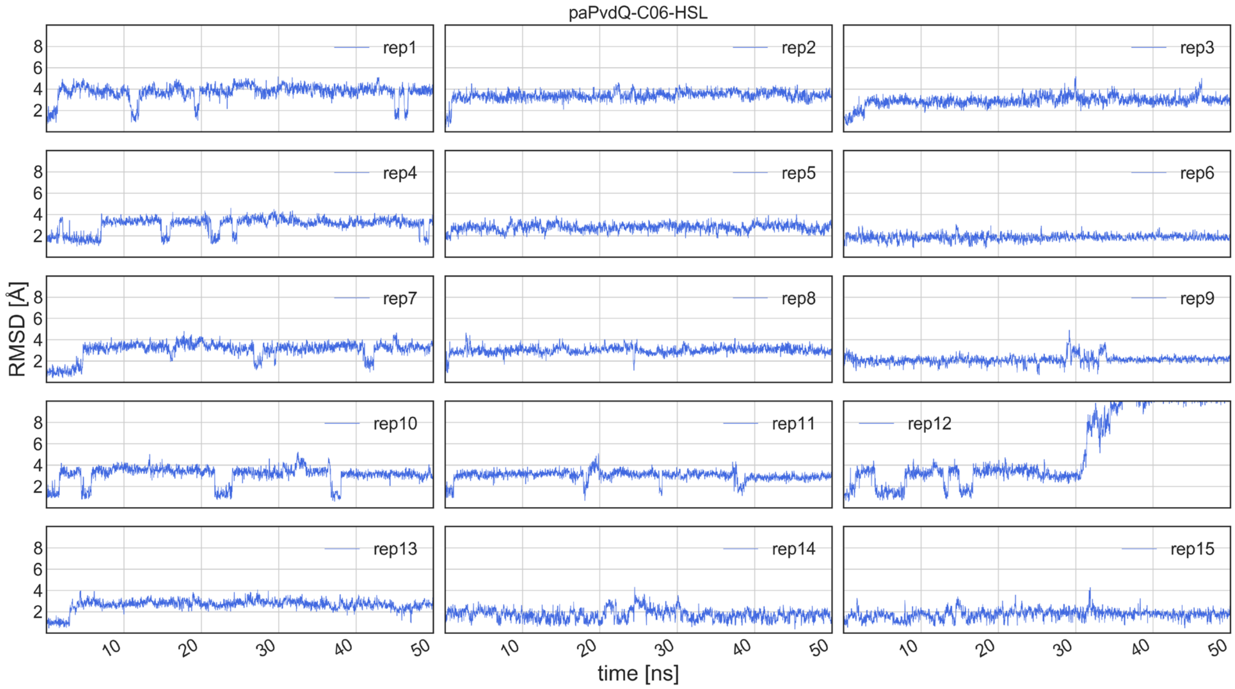

**Figure S12.** Ligand RMSD evolution for ecPGA-C06-HSL (top, orange) and paPvdQ-C06-HSL (bottom, blue) complexes across all protein-ligand complexes MD replicates.

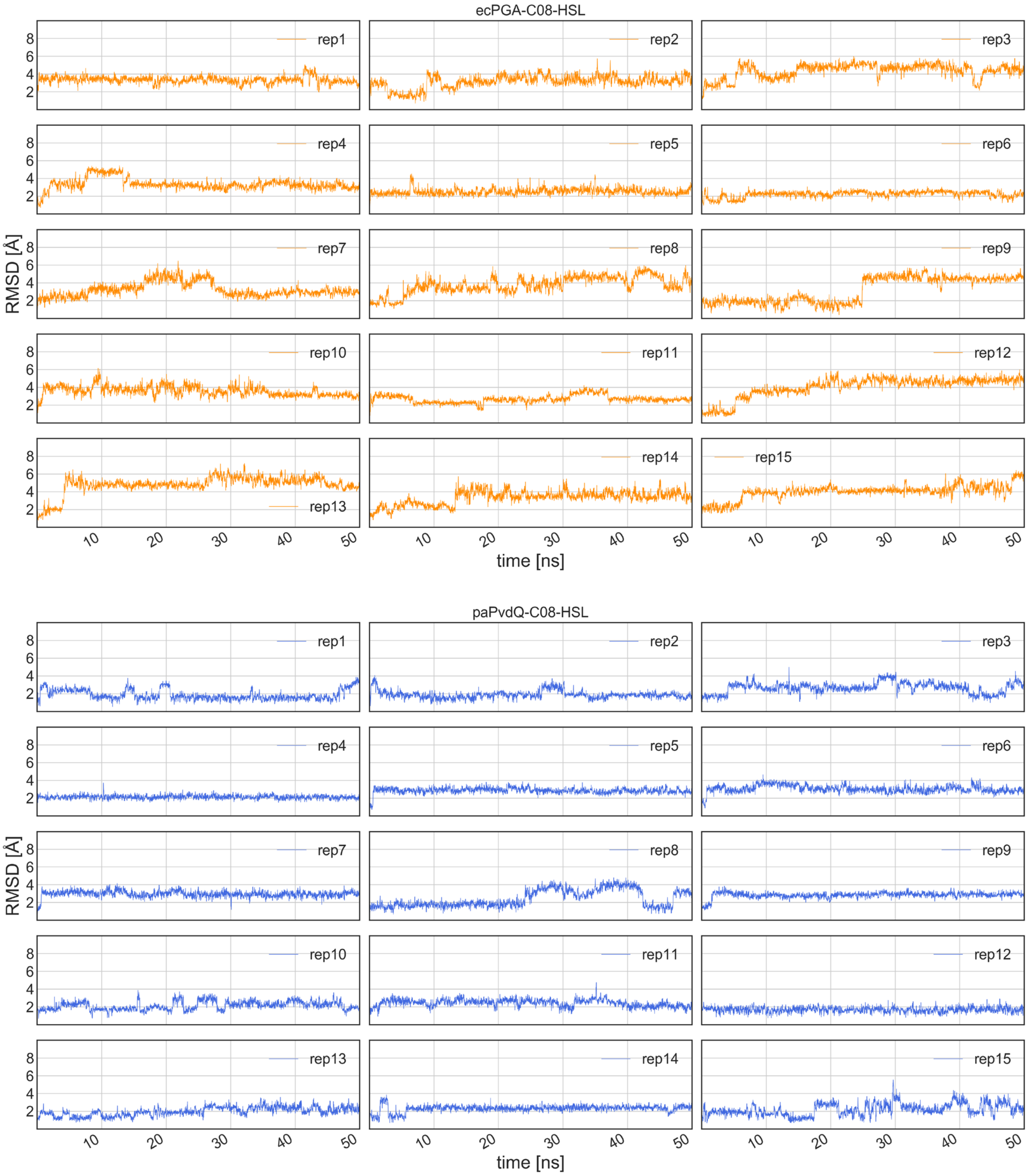

**Figure S13.** Ligand RMSD evolution for ecPGA-C08-HSL (top, orange) and paPvdQ-C08-HSL (bottom, blue) complexes across all protein-ligand complexes MD replicates.

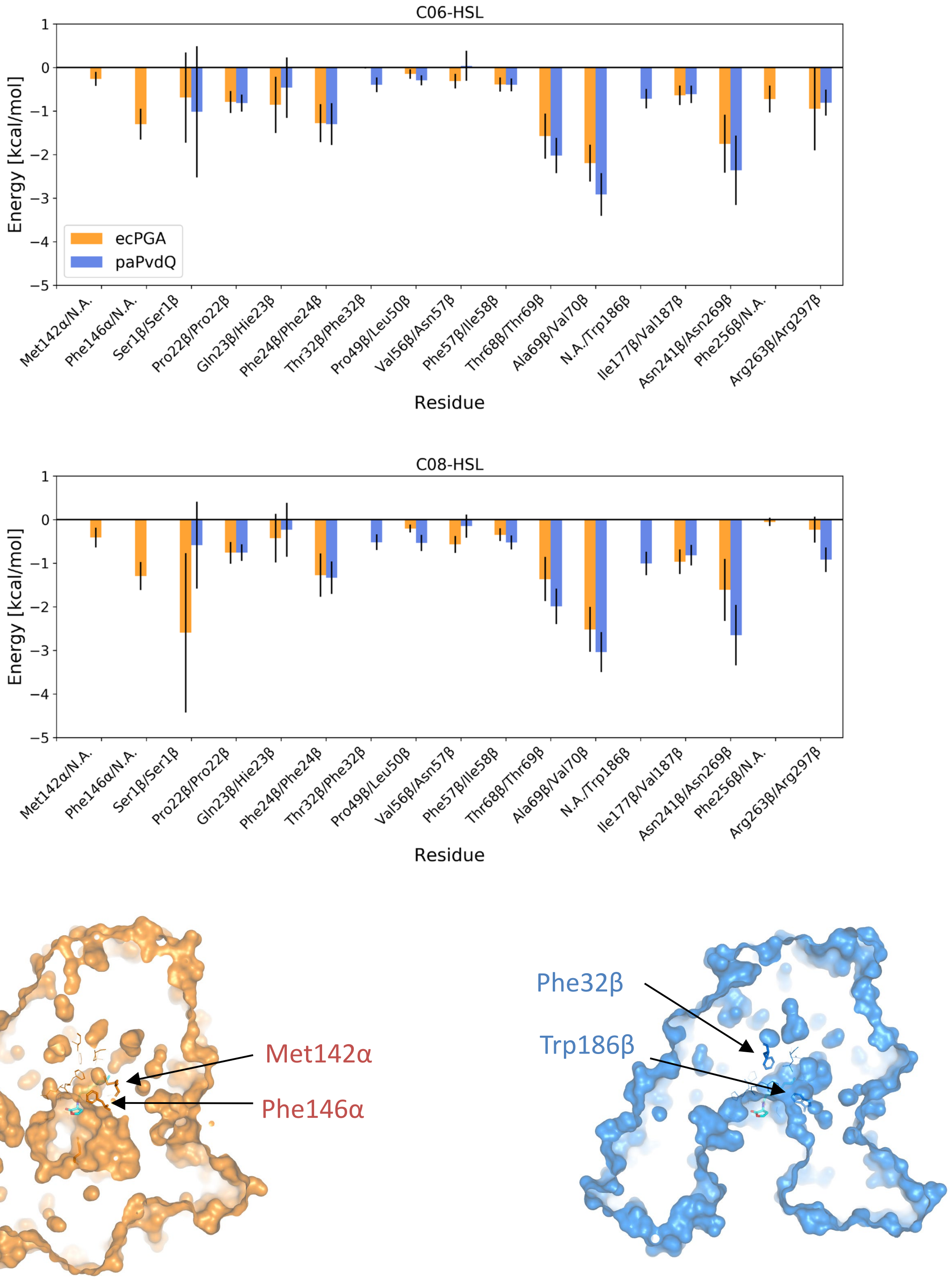

**Figure S14.** Per residue energy decomposition for enzyme-C06-HSL (top) and -C08-HSL (middle) complexes of ecPGA (orange) and paPvdQ (blue) from MMGB/SA binding energy calculations. In the bottom part of the panel, ecPGA (left, orange) and paPvdQ (right, blue) proteins are represented as the surface with all contributing residues shown as lines, while unique residues for given protein as sticks.

**Table S10.** Configurations of representative structures selected for generation of starting points for steered MD simulations of the acylation process for ecPGA and paPvdQ complexes.

| complex | distance [Å] | | | | | | | angle [°] |
|---|---|---|---|---|---|---|---|---|
| | SerO → HSL-C*[a] | $AsnNH_2$ → HSL-O**[b] | Ala/ValNH → HSL-O**[c] | Gln/HisO → HSL-H**[d] | SerH → SerN**[e] | $GlnO^{\varepsilon}/HisN^{\delta}$ → SerH**[f] | $AsnO^{\delta}$ → SerH**[g] | SerO > HSL-C > HSL-O***[h] |
| ecPGA-C06-HSL | 3.1 | 2.0 | 2.2 | 1.9 | 2.1 | 2.1 | 1.9 | 79 |
| ecPGA-C08-HSL | 2.9 | 2.0 | 1.9 | 2.6 | 2.1 | 2.0 | 1.9 | 103 |
| paPvdQ-C06-HSL | 2.8 | 2.0 | 1.9 | 2.2 | 2.0 | 2.2 | 1.9 | 83 |
| paPvdQ-C08-HSL | 3.0 | 2.0 | 2.1 | 2.1 | 2.0 | 2.5 | 1.9 | 88 |

* desired value below 3.3 Å, preferably close to 3.0 Å
** expected to be within hydrogen bond distance <3.0 Å
*** angle should be within range 75-105°, preferably close to 90°

[a] Ser1β-O-hydroxyl → HSL-C-carbonyl
[b] Asn241β/Asn269β-$N^{\delta}$-$H_x$ (closest) → HSL-O-carbonyl
[c] Ala69β/Val70β-H-backbone → HSL-O-carbonyl
[d] Gln23β/His23β-O-backbone → HSL-H-carbonyl
[e] Ser1β-H-hydroxyl → Ser1β-N-amine
[f] Gln23β-$O^{\varepsilon}$/His23β-$N^{\delta}$ → Ser1β-H-amine
[g] Asn241β/Asn269-$N^{\delta}$ → Ser1β-H-amine
[h] Ser1β-O-hydroxyl > HSL-C-carbonyl > HSL-O-carbonyl

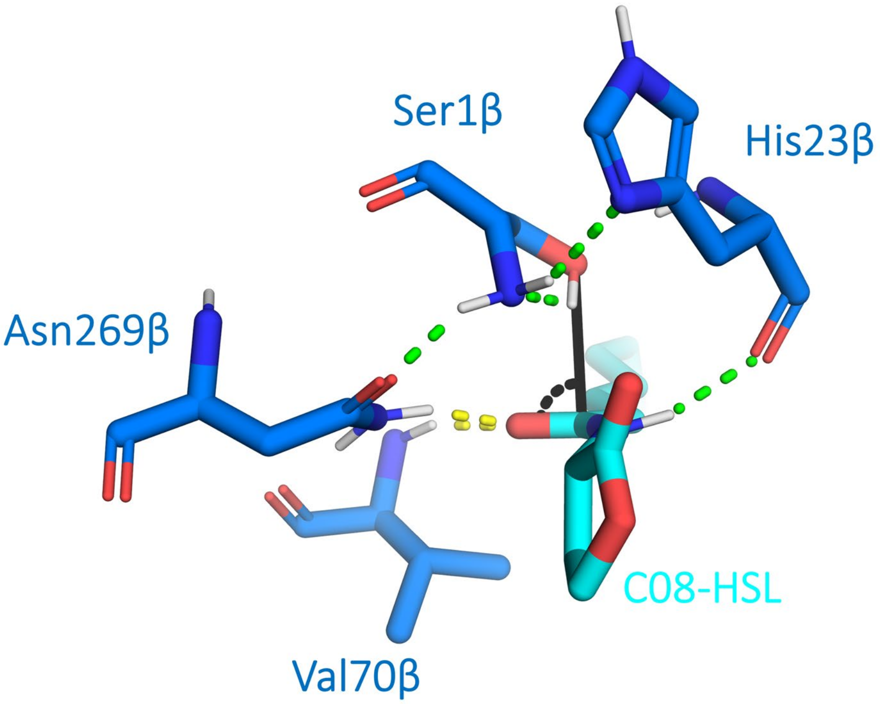

**Figure S15.** paPvdQ-C08-HSL complex showing the favorable organization of interactions network. Black and yellow lines correspond to criteria specified previously for the productive stabilization estimation, while green dashed lines are additional elements chosen to select a starting point for QM/MM steered MD simulations. Those include the stabilization by Gln23β/His23β residue, proper orientation of the hydroxyl hydrogen toward accepting amine group of the catalytic serine, and proper stabilization of this group by side chains of residues Asn241β/Asn269β and Gln23β/His23β.

**Table S11.** QM/MM MD sampling for ecPGA and paPvdQ complexes.

| complex | successful complete cycle-repetitions | TI-cumulative time [ns] | AE-cumulative time [ns] | total sMD time [ns] |
|---|---|---|---|---|
| ecPGA-C06-HSL | 224 | 7.8 | 15.7 | 23.5 |
| paPvdQ-C06-HSL | 350 | 12.3 | 24.5 | 36.8 |
| ecPGA-C08-HSL | 247 | 8.6 | 17.3 | 25.9 |
| paPvdQ-C08-HSL | 325 | 11.4 | 22.8 | 34.1 |

**Table S12.** Detailed average geometries of particular states ensembles during acylation reaction for ecPGA and paPvdQ complexes.

| complex | state | mean/ std | distance [Å] | | | | | | | angle [°] | |
|---|---|---|---|---|---|---|---|---|---|---|---|
| | | | SerO → HSL-C[a] | Ala/ValNH → HSL-O[b] | AsnNH2 → HSL-O[c] | Gln/HisO → HSL-H[d] | HSL-C → HSL-N[e] | ArgH → HSL-O[f] | ArgC →SerN[g] | ArgC > SerN > HSL-N[h] | HSL-N > SerN > ArgN1 > ArgN2[i] |
| ecPGA-C06-HSL | MC | mean | 2.8 | 2.3 | 2.3 | 2.8 | 1.4 | 4.0 | 5.2 | 109 | -74 |
| | | std | 0.3 | 0.4 | 0.5 | 0.8 | 0.1* | 1.3 | 0.5 | 9 | 17 |
| | TS1 | mean | 2.4 | 2.2 | 2.3 | 2.5 | 1.4 | 3.6 | 5.1 | 113 | -74 |
| | | std | 0.5 | 0.6 | 0.6 | 0.8 | 0.1* | 1.5 | 0.5 | 10 | 15 |
| | TI | mean | 1.6 | 1.9 | 1.9 | 2.3 | 1.5 | 3.3 | 5.0 | 119 | -78 |
| | | std | 0.1 | 0.2 | 0.2 | 0.4 | 0.1* | 1.4 | 0.3 | 8 | 12 |
| | TS2a | mean | 1.5 | 1.9 | 1.9 | 2.0 | 1.6 | 2.7 | 5.0 | 115 | -81 |
| | | std | 0.1* | 0.2 | 0.2 | 0.2 | 0.2 | 1.2 | 0.3 | 6 | 11 |
| | TS2b | mean | 1.4 | 2.0 | 2.0 | 2.0 | 2.0 | 3.1 | 4.8 | 116 | -84 |
| | | std | 0.1* | 0.2 | 0.2 | 0.2 | 0.4 | 1.3 | 0.4 | 8 | 13 |
| | AE | mean | 1.4 | 2.1 | 2.2 | 2.4 | 3.3 | 3.1 | 5.3 | 96 | -91 |
| | | std | 0.1* | 0.3 | 0.3 | 0.6 | 0.4 | 1.4 | 0.4 | 11 | 16 |
| paPvdQ-C06-HSL | MC | mean | 2.7 | 2.1 | 2.5 | 2.3 | 1.4 | 5.6 | 4.4 | 139 | -15 |
| | | std | 0.3 | 0.3 | 0.5 | 0.5 | 0.1* | 1.2 | 0.4 | 12 | 47 |
| | TS1 | mean | 2.2 | 2.0 | 2.2 | 2.1 | 1.4 | 5.0 | 4.4 | 144 | -5 |
| | | std | 0.3 | 0.3 | 0.4 | 0.3 | 0.1* | 1.2 | 0.3 | 9 | 43 |
| | TI | mean | 1.5 | 1.9 | 1.9 | 2.1 | 1.5 | 4.7 | 4.5 | 149 | -4 |
| | | std | 0.1 | 0.2 | 0.2 | 0.3 | 0.1* | 1.2 | 0.3 | 6 | 46 |
| | TS2a | mean | 1.5 | 1.9 | 2.0 | 1.9 | 1.6 | 4.6 | 4.5 | 144 | -28 |
| | | std | 0.1* | 0.2 | 0.2 | 0.2 | 0.1* | 1.0 | 0.2 | 5 | 37 |
| | TS2b | mean | 1.4 | 2.1 | 2.2 | 2.0 | 1.9 | 5.1 | 4.3 | 146 | -21 |
| | | std | 0.1* | 0.2 | 0.3 | 0.2 | 0.3 | 1.0 | 0.3 | 7 | 44 |
| | AE | mean | 1.4 | 2.1 | 2.3 | 2.8 | 3.4 | 5.6 | 4.6 | 118 | -38 |
| | | std | 0.1* | 0.2 | 0.4 | 0.9 | 0.3 | 1.4 | 0.3 | 11 | 32 |

*std rounded up to meaningful magnitudes and maintain consistent formatting; **Note: Table continuation on the following page**

**Table S12 continued.** Detailed average geometries of particular states ensembles during acylation reaction for ecPGA and paPvdQ complexes.

| complex | state | mean/ std | distance [Å] | | | | | | | angle [°] | |
|---|---|---|---|---|---|---|---|---|---|---|---|
| | | | SerO → HSL-C[a] | Ala/ValNH → HSL-O[b] | AsnNH2 → HSL-O[c] | Gln/HisO → HSL-H[d] | HSL-C → HSL-N[e] | ArgH → HSL-O[f] | ArgC →SerN[g] | ArgC > SerN > HSL-N[h] | HSL-N > SerN > ArgN1 > ArgN2[i] |
| ecPGA-C08-HSL | MC | mean | 3.0 | 2.6 | 2.3 | 2.8 | 1.4 | 3.3 | 5.3 | 103 | -72 |
| | | std | 0.3 | 0.6 | 0.5 | 0.7 | 0.1* | 1.0 | 0.5 | 9 | 17 |
| | TS1 | mean | 2.7 | 2.5 | 2.1 | 2.4 | 1.4 | 3.1 | 5.3 | 103 | -71 |
| | | std | 0.4 | 0.6 | 0.4 | 0.6 | 0.1* | 1.1 | 0.5 | 10 | 15 |
| | TI | mean | 1.6 | 2.0 | 1.9 | 2.2 | 1.5 | 2.6 | 5.0 | 116 | -76 |
| | | std | 0.1 | 0.2 | 0.2 | 0.3 | 0.1* | 1.0 | 0.3 | 7 | 15 |
| | TS2a | mean | 1.5 | 2.1 | 1.9 | 1.9 | 1.6 | 2.3 | 4.9 | 116 | -81 |
| | | std | 0.1* | 0.2 | 0.2 | 0.1 | 0.1* | 1.0 | 0.4 | 5 | 10 |
| | TS2b | mean | 1.4 | 2.1 | 2.1 | 2.0 | 1.8 | 2.6 | 4.9 | 114 | -79 |
| | | std | 0.1* | 0.3 | 0.2 | 0.2 | 0.2 | 1.0 | 0.3 | 6 | 11 |
| | AE | mean | 1.4 | 2.2 | 2.2 | 2.6 | 3.3 | 2.5 | 5.3 | 91 | -81 |
| | | std | 0.1* | 0.3 | 0.3 | 0.6 | 0.3 | 1.1 | 0.4 | 10 | 15 |
| paPvdQ-C08-HSL | MC | mean | 2.7 | 2.2 | 2.4 | 2.8 | 1.4 | 5.6 | 4.2 | 140 | 17 |
| | | std | 0.3 | 0.4 | 0.5 | 0.7 | 0.1* | 1.2 | 0.3 | 12 | 38 |
| | TS1 | mean | 2.3 | 2.1 | 2.2 | 2.5 | 1.4 | 5.1 | 4.3 | 143 | 23 |
| | | std | 0.3 | 0.3 | 0.3 | 0.6 | 0.1* | 1.1 | 0.3 | 10 | 27 |
| | TI | mean | 1.6 | 1.9 | 1.9 | 2.2 | 1.5 | 4.7 | 4.4 | 149 | 2 |
| | | std | 0.1 | 0.3 | 0.2 | 0.2 | 0.1* | 1.3 | 0.2 | 8 | 44 |
| | TS2a | mean | 1.5 | 2.0 | 2.0 | 1.9 | 1.6 | 4.7 | 4.3 | 144 | -19 |
| | | std | 0.1* | 0.3 | 0.2 | 0.2 | 0.1* | 1.1 | 0.3 | 6 | 31 |
| | TS2b | mean | 1.4 | 2.0 | 2.2 | 1.9 | 1.8 | 5.1 | 4.2 | 147 | -9 |
| | | std | 0.1* | 0.2 | 0.3 | 0.2 | 0.2 | 1.1 | 0.2 | 6 | 36 |
| | AE | mean | 1.4 | 2.2 | 2.3 | 2.9 | 3.3 | 5.3 | 4.4 | 119 | -26 |
| | | std | 0.1* | 0.4 | 0.4 | 0.9 | 0.3 | 1.4 | 0.3 | 12 | 25 |

* std rounded up to meaningful magnitudes and maintain consistent formatting; [a] Ser1β-O-hydroxyl → HSL-C-carbonyl; [b] Asn241β/Asn269β-N$^{\delta}$-Hx (closest) → HSL-O-carbonyl; [c] Ala69β/Val70β-H-backbone → HSL-O-carbonyl; [d] Gln23β/His23β-O-backbone → HSL-H-carbonyl; [e] HSL-C-carbonyl → HSL-N; [f] Arg263β/Arg297β-$NH_x$ (guanidyl closest) → HSL-O-lactone; [g] Arg263β/Arg297β-C-guanidyl → Ser1β-N-amine; [h] Arg263β/Arg297β-C-guanidyl > Ser1β-N-amine > HSL-N; [i] Ser1β-N-amine > HSL-N > Arg263β/Arg297β-N1-guanidyl > Arg263β/Arg297β-N2-guanidyl

**Table S13.** Energetics of the first step of the acylation reaction for ecPGA and paPvdQ complexes.

| complex | energy [kcal/mol] | | |
|---|---|---|---|
| | MC | TS1 | TI |
| ecPGA-C06-HSL | 0 | 7.0 ± 1.1 | -18.7 ± 2.3 |
| ecPGA-C08-HSL | 0 | 9.5 ± 0.8 | -11.1 ± 1.8 |
| paPvdQ-C06-HSL | 0 | 7.3 ± 0.5 | -13.4 ± 1.3 |
| paPvdQ-C08-HSL | 0 | 7.3 ± 0.6 | -16.5 ± 1.6 |

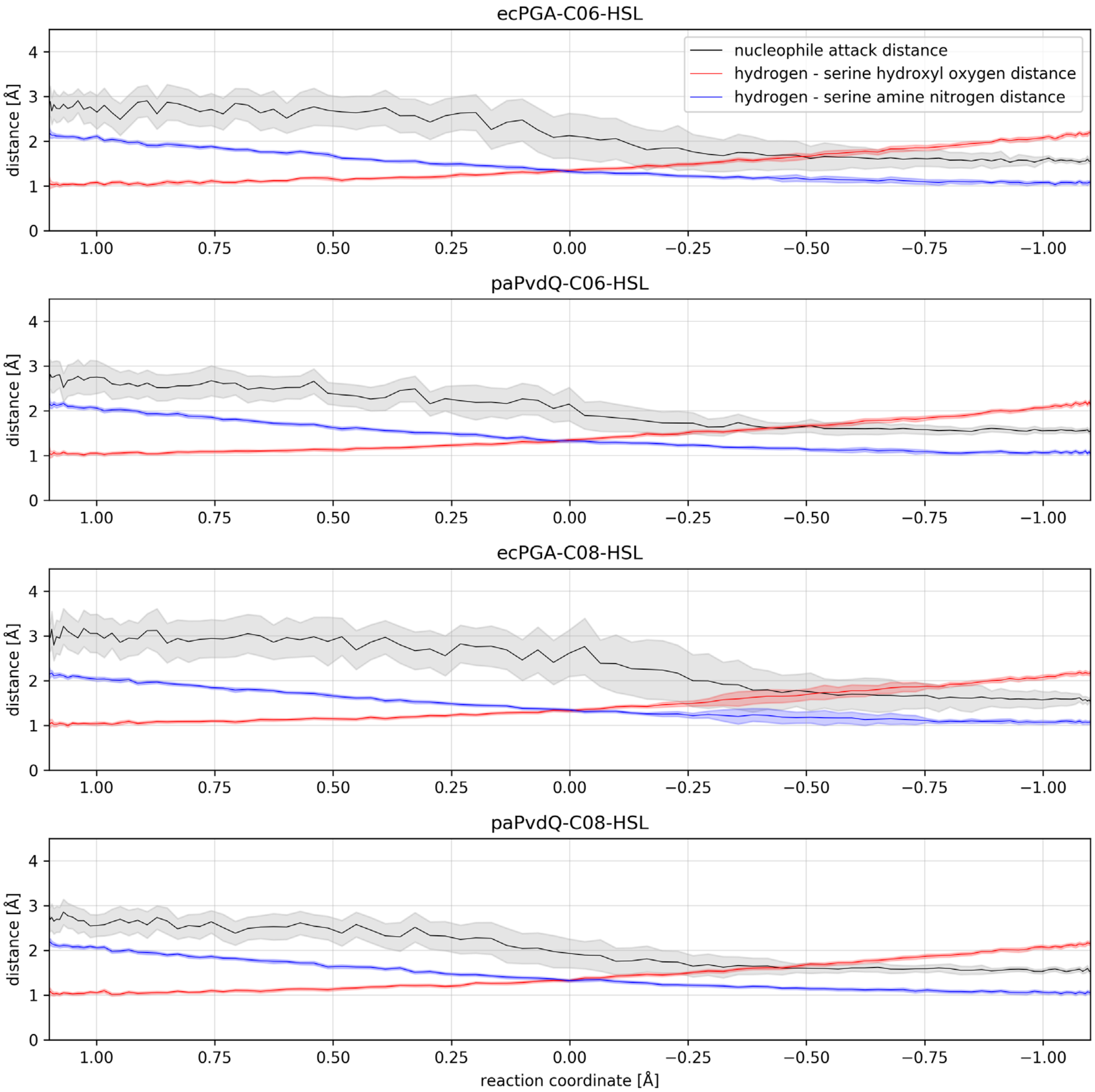

**Figure S16.** Evolution of reaction coordinate (RC) components on the first acylation step for ecPGA and paPvdQ complexes.

**Table S14.** Energetics of the second step of the acylation reaction for ecPGA and paPvdQ complexes.

| complex | energy [kcal/mol] TI | TS2a | TS2b | AE |
|---|---|---|---|---|
| ecPGA-C06-HSL | 0 | 6.8 ± 0.9 | 6.5 ± 1.5 | 2.0 ± 2.0 |
| ecPGA-C08-HSL | 0 | 6.9 ± 0.4 | 6.4 ± 0.8 | -0.9 ± 1.8 |
| paPvdQ-C06-HSL | 0 | 8.5 ± 0.4 | 10.1 ± 1.0 | 4.4 ± 1.2 |
| paPvdQ-C08-HSL | 0 | 8.4 ± 0.5 | 9.2 ± 1.0 | 1.9 ± 1.5 |

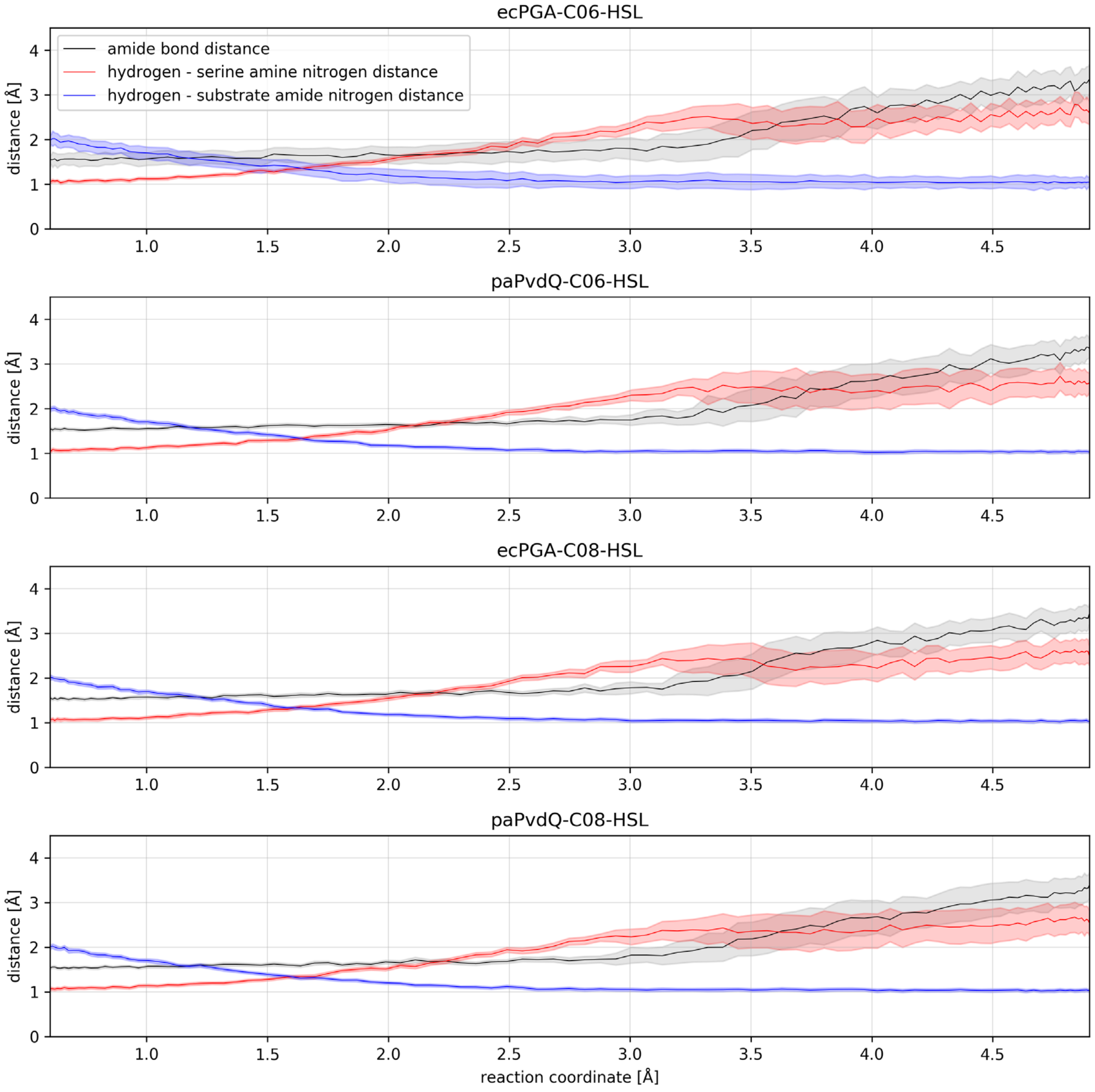

**Figure S17.** Evolution of reaction coordinate (RC) components on the second acylation step for ecPGA and paPvdQ complexes.

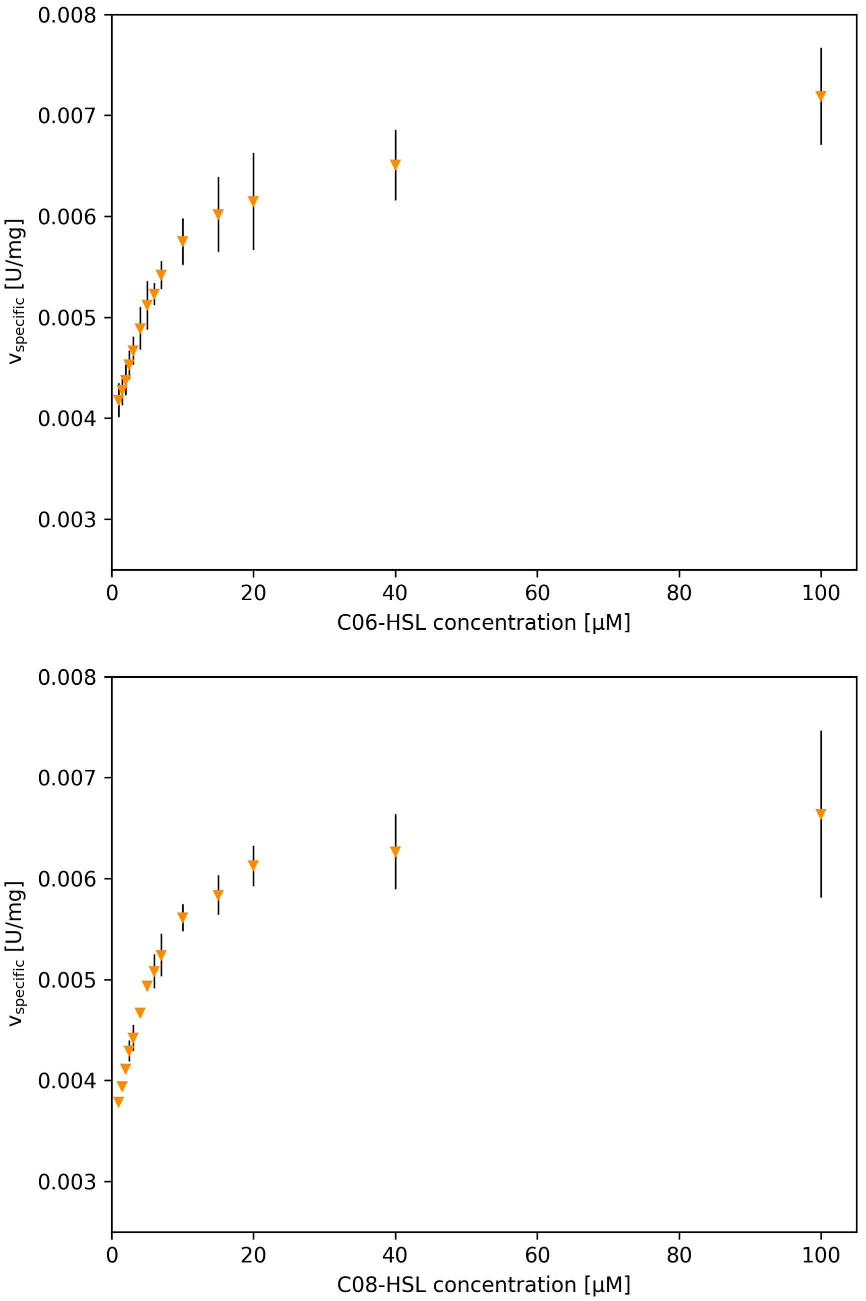

**Figure S18.** ecPGA activity as a function of substrate concentration for conversion of C06-HSL (top) and C08-HSL (bottom).

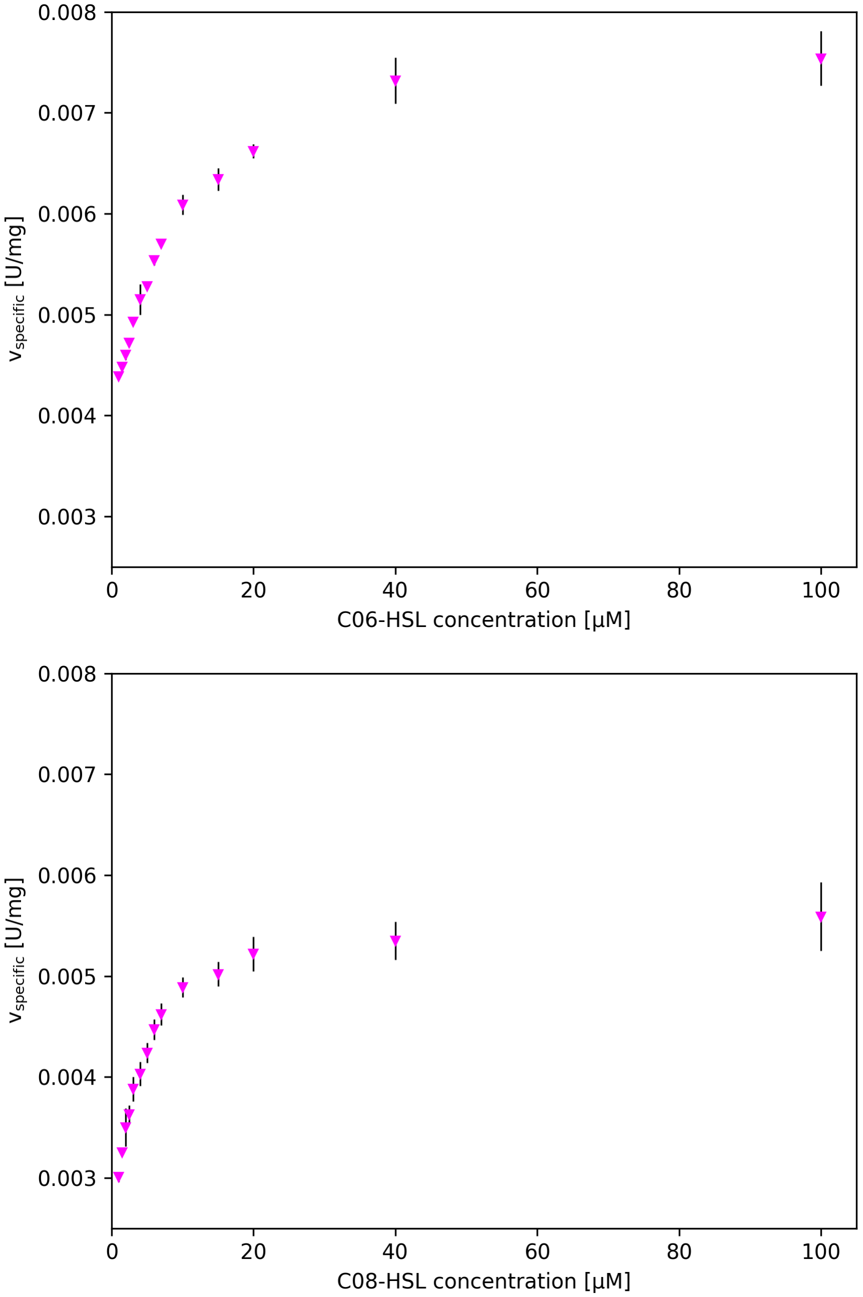

**Figure S19.** aPGA activity as a function of substrate concentration for conversion of C06-HSL (top) and C08-HSL (bottom).

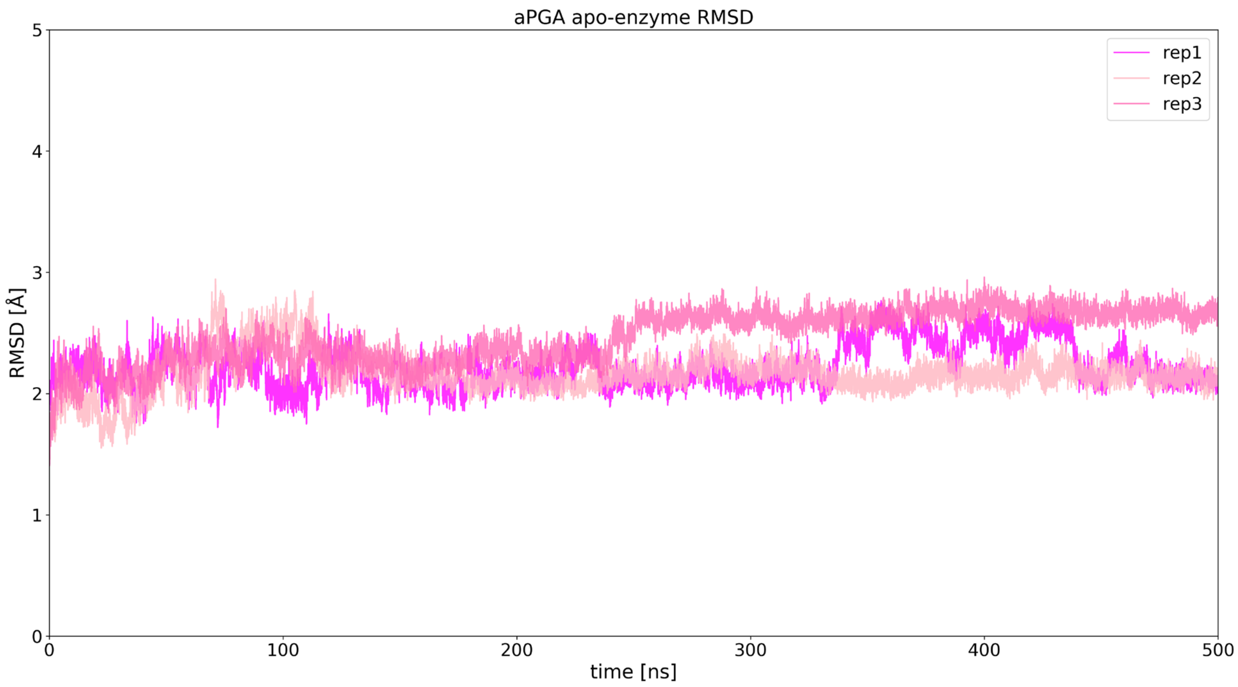

**Figure S20.** RMSD of backbone heavy atoms from three independent simulation replicates of aPGA in the absence of substrate.

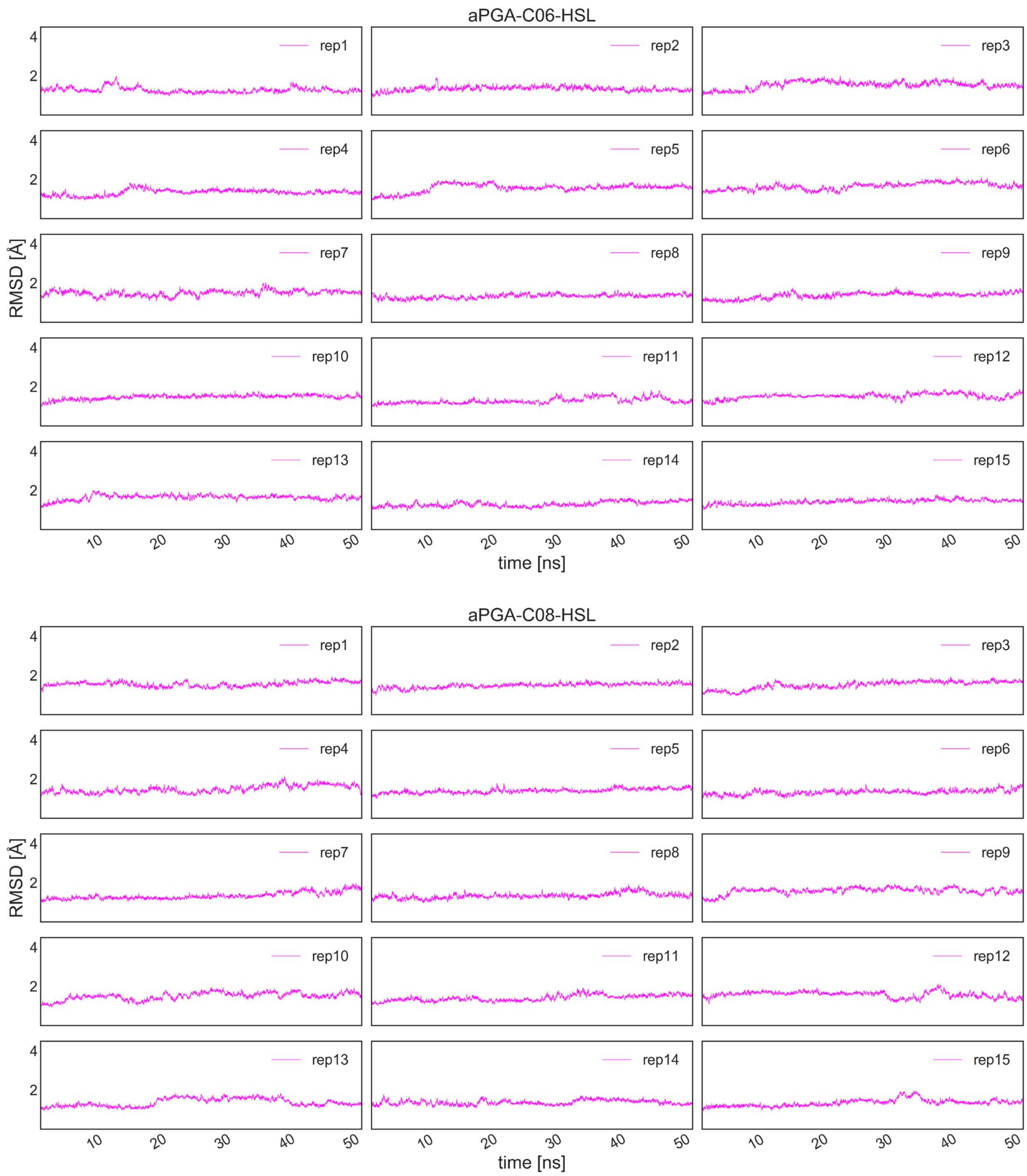

**Figure S21.** RMSD of backbone heavy atoms from 15 independent simulation replicates of aPGA-C06-HSL (top) and aPGA-C08-HSL (bottom) complexes.

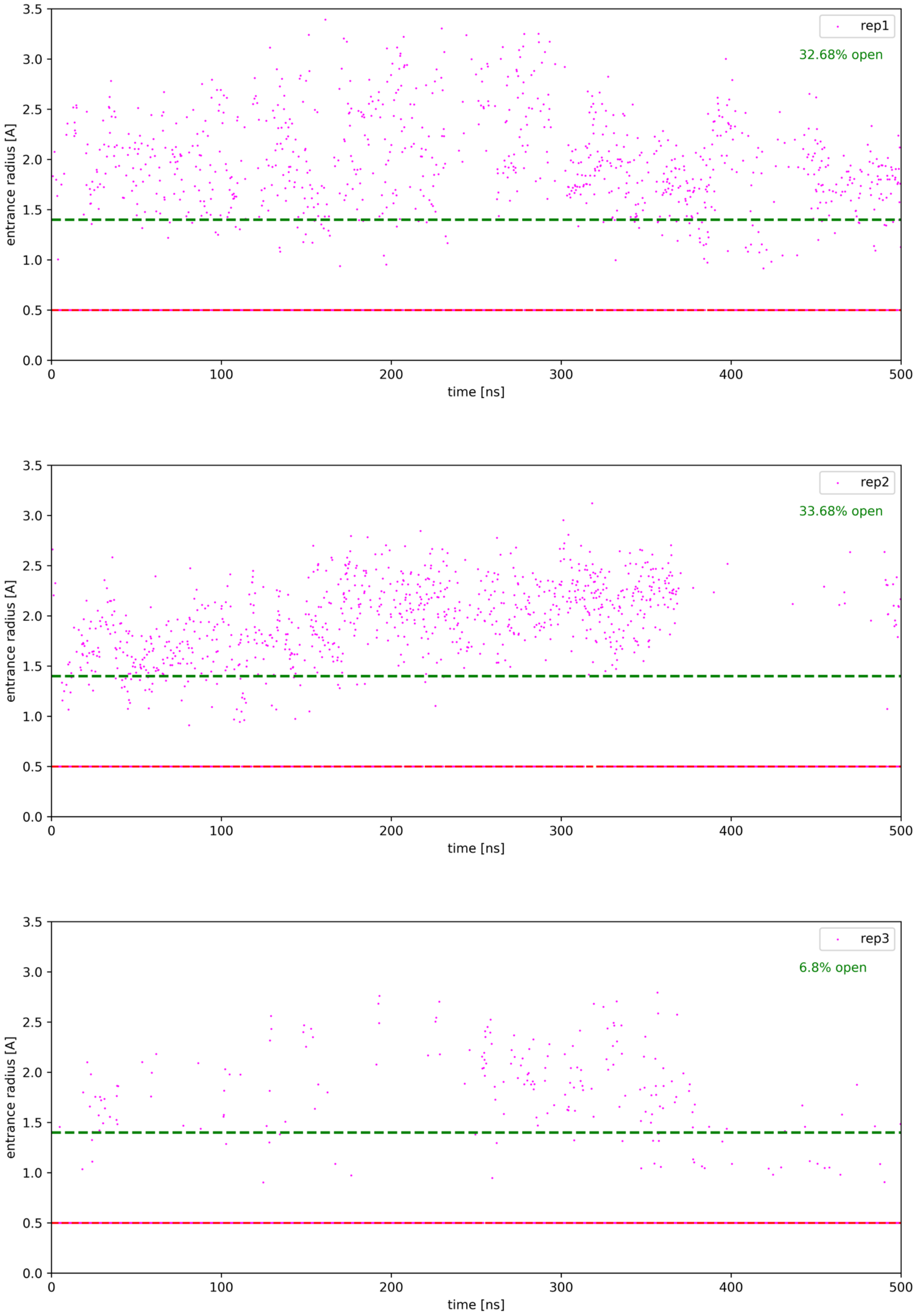

**Figure S22.** Time evolution of the entrance bottleneck to the acyl-binding cavity in aPGA. The open entrance is defined as 1.4 Å (green dashed line). For the frames where CAVER did not detect the pocket, the entrance bottleneck was defined as the searching probe of 0.5 Å (red dashed line).

**Table S15.** Opening of the acyl-binding cavity during free molecular dynamics simulation sufficient for HSLs binding for aPGA enzyme.

| protein | replica | cluster size | analyzed frames | well-shaped* | for docking** |
|---|---|---|---|---|---|
| aPGA | 1 | 896 | 2500 | 142 | 6 |
| | 2 | 902 | 2500 | 299 | 0 |
| | 3 | 206 | 2500 | 31 | 2 |

*Well-shaped binding sites for HSLs defined based on the radius of the entrance bottleneck (narrowest point) > 1.4 Å, minimum depth of the cavity (corresponding to the acyl-chain length) > 5.0 Å, and the real starting point for the calculation is within 1 Å to the desired location.

**Filtering for docking based on a favorable arrangement of catalytic residues promoting productive binding of HSLs defined as: Ser1β-H-hydroxyl → Ser1β-N-amine < 3.0 Å, Asn241β-$N^{\delta}$ → Ala69β-N-backbone < 5.0 Å, Asn241β-$N^{\delta}$ → Ser1β-O-hydroxyl < 5.0 Å, Ala69β-N-backbone → Ser1β-O-hydroxyl < 5.0 Å from PCA analysis of representative conformational states.

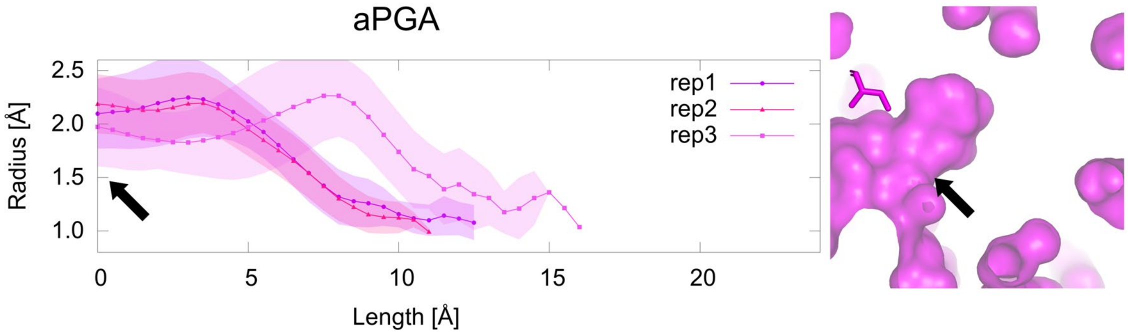

**Figure S23.** Well-shaped binding cavity profiles of aPGA in terms of HSLs binding as defined in **Table S15** (left). The volumetric shape of the cavity is shown as surface representation based on the protein crystal structure (right) for aPGA. The narrowest part of the entrance to the acyl-binding cavity is indicated by black arrows.

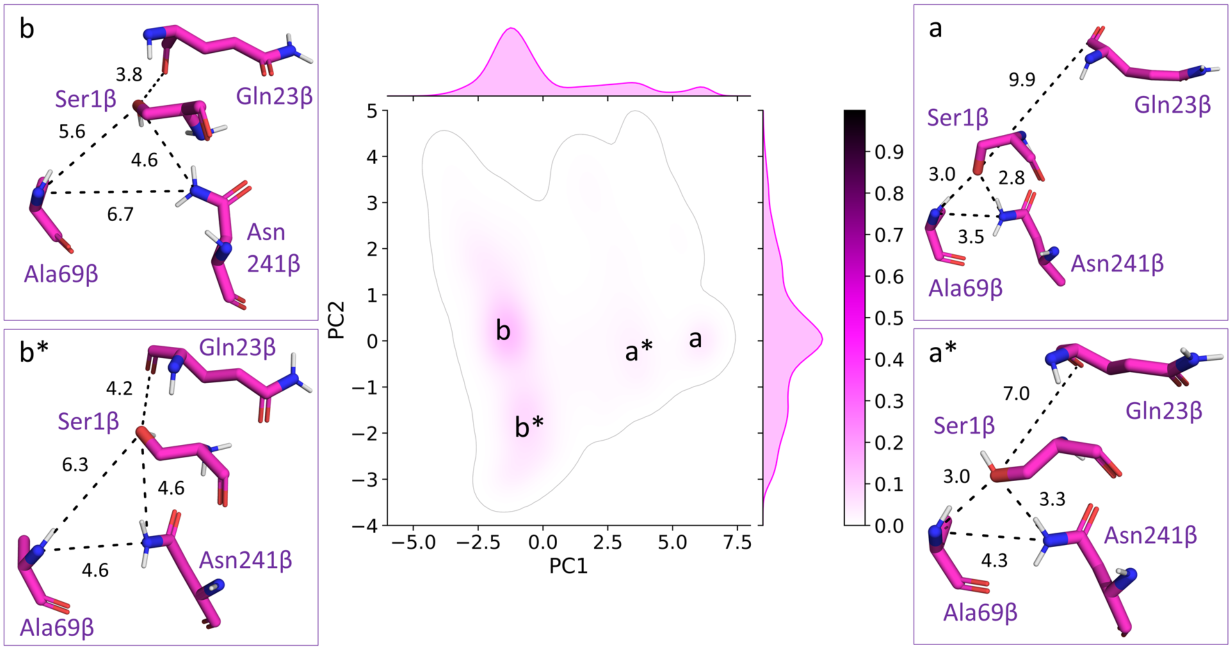

**Figure S24.** Primary conformations adopted by catalytic machinery of aPGA. PCA analysis of the aPGA based on catalytic residues distances obtained from MD simulations without substrates. States b and b* represent catalytic residues favorably pre-organized for productive substrate binding. Remaining states (a and a*) require conformational re-arrangements to enable productive substrate binding.

**Table S16.** The geometry of representative states of catalytic residues extracted from the principal component analysis of free-enzyme molecular dynamics simulations for aPGA.

| Protein | state | Probability density | distance [Å] | | | |
|---|---|---|---|---|---|---|
| | | | AsnN → AlaN[a] | AsnN → SenO[b] | AlaN → SenO[c] | GlnO → SenO[d] |
| aPGA | b | 0.14 | 6.7 | 4.6 | 5.6 | 3.8 |
| | b* | 0.08 | 4.6 | 4.6 | 6.3 | 4.2 |
| | a | 0.04 | 3.5 | 2.8 | 3.0 | 9.9 |
| | a* | 0.03 | 4.3 | 3.3 | 3.0 | 7.0 |

[a] Asn241β-$N^{\delta}$ → Ala69β-N-backbone
[b] Asn241β-$N^{\delta}$ → Ser1β-O-hydroxyl
[c] Ala69β-N-backbone → Ser1β-O-hydroxyl
[d] Gln23β-O-backbone → Ser1β-O-hydroxyl

**Table S17.** Geometries and binding energies of representative protein-ligand complexes obtained in molecular docking experiment with strict filtering based on reaction mechanism for aPGA enzyme.

| protein | ligand | energy [kcal/mol] | distance [Å] | | |
|---|---|---|---|---|---|
| | | | SerO → HSL-C[a] | AlaNH → HSL-O[b] | $AsnNH_2$ → HSL-O[c] |
| aPGA | C06 | -4.0 | 3.0 | 2.6 | 2.1 |
| | C08 | -4.5 | 3.2 | 2.6 | 2.1 |

[a] Ser1β-O-hydroxyl → HSL-C-carbonyl
[b] Ala69β-H-backbone → HSL-O-carbonyl
[c] Asn241β-$N^{\delta}$-$H_x$ (closest) → HSL-O-carbonyl

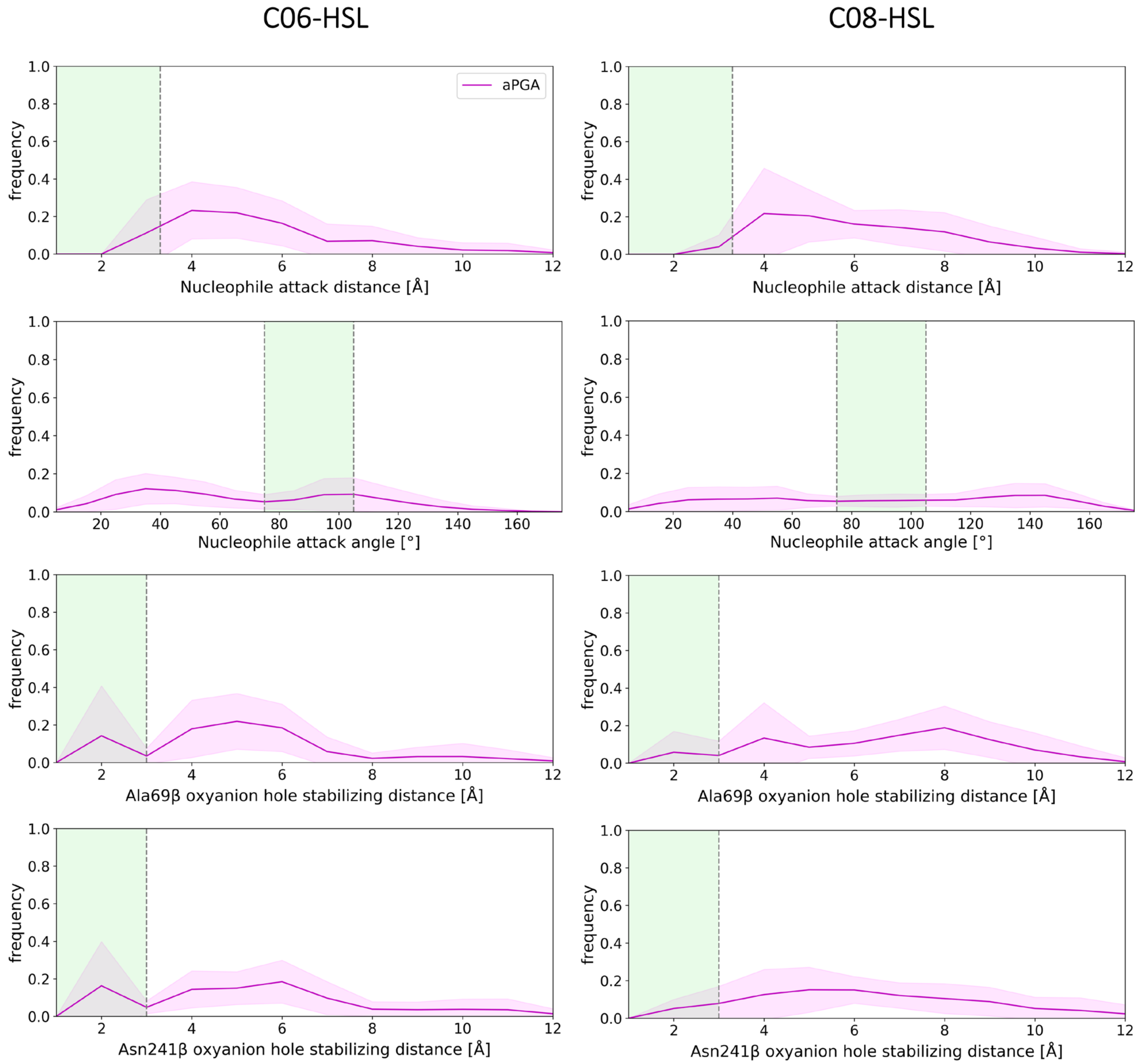

**Figure S25.** Maintenance of the properly stabilized protein-ligand complexes for aPGA. Distribution of nucleophile attack distance (first row), angle (second row), and oxyanion hole stabilizing distances - Ala (third row) and Asn (fourth row) for aPGA enzyme in complex with C06-HSL (left) and C08-HSL (right) from simulations where the average ligand RMSD did not exceed 5.0 Å (**Figure S27**). Regions of the plots highlighted in green indicate the optimal range of the parameter.

**Table S18.** PCA weights for aPGA enzyme's catalytic machinery dynamics in ligand free form.

| PC/distance | SerNH1 → SerO[a] | SerNH2 → SerO[b] | SerH → SerN[c] | AsnN → AlaN[d] | AsnN → SerO[e] | AlaN → SerO[f] | GlnO → SerO[g] |
|---|---|---|---|---|---|---|---|
| | | | | **aPGA** | | | |
| PC1 (0.53) | -0.04 | -0.05 | 0.05 | -0.37 | -0.31 | -0.49 | 0.72 |
| PC2 (0.22) | 0.00 | -0.02 | 0.03 | 0.86 | 0.23 | -0.30 | 0.33 |

[a] Ser1β-H1-amine → Ser1β-O-hydroxyl
[b] Ser1β-H2-amine → Ser1β-O-hydroxyl
[c] Ser1β-H-hydroxyl → Ser1β-N-amine
[d] Asn241β-$N^{\delta}$ → Ala69β-N-backbone
[e] Asn241β-$N^{\delta}$ → Ser1β-O-hydroxyl
[f] Ala69β-N-backbone → Ser1β-O-hydroxyl
[g] Gln23β-O-backbone → Ser1β-O-hydroxyl

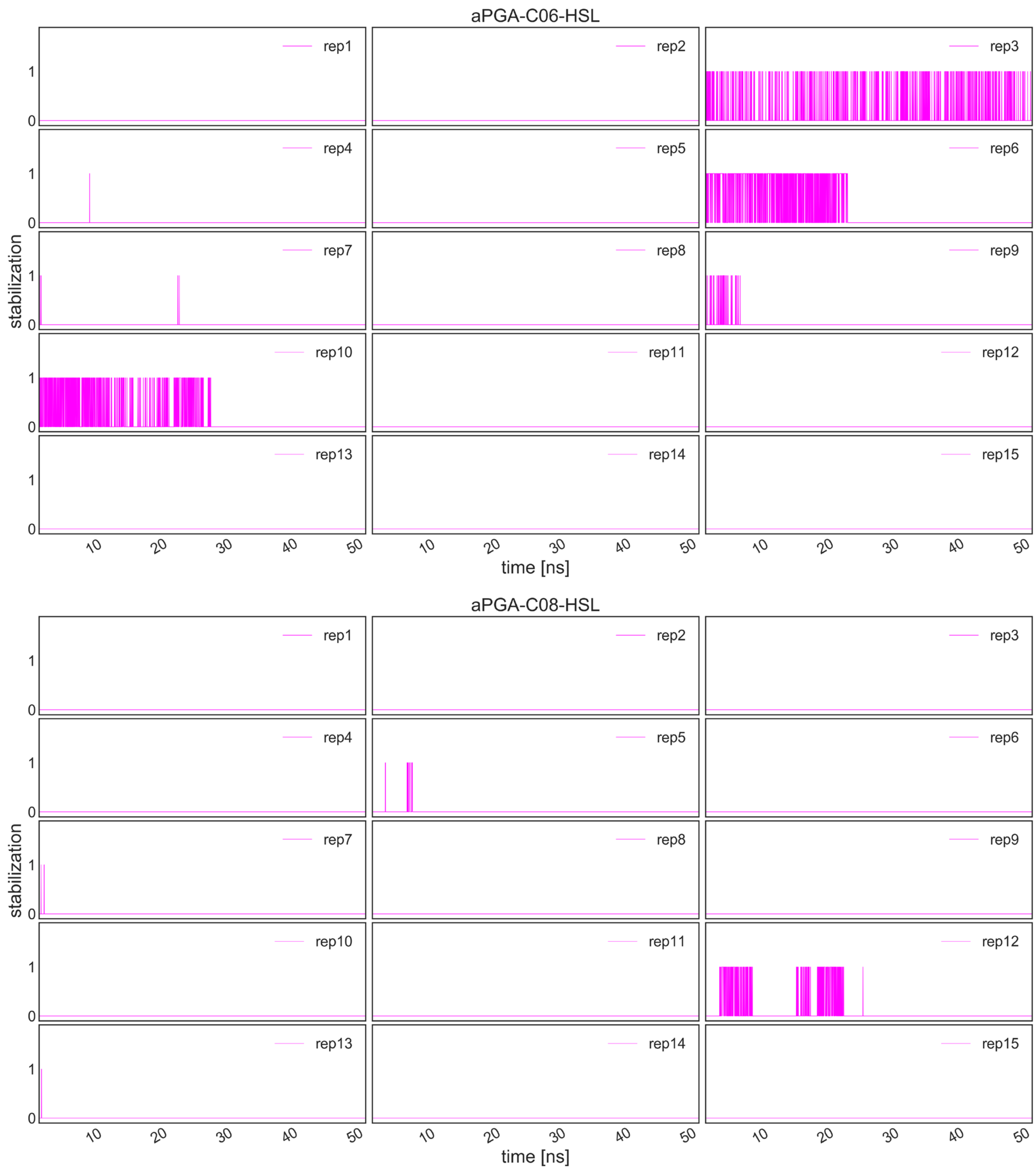

**Figure S26.** Simultaneous stabilization of aPGA-C06-HSL (top) and aPGA-C08-HSL (bottom) complexes in fully productive conformation.

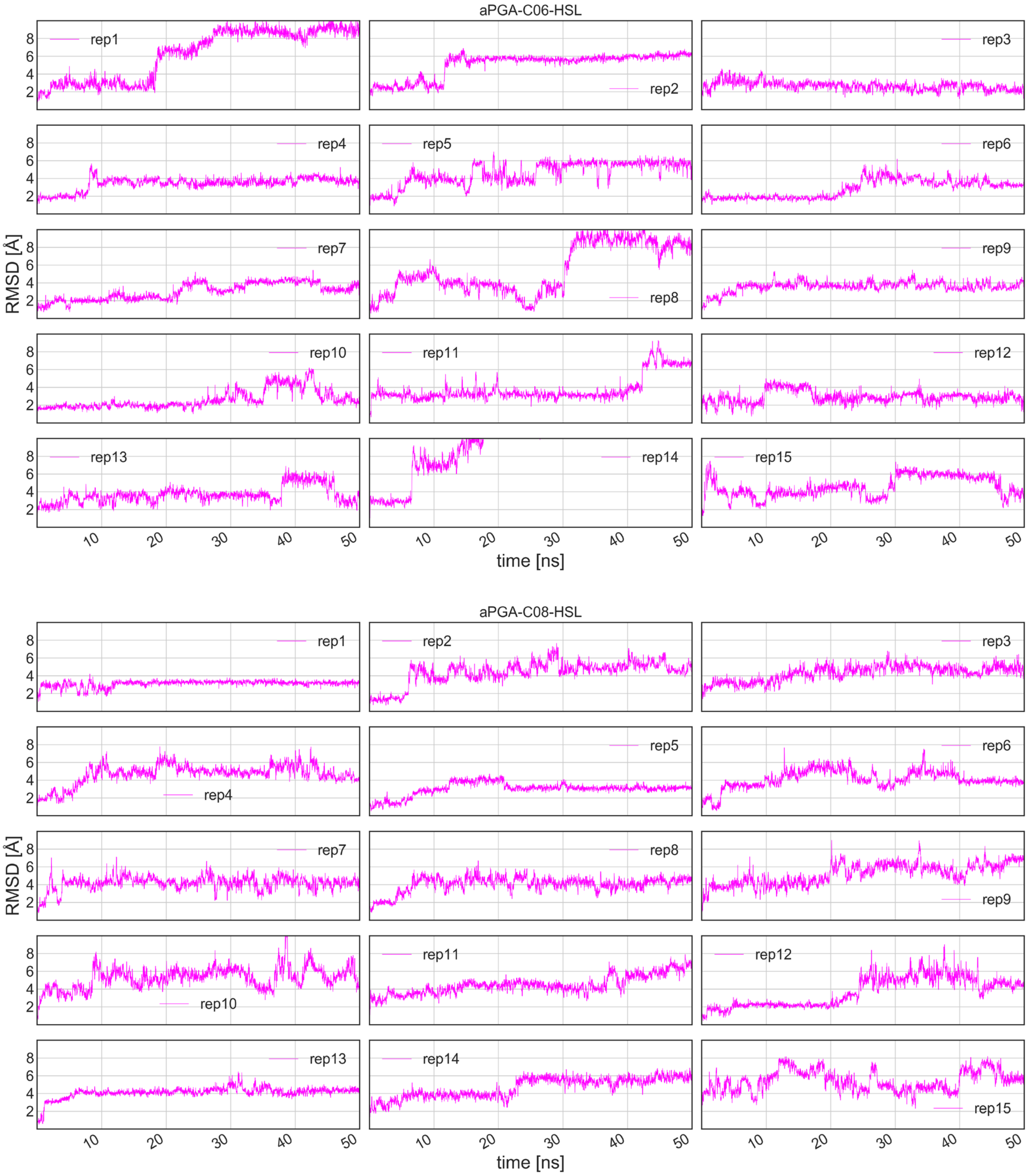

**Figure S27.** Ligand RMSD evolution for aPGA-C06-HSL (top) and aPGA-C08-HSL (bottom) complexes across all protein-ligand complexes MD replicates.

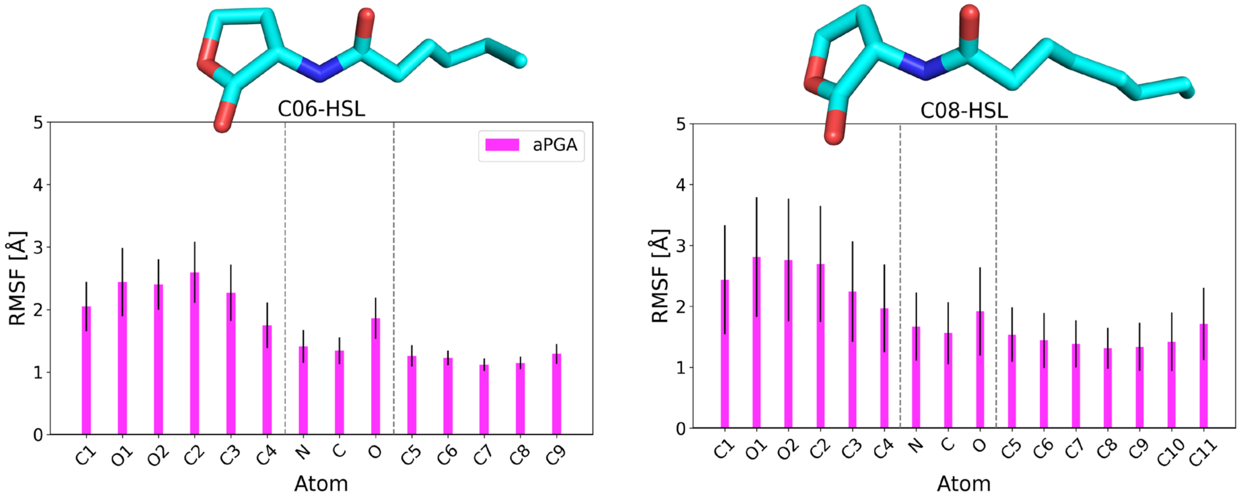

**Figure S28.** The mobility of the HSL atoms during simulations in aPGA complexes. Root-mean-square fluctuation (RMSF) of HSL heavy atoms in simulations where the average ligand RMSD did not exceed 5.0 Å (**Figure S27)**.

**Table S19.** Absolute binding free energies from MMGB/SA calculations for protein-ligand complexes properly stabilized during molecular dynamics simulations of aPGA complexes.

| protein | ligand | binding energy [kcal/mol] |
|---|---|---|
| aPGA | C06-HSL | -31 ± 6 |
| | C08-HSL | -35 ± 3 |

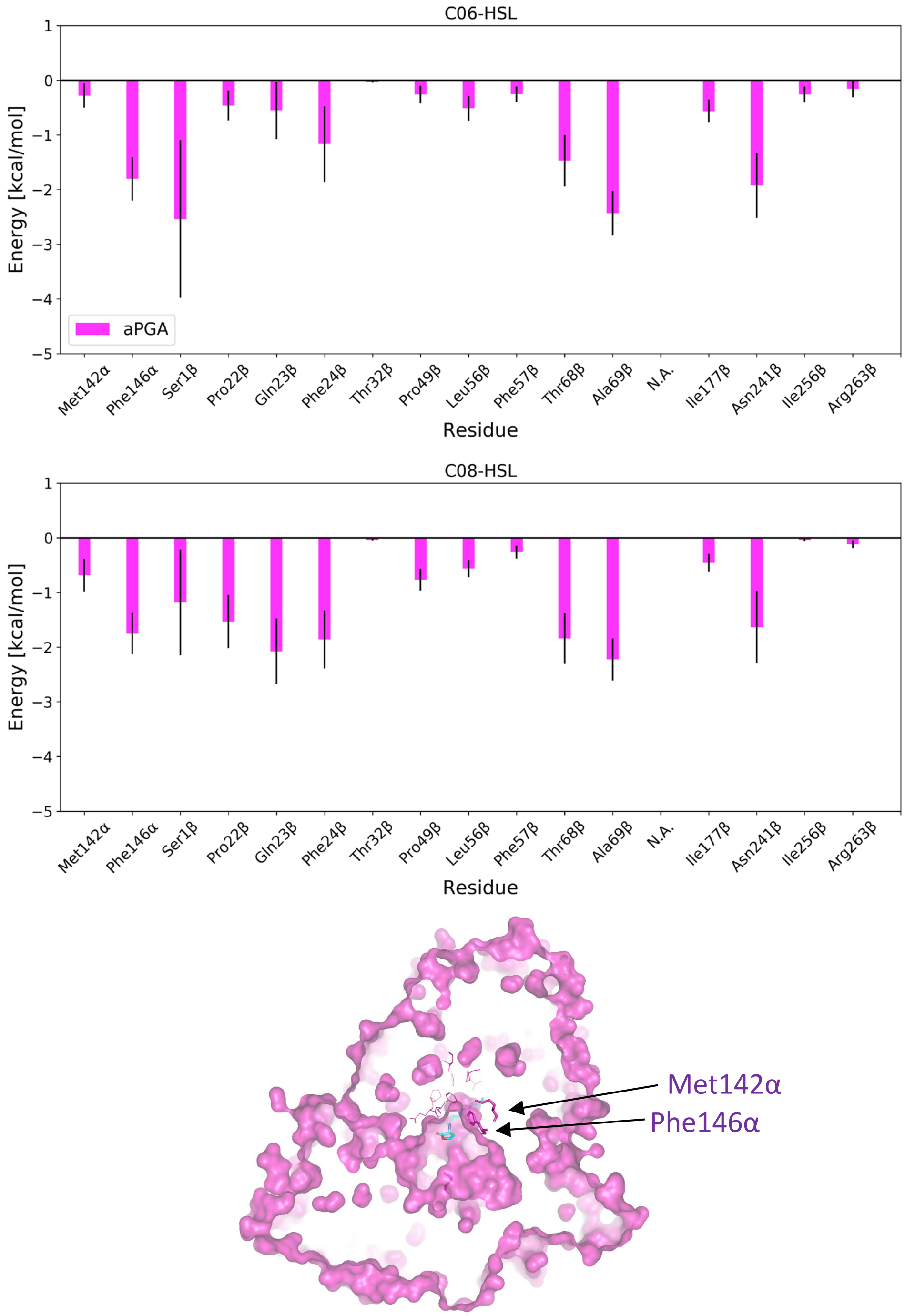

**Figure S29.** Per residue energy decomposition for aPGA-C06-HSL (top) and aPGA-C08-HSL (middle) complexes from MMGB/SA binding energy calculations. In the bottom part of the panel, protein is represented as a surface with all contributing residues shown as lines, while uniquely contributing residues as sticks.

**Table S20.** Configurations of representative structures selected for generation of starting points for steered MD simulations of the acylation process for aPGA complexes.

| complex | distance [Å] | | | | | | | angle [°] |
|---|---|---|---|---|---|---|---|---|
| | SerO → HSL-C*[a] | $AsnNH_2$ → HSL-O**[b] | AlaNH → HSL-O**[c] | GlnO → HSL-H**[d] | SerH → SerN**[e] | $GlnO^{\varepsilon}$ → SerH**[f] | $AsnO^{\delta}$ → SerH**[g] | SerO > HSL-C > HSL-O***[h] |
| aPGA-C06-HSL | 3.0 | 2.3 | 2.0 | 3.0 | 2.6 | 3.0 | 1.9 | 89 |
| aPGA-C08-HSL | 3.4 | 2.1 | 4.6 | 2.2 | 3.4 | 1.7 | 4.0 | 102 |

*desired value below 3.3 Å, preferably close to 3.0 Å
**expected to be within hydrogen bond distance <3.0 Å
***angle should be within the range 75-105°, preferably close to 90°

[a] Ser1β-O-hydroxyl → HSL-C-carbonyl
[b] Asn241β-$N^{\delta}$-$H_x$ (closest) → HSL-O-carbonyl
[c] Ala69β-H-backbone → HSL-O-carbonyl
[d] Gln23β-O-backbone → HSL-H-carbonyl
[e] Ser1β-H-hydroxyl → Ser1β-N-amine
[f] Gln23β-$O^{\varepsilon}$ → Ser1β-H-amine
[g] Asn241β-$N^{\delta}$ → Ser1β-H-amine
[h] Ser1β-O-hydroxyl > HSL-C-carbonyl > HSL-O-carbonyl

**Table S21.** QM/MM MD sampling for aPGA complexes.

| complex | successful complete cycle-repetitions | TI-cumulative time [ns] | AE-cumulative time [ns] | total sMD time [ns] |
|---|---|---|---|---|
| aPGA-C06-HSL | 210 | 7.4 | 14.7 | 22.1 |
| aPGA-C08-HSL | 240 | 8.4 | 16.8 | 25.2 |

**Table S22.** Detailed average geometries of particular states ensembles during acylation reaction for aPGA complexes.

| complex | state | mean / std | distance [Å]: SerO → HSL-C[a] | AlaNH → HSL-O[b] | AsnNH2 → HSL-O[c] | Gln → HSL-H[d] | HSL-C → HSL-N[e] | ArgH → HSL-O[f] | ArgC →SerN[g] | angle [°]: ArgC > SerN > HSL-N[h] | HSL-N > SerN > ArgN1 > ArgN2[i] |
|---|---|---|---|---|---|---|---|---|---|---|---|
| aPGA-C06-HSL | MC | mean | 2.9 | 2.0 | 2.2 | 2.4 | 1.4 | 3.8 | 5.2 | 103 | -61 |
| | | std | 0.3 | 0.2 | 0.3 | 0.6 | 0.1* | 1.0 | 0.4 | 9 | 13 |
| | TS1 | mean | 2.7 | 2.0 | 2.3 | 2.5 | 1.4 | 3.7 | 5.4 | 104 | -64 |
| | | std | 0.4 | 0.2 | 0.4 | 0.6 | 0.1* | 1.1 | 0.4 | 9 | 13 |
| | TI | mean | 1.5 | 2.0 | 1.8 | 2.3 | 1.5 | 3.2 | 5.3 | 113 | -74 |
| | | std | 0.1 | 0.2 | 0.2 | 0.4 | 0.1* | 0.9 | 0.4 | 8 | 19 |
| | TS2a | mean | 1.5 | 2.0 | 1.8 | 2.0 | 1.7 | 2.4 | 5.3 | 107 | -75 |
| | | std | 0.1* | 0.2 | 0.2 | 0.2 | 0.2 | 0.7 | 0.3 | 6 | 18 |
| | TS2b | mean | 1.4 | 2.1 | 2.0 | 2.0 | 2.0 | 2.8 | 5.2 | 101 | -60 |
| | | std | 0.1* | 0.2 | 0.2 | 0.2 | 0.3 | 0.9 | 0.4 | 10 | 20 |
| | AE | mean | 1.4 | 2.2 | 2.2 | 2.3 | 3.1 | 3.0 | 5.3 | 85 | -61 |
| | | std | 0.1* | 0.3 | 0.3 | 0.4 | 0.2 | 1.1 | 0.4 | 10 | 29 |
| aPGA-C08-HSL | MC | mean | 2.7 | 2.0 | 2.2 | 2.2 | 1.4 | 3.0 | 4.8 | 103 | -60 |
| | | std | 0.3 | 0.2 | 0.3 | 0.6 | 0.1* | 0.9 | 0.4 | 8 | 19 |
| | TS1 | mean | 2.4 | 2.0 | 2.1 | 2.2 | 1.4 | 3.0 | 5.2 | 104 | -66 |
| | | std | 0.5 | 0.2 | 0.3 | 0.4 | 0.1* | 1.0 | 0.4 | 11 | 23 |
| | TI | mean | 1.5 | 1.9 | 1.7 | 2.5 | 1.5 | 2.6 | 5.1 | 114 | -73 |
| | | std | 0.1* | 0.2 | 0.2 | 0.6 | 0.1* | 0.7 | 0.3 | 7 | 29 |
| | TS2a | mean | 1.5 | 2.0 | 1.9 | 2.1 | 1.7 | 2.5 | 5.0 | 109 | -76 |
| | | std | 0.1* | 0.2 | 0.2 | 0.3 | 0.1 | 0.6 | 0.3 | 6 | 24 |
| | TS2b | mean | 1.4 | 2.0 | 2.0 | 2.2 | 2.0 | 2.5 | 5.1 | 102 | -70 |
| | | std | 0.1* | 0.2 | 0.2 | 0.4 | 0.3 | 0.7 | 0.3 | 8 | 26 |
| | AE | mean | 1.4 | 2.0 | 2.1 | 2.5 | 3.1 | 2.7 | 5.0 | 89 | -69 |
| | | std | 0.1* | 0.2 | 0.3 | 0.5 | 0.2 | 1.0 | 0.4 | 11 | 32 |

* std rounded up to meaningful magnitudes and maintain consistent formatting; [a] Ser1β-O-hydroxyl → HSL-C-carbonyl; [b] Asn241β-N$^{\delta}$-Hx (closest) → HSL-O-carbonyl; [c] Ala69β-H-backbone → HSL-O-carbonyl; [d] Gln23β-O-backbone → HSL-H-carbonyl; [e] HSL-C-carbonyl → HSL-N; [f] Arg263β-$NH_x$ (guanidyl closest) → HSL-O-lactone; [g] Arg263β-C-guanidyl → Ser1β-N-amine; [h] Arg263β-C-guanidyl > Ser1β-N-amine > HSL-N; [i] Ser1β-N-amine > HSL-N > Arg263β-N1-guanidyl > Arg263β-N2-guanidyl

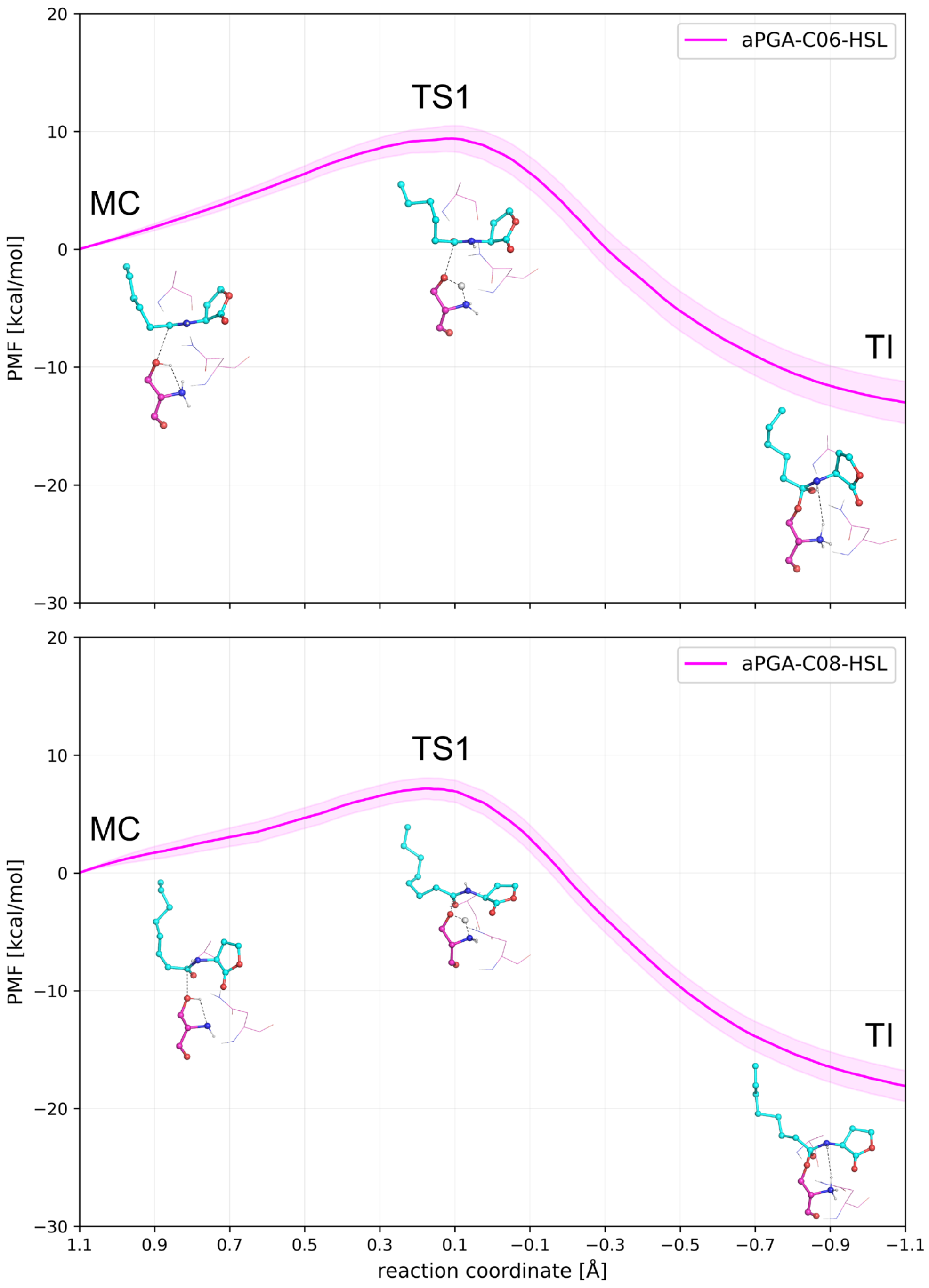

**Figure S30.** Potential of mean force profiles from steered MD simulations of the tetrahedral intermediate formation step for aPGA enzyme in complex with C06-HSL (upper panel) and C08-HSL (lower panel). Starting from the MC (RC 1.1 Å), systems proceed towards TS1 (RC ~ 0.2 Å), where the catalytic serine hydroxyl hydrogen was shared between its hydroxyl and amine group, which was synchronized with the nucleophile attack of the hydroxyl oxygen to the substrate carbonyl carbon and resulted in TI formation (RC -1.1 Å).

**Table S23.** Energetics of the first step of the acylation reaction for aPGA complexes.

| complex | energy [kcal/mol] | | |
|---|---|---|---|
| | MC | TS1 | TI |
| aPGA-C06 | 0 | 9.4 ± 1.1 | -13.0 ± 1.8 |
| aPGA-C08 | 0 | 7.2 ± 0.9 | -18.1 ± 1.3 |

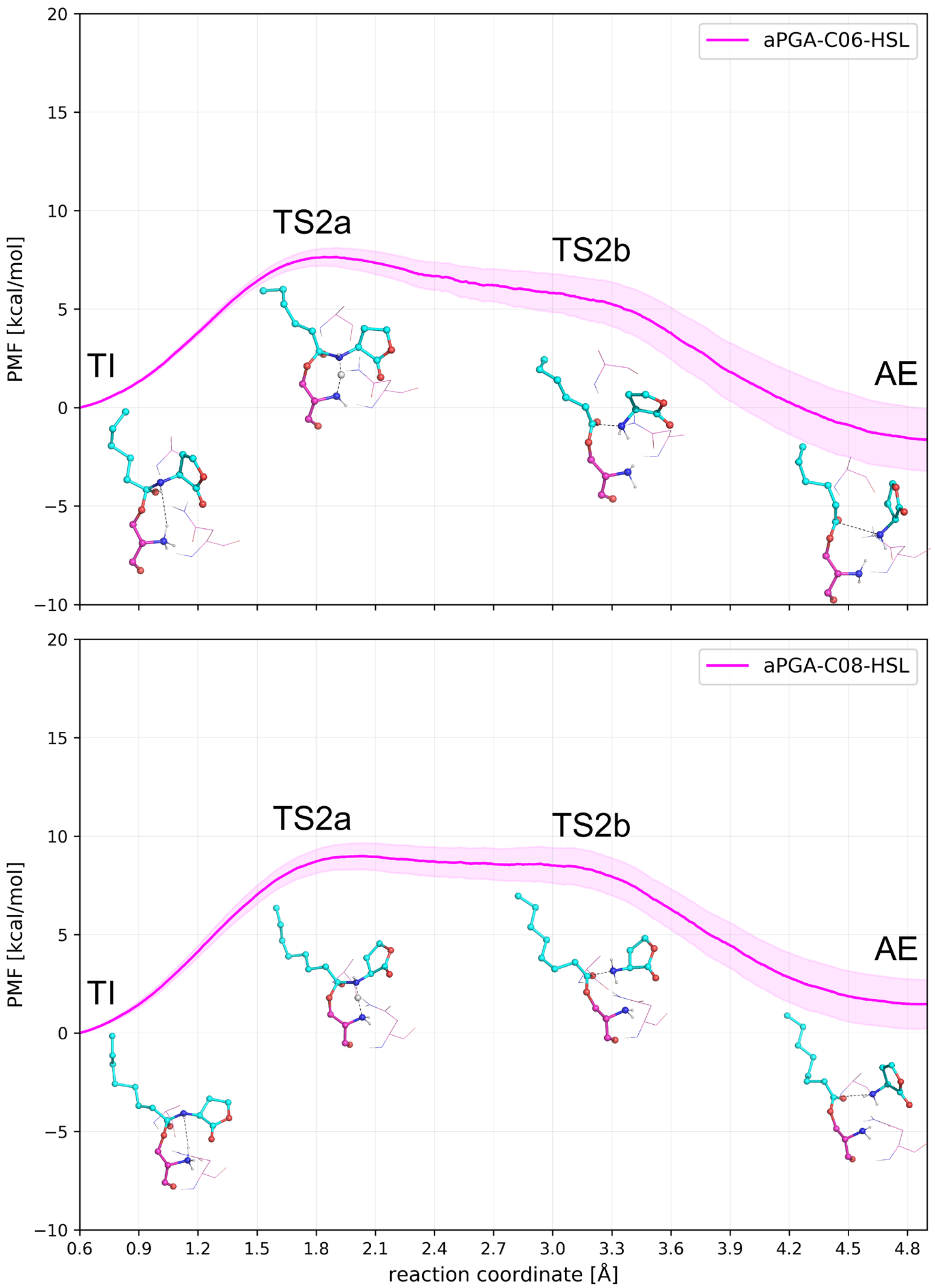

**Figure S31.** Potential of mean force profiles from steered MD simulations of the acyl-enzyme formation step for aPGA enzyme with C06-HSL (upper panel) and C08-HSL (lower panel). Starting from the TI (RC 0.6 Å), systems follow towards TS2a (RC ~ 1.9 Å), corresponding to the proton shared between the catalytic serine amine group and the nitrogen of the leaving group. Further, the proton was fully transferred to the leaving group nitrogen and the amide bond was broken, accounting for TS2b (RC ~ 3.3 Å). Finally, the first reaction product, homoserine lactone, was released from the active site and resulted in AE formation (RC 4.9 Å).

**Table S24.** Energetics of the second step of the acylation reaction for aPGA complexes.

| complex | energy [kcal/mol] | | | |
|---|---|---|---|---|
| | TI | TS2a | TS2b | AE |
| aPGA-C06 | 0 | 7.7 ± 0.5 | 5.1 ± 1.2 | -1.6 ± 1.6 |
| aPGA-C08 | 0 | 9.0 ± 0.7 | 8.3 ± 1.0 | 1.5 ± 1.3 |

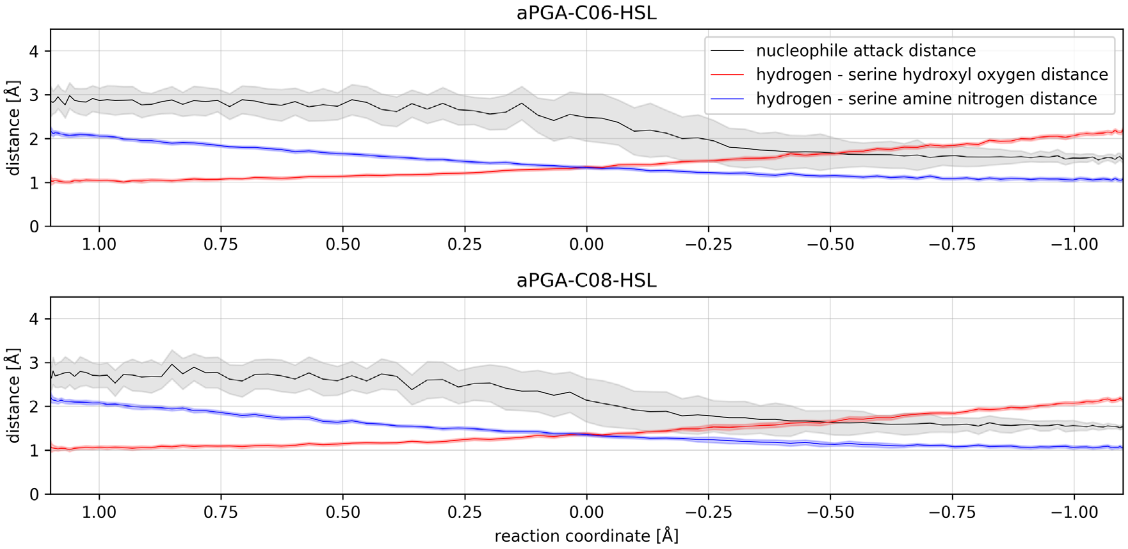

**Figure S32.** Evolution of reaction coordinate (RC) components on the first acylation step for aPGA complexes.

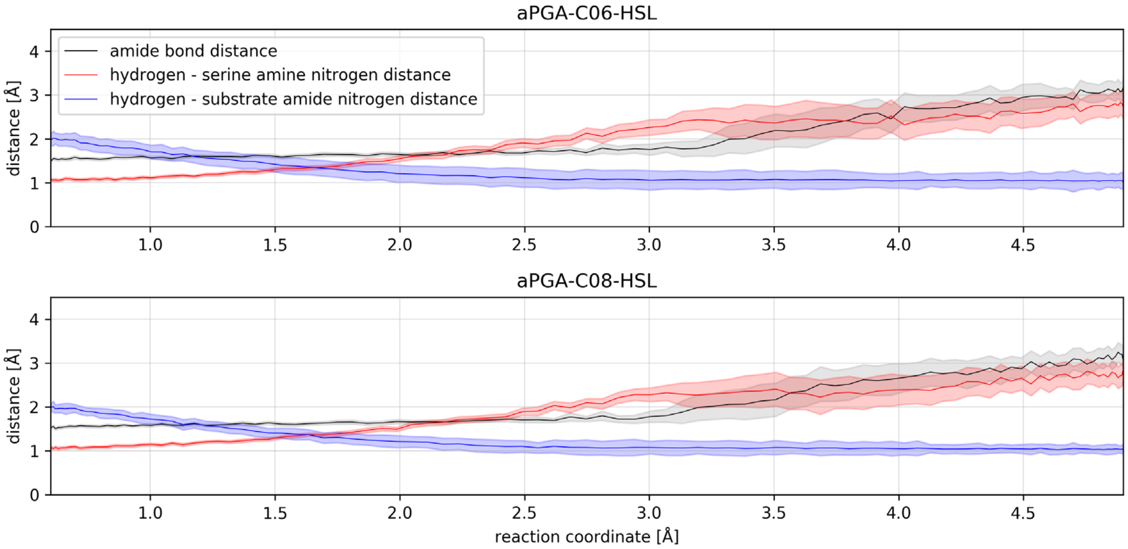

**Figure S33.** Evolution of reaction coordinate (RC) components on the second acylation step for aPGA complexes.

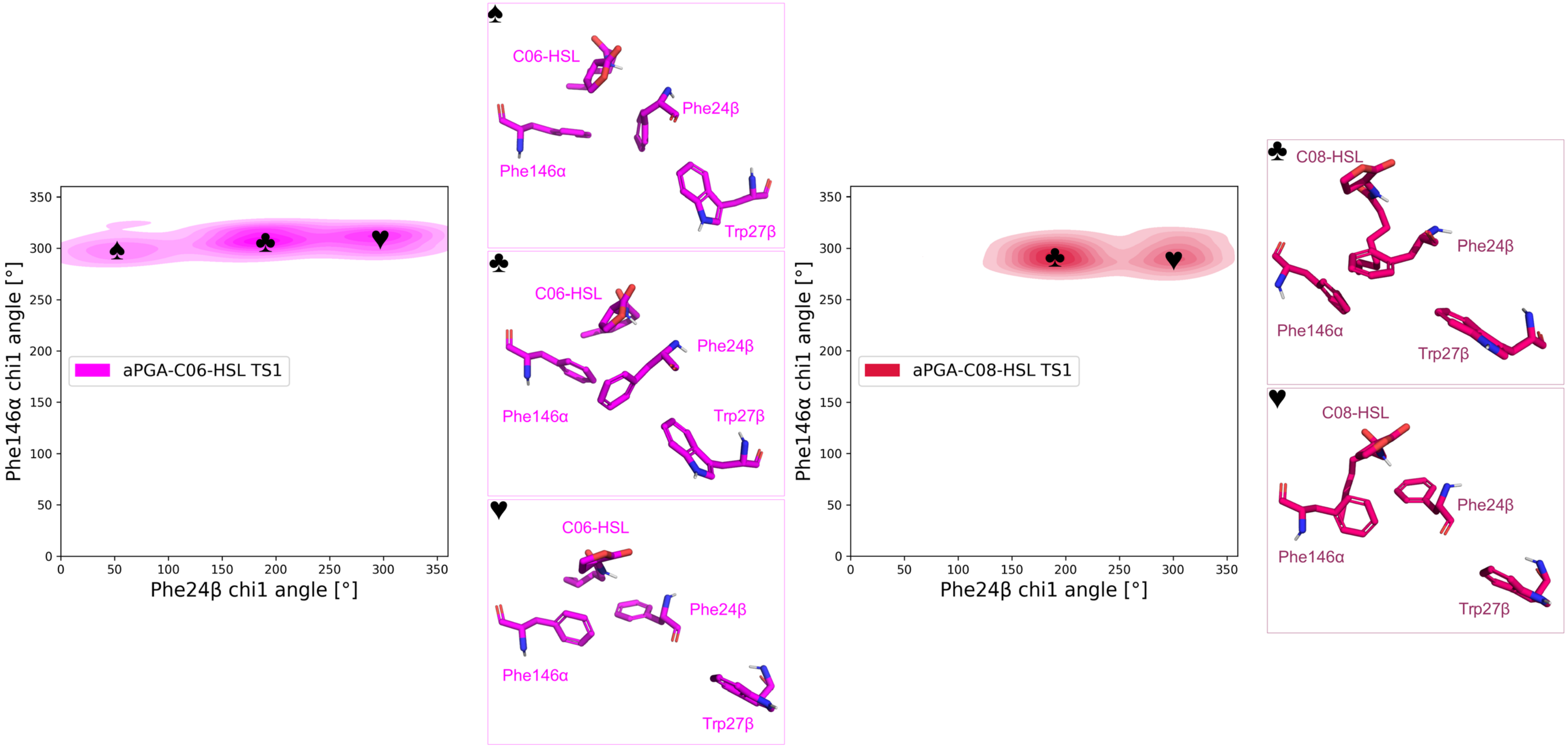

**Figure S34.** Dynamics of the gate to the acyl-binding cavity in aPGA. Due to the difference in position 27β (Trp), the Phe24β was more mobile in aPGA, which affected the dynamics of Phe146α. This resulted in the opening of the side-pocket in aPGA and enabled deeper penetration of the ligand compared to ecPGA.

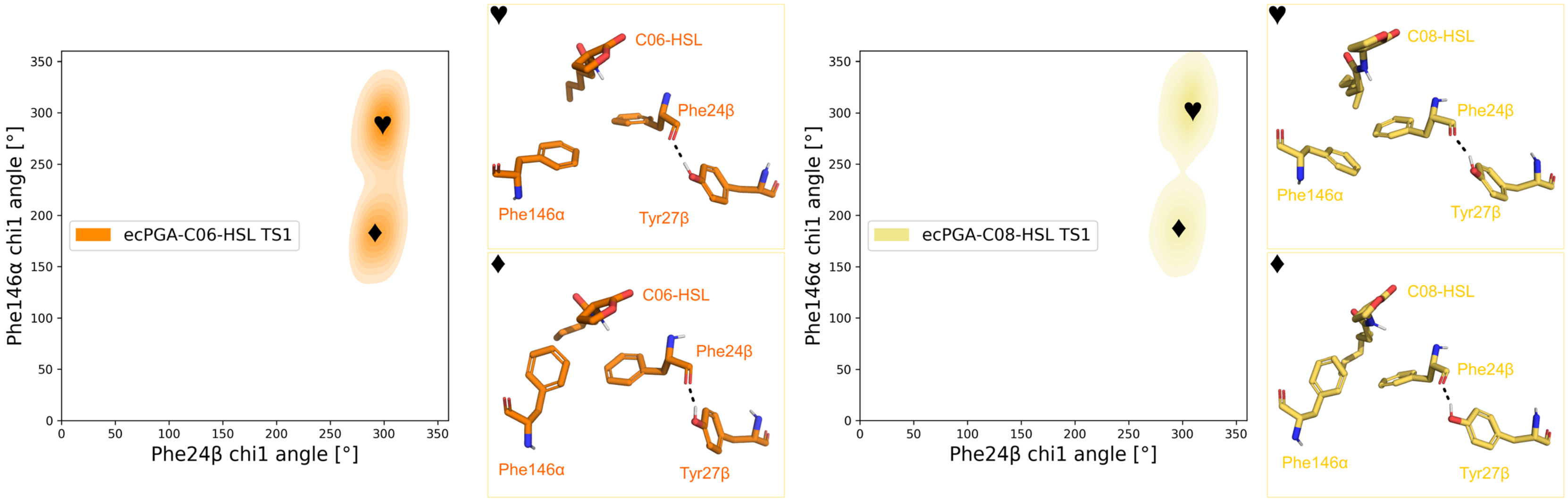

**Figure S35.** Dynamics of the gate to the acyl-binding cavity in ecPGA. Due to the difference in position 27β (Tyr), the Phe24β presented reduced dynamics, which prevented the opening of the side-pocket necessary for binding the longer substrate (C08-HSL).

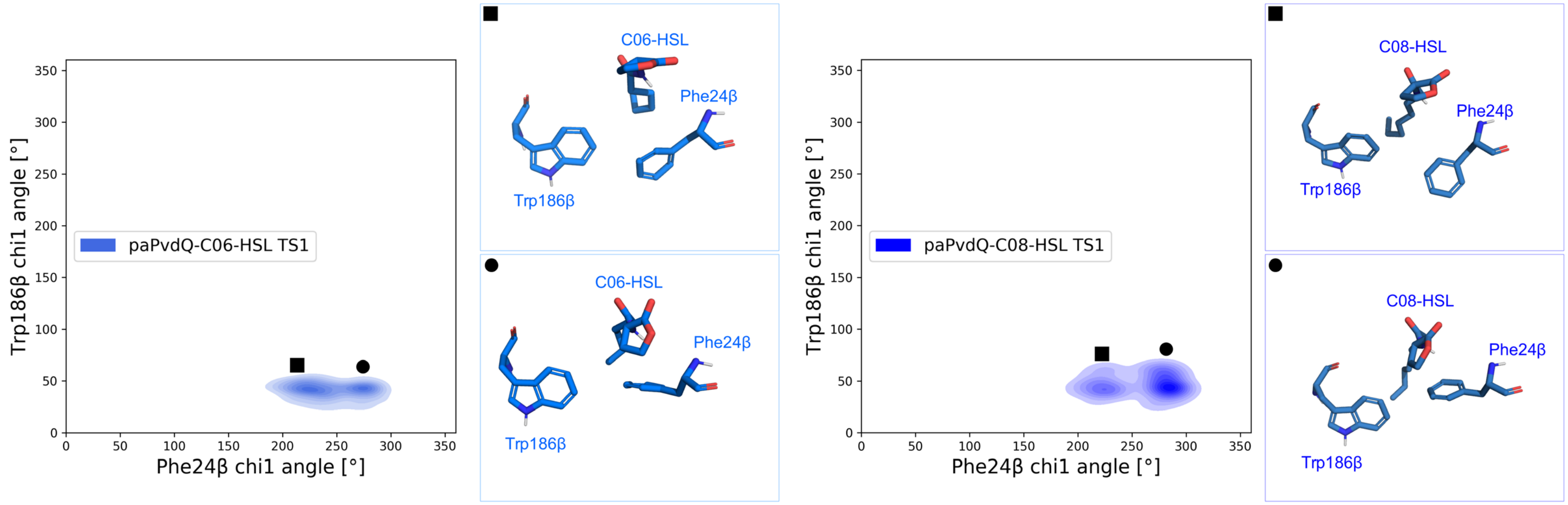

**Figure S36.** Dynamics of the gate to the acyl-binding cavity in paPvdQ. Here the gating residues maintained stable interactions and exhibited only small fluctuations between two states differing in relative Phe24β orientation. This resulted in a persistent geometry of the acyl-binding cavity.

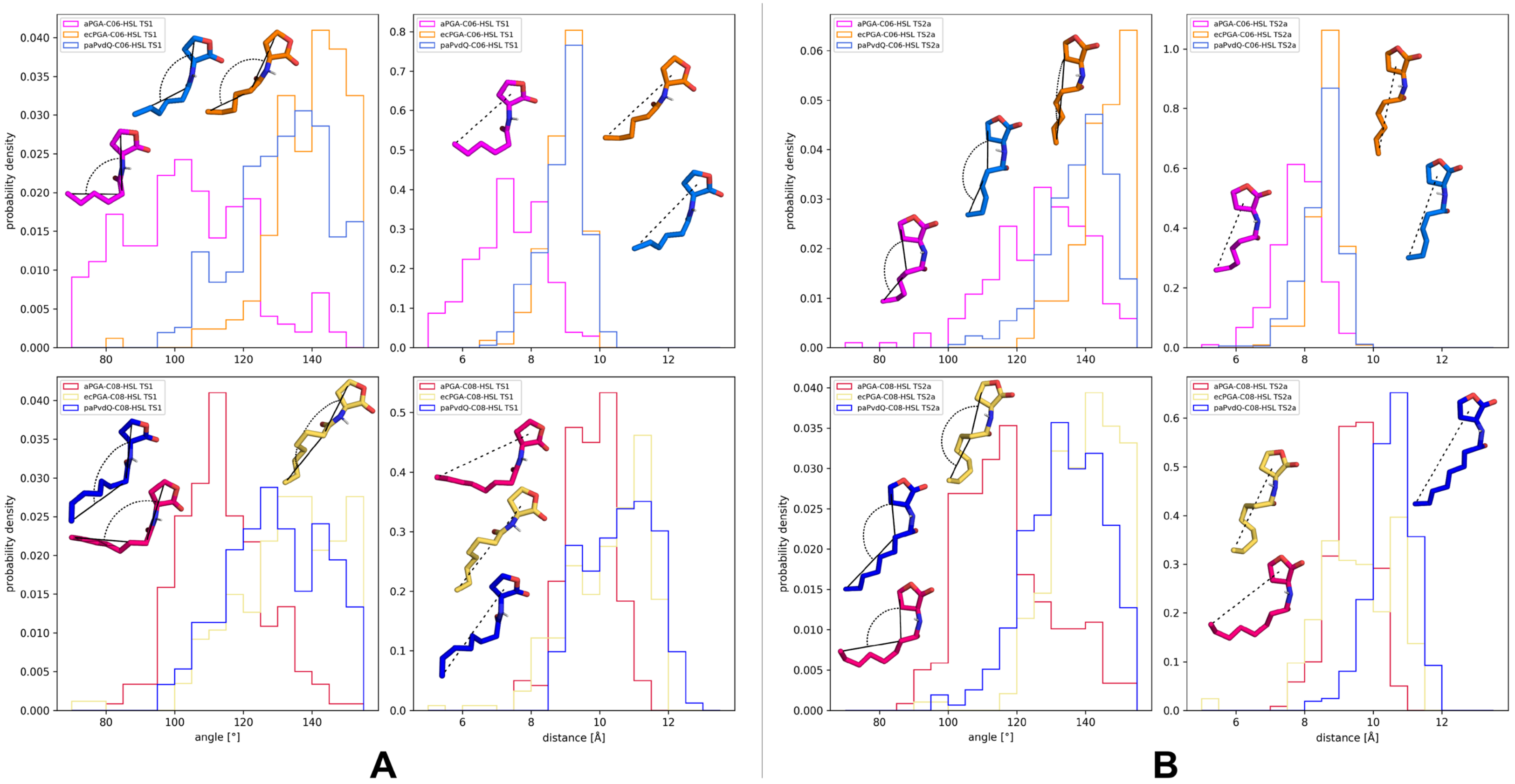

**Figure S37.** Distributions of the HSLs curvature and their acyl-chain extension at TS1 and TS2a of the reaction. The ligand curvature was measured as an angle between the furthest lactone ring carbon, an acyl-chain carbon atom neighboring amide bond, and the last carbon of the acyl-chain, while the acyl-chain extension was represented by the distance between the last carbon of the chain and the center of mass of the lactone ring. (A) in TS1, more pronounced dynamics of the acyl-binding cavity resulted in the opening of a distinct side-pocket capable of binding HSLs with bent acyl-chains favorable for the longer substrate (C08-HSL). However, the extra space available within this side-pocket was responsible for the suboptimal reactivity of the shorter substrate (C06-HSL) bound too deep in the cavity. Conversely, ecPGA unable to open the side-pocket had to accommodate both HSLs in an approximately extended conformation. Similarly, the relatively static and elongated acyl-binding cavity of paPvdQ favored extended conformations of both HSLs. (B) in TS2, only C08-HCL tended to exhibit a markedly curved acyl-chain due to its fit into the side-pocket formed in the acyl-binding cavity of aPGA.

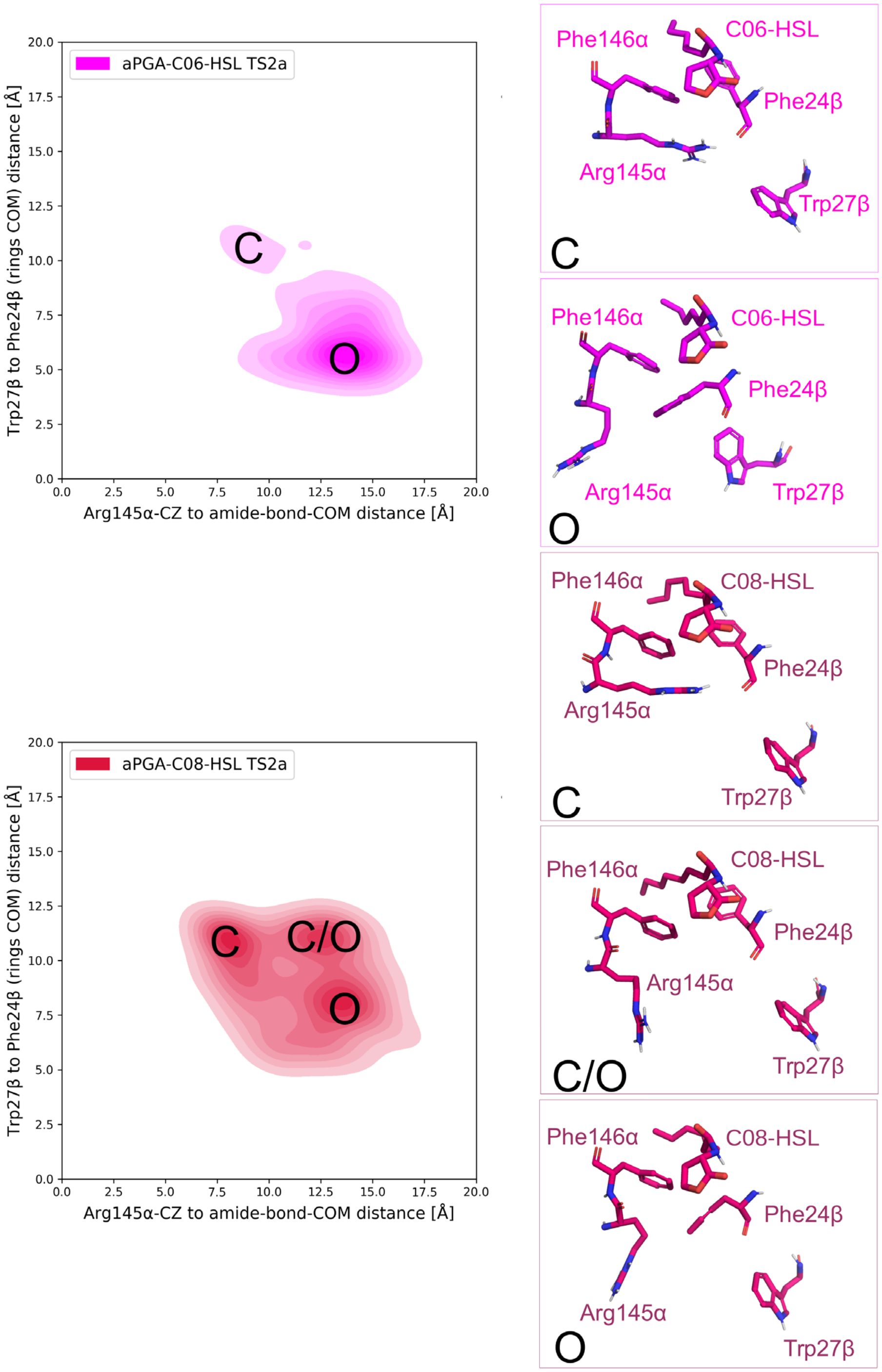

**Figure S38.** Dynamics of the gate controlling overall access to the active site of aPGA. In the aPGA-C08-HSL complex, we observed promoted closed conformation (C) defined by Arg145α guanidyl moiety being close to the backbone oxygen of Phe24β residue and the amide bond of the substrate, resulting in less favorable energetics for this complex at the second acylation step (TS2a).

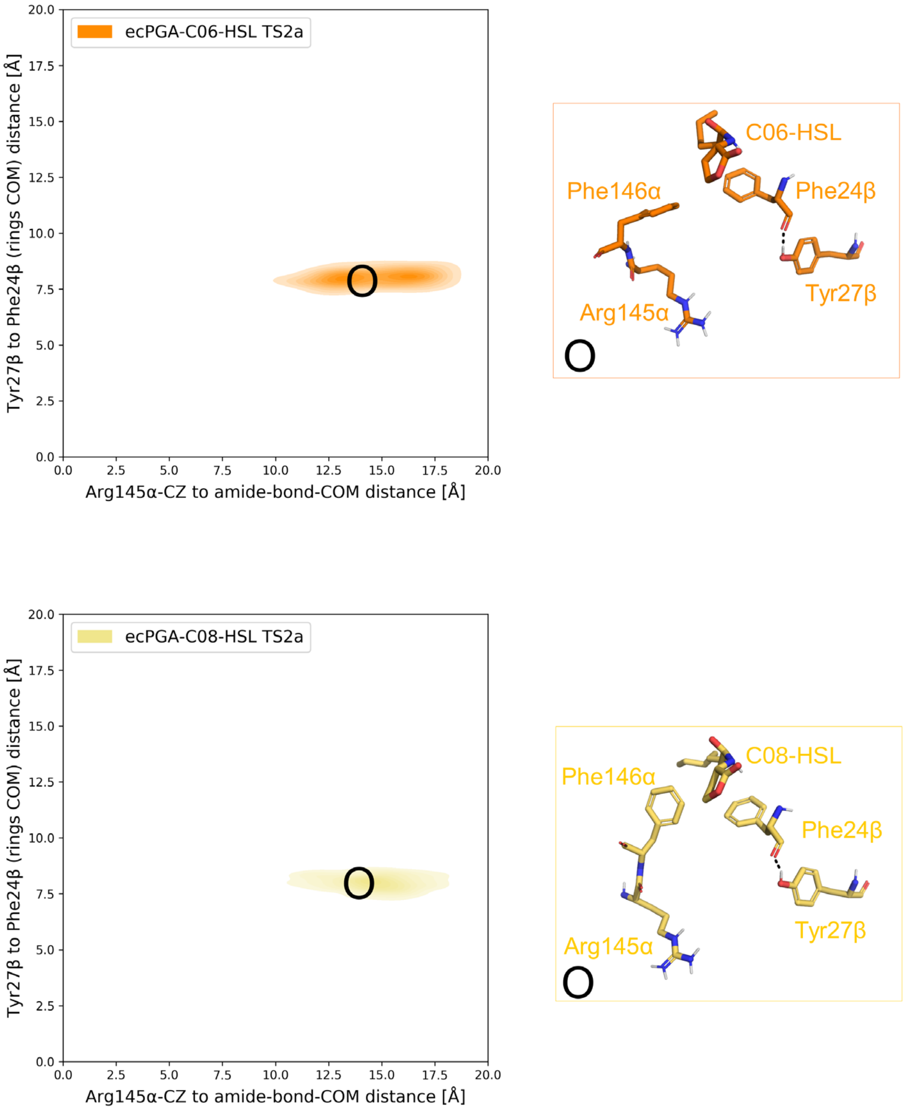

**Figure S39.** Dynamics of the gate controlling overall access to the active site of ecPGA. For both substrates, we observed that ecPGA preferred the open conformation defined by Arg145α guanidyl moiety being solvent-exposed instead of being near the backbone oxygen of Phe24β, which is primarily unavailable due to its hydrogen-bonding with Tyr27β. Such arrangement keeps the Arg145α side-chain far from the amide bond of the substrate, resulting in a lower energy barrier at TS2a than in aPGA-C08-HSL, which preferred the closed state.
